# Lytic bacteriophages facilitate antibiotic sensitization of *Enterococcus faecium*

**DOI:** 10.1101/2020.09.22.309401

**Authors:** Gregory S. Canfield, Anushila Chatterjee, Juliel Espinosa, Mihnea R. Mangalea, Emma K. Sheriff, Micah Keidan, Sara W. McBride, Bruce D. McCollister, Howard C. Hang, Breck A. Duerkop

## Abstract

*Enterococcus faecium*, a commensal of the human intestine, has emerged as a hospital-adapted, multi-drug resistant (MDR) pathogen. Bacteriophages (phages), natural predators of bacteria, have regained attention as therapeutics to stem the rise of MDR bacteria. Despite their potential to curtail MDR *E. faecium* infections, the molecular events governing *E. faecium-phage* interactions remain largely unknown. Such interactions are important to delineate because phage selective pressure imposed on *E. faecium* will undoubtedly result in phage resistance phenotypes that could threaten the efficacy of phage therapy. In an effort to understand the emergence of phage resistance in *E. faecium*, three newly isolated lytic phages were used to demonstrate that *E. faecium* phage resistance is conferred through an array of cell wall-associated molecules, including secreted antigen A (SagA), enterococcal polysaccharide antigen (Epa), wall teichoic acids, capsule, and an arginine-aspartate-aspartate (RDD) protein of unknown function. We find that capsule and Epa are important for robust phage adsorption and that phage resistance mutations in *sagA, epaR*, and *epaX* enhance *E. faecium* susceptibility to ceftriaxone, an antibiotic normally ineffective due to its low affinity for enterococcal penicillin binding proteins. Consistent with these findings, we provide evidence that phages potently synergize with cell wall (ceftriaxone and ampicillin) and membrane-acting (daptomycin) antimicrobials to slow or completely inhibit the growth of *E. faecium*. Our work demonstrates that the evolution of phage resistance comes with fitness defects resulting in drug sensitization and that lytic phages could potentially serve as antimicrobial adjuvants in treating *E. faecium* infections.

## Introduction

Enterococci are intestinal commensal bacteria and important opportunistic human pathogens (1). Of the two most clinically relevant enterococcal species, *Enterococcus faecalis* and *Enterococcus faecium*, the emergence of multidrug resistance is observed most commonly with *E. faecium* (2). Considering that effective antibiotics with activity against multidrug-resistant (MDR) *E. faecium* are limited, clinicians are often forced to use antibiotic combination therapy to treat these infections (3). Although this approach can be life-saving, these regimens increase the risk of patient adverse drug events, drug-drug interactions, dysbiosis, and may fail to cure the infection (4). Rising from desperate treatment dilemmas like these are several examples of the successful use of phage therapy to treat MDR bacterial infections in humans (5-8). These success-stories have motivated renewed interest in the use of phage therapy for treatment of bacterial infections. Despite this motivation, relatively little is understood about the bacterial receptors exploited by phages to infect their bacterial hosts and the counter-measures employed by bacteria to avoid phage infection. We believe that understanding the molecular events that lead to phage resistance in MDR bacteria may help mitigate the threat of phage therapy failure.

Recently, our group and others have begun to elucidate the molecular mechanisms that enable successful phage infection of enterococci and the bulk of these studies were performed for *E. faecalis* and its interactions with tailed dsDNA phages (8-15). The molecular mechanisms enabling phage infection in *E. faecium* are poorly understood. Our knowledge of potential *E. faecium* phage receptors comes from an *in vitro* study where the co-existence of phages and *E. faecium* was studied through multiple passages in laboratory media (12). Whole genome sequencing of phage resistant survivors showed mutations in the capsule tyrosine kinase *ywqD2* (equivalent to *wze*), RNA polymerase β-subunit (*rpoC*), several predicted hydrolases, and a cell wall precursor enzyme. It was proposed that these mutations conferred phage resistance, though direct genetic testing of this hypothesis was not performed. Tandem-duplications in a putative phage tail fiber gene (EFV12PHI1_98) supported evolution of phages that overcame adaptive changes that resulted in phage resistance of *E. faecium* (12).

In this work, we expand on our understanding of phage-enterococcal interactions by identifying genes important for lytic phage infection of clade B strains of *E. faecium.* We have isolated three previously uncharacterized *E. faecium-specific* phages and show that each belong to the *Siphoviridae* morphotype of the *Caudovirales* and resemble previously described lytic enterococcal phages (9-11, 14). Protein coding sequence comparison to other enterococcal phages reveals that one phage belongs to a novel enterococcal phage orthocluster and the remaining two phages belong to previously described enterococcal phage orthoclusters (16). To identify the molecular determinants of *E. faecium* phage infection, we used these three phages to generate a collection of *E. faecium* phage resistant mutants. Phage resistance mutations mapped to genes encoding the cell wall hydrolase secreted antigen A (*sagA*), putative teichoic acid precursors of the enterococcal polysaccharide antigen (*epa*), capsule biosynthesis enzymes, and an arginine-aspartate-aspartate (RDD) protein of unknown function. Capsule and putative teichoic acid biosynthesis proteins were shown to influence phage adsorption. Considering that all of the genes identified are involved in cell wall biochemistry and/or architecture, we determined if these phage resistance mutations result in fitness tradeoffs that lead to altered antimicrobial susceptibility. Phage resistant strains harboring mutations in *sagA, epaX*, and *epaR* showed enhanced susceptibility to cell wall and/or membrane-acting antibiotics, including ceftriaxone, ampicillin, and daptomycin. We discovered that combining phages with cell wall or membrane-acting antimicrobials acts synergistically to inhibit the growth of *E. faecium*. These findings suggest lytic phages might be leveraged as antibiotic adjuvants to offset the emergence of multi-drug resistant strains of *E. faecium* in hospitalized patients.

## Results

### Genome sequence analysis and morphology of novel lytic *E. faecium* bacteriophages

*E. faecium* phages 9181, 9183 and 9184 were isolated from raw sewage by plaque assay using *E. faecium* clade B strains Com12 and 1,141,733 (17). We chose to focus on clade B strains (commensal-associated) given reports that these strains can serve as a reservoir for transmission of multidrug resistance plasmids to clade A1 (hospital-associated) strains (18). Evaluation of phage morphology by TEM revealed that all three phages were non-contractile tailed phages characteristic of the *Siphoviridae* morphotype (Fig. 1) (19). DNA sequence analysis demonstrated that the phage 9181, 9183 and 9184 genomes are 71,854bp, 86,301bp, and 44,601bp in length, respectively (Fig. 1). The genomes of phages 9181 and 9183 were assembled into single contigs. The phage 9184 genome assembled into two contigs, with a 53-bp sequencing gap located near the 5’ end of a predicted BppU-family phage baseplate upper protein. In total, 123, 128, and 73 open reading frames (ORFs) were identified for phages 9181, 9183 and 9184, respectively (Table S1). Genome modularity based on predicted gene function was observed for each phage genome, however, for phage 9181 the lysin and holin genes are located at the 5’ and 3’ termini of the genome (Fig. 1). Functional classifications, consisting of replication or biosynthesis, DNA packaging, phage particle morphogenesis, nucleic acid restriction and modification, host cell lysis, sensory function, sugar transferase and a potential β-lactamase, could be predicted for approximately 30%, 47%, and 48% of the phage 9181, 9183, and 9184 ORFs, respectively (Table S1). The remaining genes were predicted to be hypothetical genes or genes containing domains of unknown function. A PCR screen for phage lysogeny in phage-resistant *E. faecium* mutants failed to identify phage 9181, 9183, and 9184 DNA integration within their respective *E. faecium* host genomes (Table 1; Fig. S1). These data are consistent with a lack of phage DNA among genomic reads from phage 9181, 9183, and 9184-resistant *E. faecium* mutants and the absence of turbid plaques, a feature often attributed to lysogenic phages. Together, these data indicate that phage 9181, 9183, and 9184 are most likely obligate lytic phages when preying on *E. faecium* Com12 or 1,141,733.

**Figure 1.**
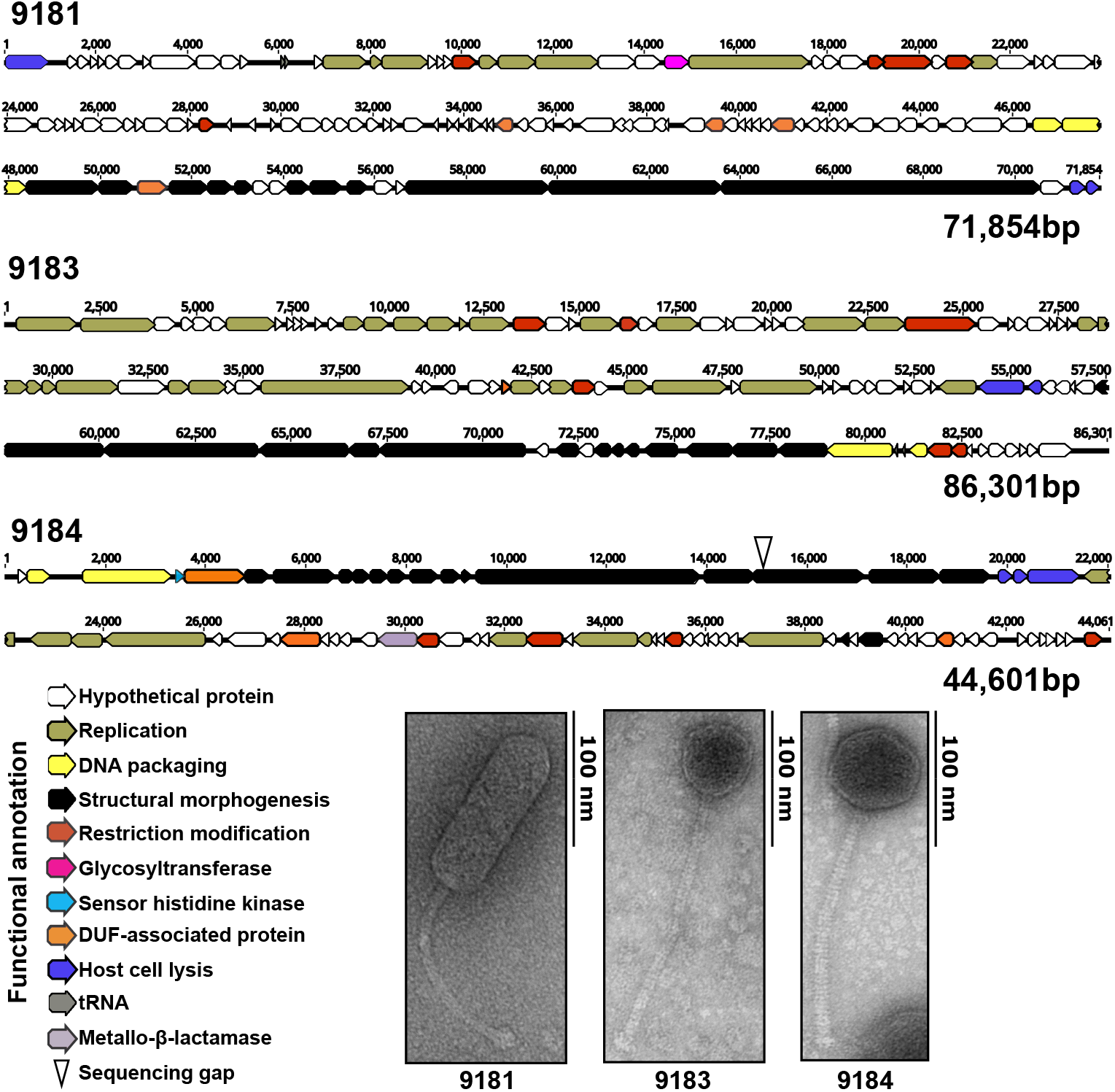
Genome organization and morphogenesis of three previously uncharacterized *E. faecium* phages. Whole genome sequencing reveals a modular functional organization of phage 9181, 9183 and 9184 genomes. Open reading frames for each phage were determined by RAST version 2.0 and by the Texas A&M Center for Phage Therapy structural analysis workflow version 2020.01. Colored open reading frames correspond to functional prediction. Beneath the phage genome maps, TEM shows phage 9181, 9183 and 9184 are non-contractile tailed *Siphoviridae*. The *E. faecium* host strain for phage 9181 is *E. faecium* Com12. The host strain for phage 9183 and 9184 is *E. faecium* 1,141,733.

**Table 1.**
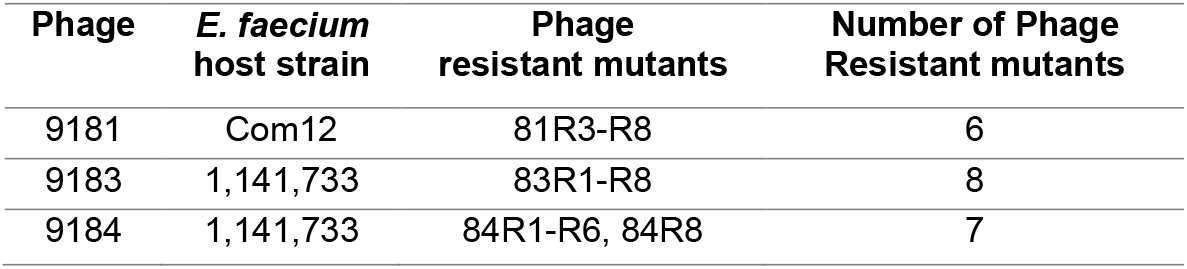
Phages, *E. faecium* host strain, and phage resistant mutants

| **Phage** | ***E. faecium***<br>**host strain** | **Phage**<br>**resistant mutants** | **Number of Phage**<br>**Resistant mutants** |
| --- | --- | --- | --- |
| 9181 | Com12 | 81R3-R8 | 6 |
| 9183 | 1,141,733 | 83R1-R8 | 8 |
| 9184 | 1,141,733 | 84R1-R6, 84R8 | 7 |

### Comparative genome analysis places phages 9181, 9183, and 9184 in distinct orthoclusters

Comparative genome analysis of phages 9181, 9183 and 9184 was performed with all publicly available enterococcal phage genomes using OrthoMCL, an algorithm that identifies clusters of orthologous proteins from at least two phages enabling phylogenetic categorization of phage proteins into orthoclusters (16, 20). Of the 10 enterococcal phage orthoclusters originally identified by Bolocan et al. (16), OrthoMCL clustering places phage 9184 into orthocluster I and phage 9183 into orthocluster X (Fig. 2). Phage 9181 forms a new orthocluster that we have named orthocluster XI (Fig. 2). Whole genome alignments of phages 9183 and 9184 to their nearest orthocluster neighbors, VPE25 and VFW for 9183 and vB_EfaS-DELF1 and IME-EFm5 for 9184, revealed conserved protein sequence identity and similar genome organization (Fig. S2A and S2B). Conversely, phage 9181 shared little protein sequence identity and genome organization to its nearest neighbors, phage EFC-1 and phage FL4A, supporting its placement as the sole member of a new orthocluster (Fig. S2C). Higher protein sequence identity and more similar genome organization was observed for phages belonging to the same orthocluster rather than phages belonging to different orthoclusters. Since the publication of Bolocan et al., an additional 45 phage genomes have been made publically available, resulting in the identification of a 12^th^ orthocluster consisting of phages EFA-1 and EFA-2, two recently described phages of unknown morphology (Fig. 2). Consistent with prior observations of orthocluster I phages, a β-lactamase domain-containing protein (ORF35) was found in the genome of phage 9184 (Fig. 1 and Table S1C). Similar to phages in orthocluster X, an integrase-family recombinase was found in the genome of phage 9183 (Table S1B). However, prior evidence demonstrates that other members of this orthocluster are unable to lysogenize their *E. faecalis* host (11), which is consistent with absence of lysogenized phage 9183 in phage 9183 resistant mutants as mentioned above (Fig. S1B).

**Figure 2.**
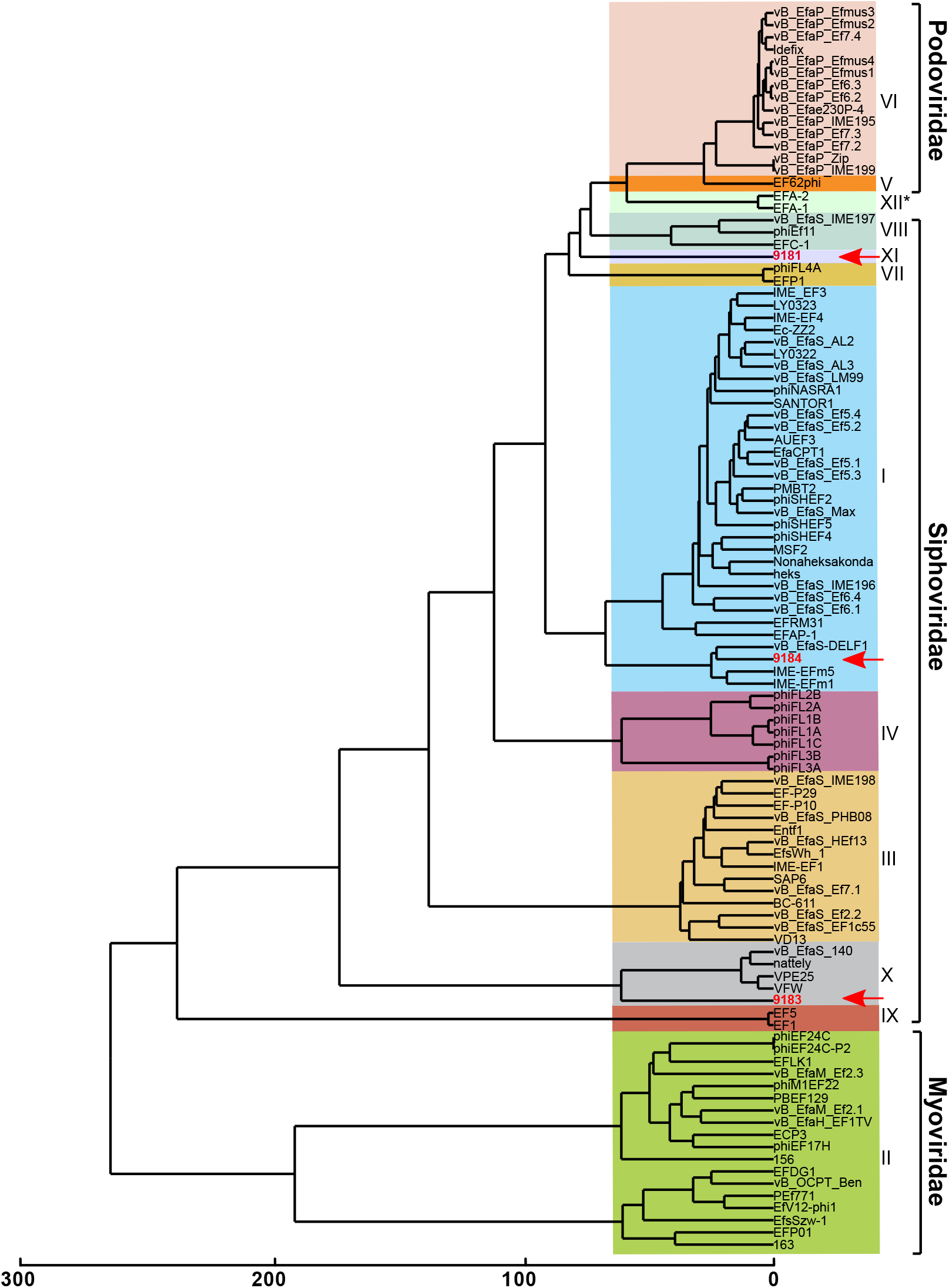
Comparative genomic analysis identifies two novel enterococcal phage orthoclusters. A comparative genome analysis was performed using OrthoMCL as described previously by Bolocan et al. (16). A phylogenetic proteomic tree was constructed from the OrthoMCL matrix using the Manhattan distance metric and hierarchical clustering using an average linkage with 1000 iterations. Ninety-nine enterococcal phage genomes available from NCBI were used for comparison to *E. faecium* phages 9181, 9183, and 9184 (highlighted in red, emphasized by red arrows). Distinct phage orthoclusters are represented by colored boxes. Roman numerals to the right of the shaded boxes signify the phage orthocluster number. Phage orthocluster morphology is indicated by calipers (if known) or an asterisk symbol (if unknown) to the right of the roman numerals.

### *E. faecium* phages have broad and narrow tropism for laboratory and clinical *E. faecium* isolates

We next sought to determine the host range of each phage against strains of *E. faecium* and *E. faecalis.* To achieve this, a phage susceptibility assay was performed by spotting 10-fold serially-diluted enterococcal cultures on Todd-Hewitt broth (THB) agar embedded with phages 9181, 9183 or 9184. A panel of 10 laboratory *E. faecium* isolates and 11 contemporary MDR clinical *E. faecium* isolates were selected for this analysis (Table S4) (17). An *E. faecium* strain was considered phage-susceptible if less than 1 x 10^5^ CFU/mL were recovered following phage exposure, representing greater than 4-log of bacterial killing. Phages 9181 and 9183 demonstrated narrow host ranges against laboratory *E. faecium* strains (Fig. 3A). Besides the host strain on which the phage was isolated (Com12 for phage 9181 and 1,141,733 for phage 9183), only *E. faecium* Com15 was susceptible to phage 9181, while no other *E. faecium* laboratory strain tested was susceptible to phage 9183. Contrarily, 60% of the laboratory *E. faecium* strains were susceptible to phage 9184, including clade A and B strains (Fig. 3A). There was an absence of susceptibility to phage 9181 and 9183, and reduced susceptibility (~36%) to phage 9184 for the contemporary MDR clinical *E. faecium* isolates (Fig. 3B). Efficiency of plaquing assay revealed that phages 9181 and 9184 most efficiently plaqued on their respective host strains (Fig. 3C-D). Together these data show that phage 9184 has a broader host range compared to phages 9181 and 9183 and that these phages plaque most efficiently on their *E. faecium* host strains, a likely byproduct of repeated phage propagation on the same strain (21). Interestingly, *E. faecium* 1,231,501 and 1,230,933, the latter of which is a multi-drug resistant clade A strain, lacked susceptibility to phage 9181, 9183 and 9184. None of the three phages were capable of infecting any of the 10 clinical *E. faecalis* strains tested (designated UCH12-20 in Table S4), suggesting that these phages are specific for *E. faecium*.

**Figure 3.**
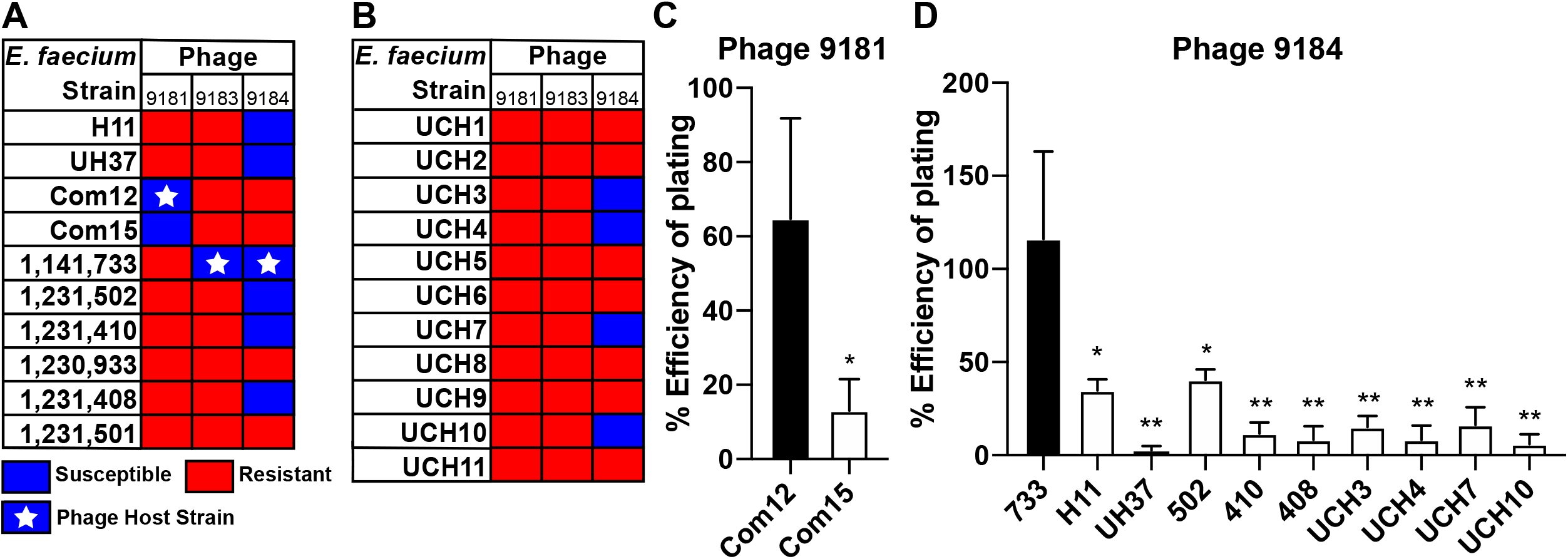
*E. faecium* phages demonstrate broad and narrow host ranges and plaque most efficiently on their laboratory host strain. Host ranges of phage 9181, 9183, and 9184. Phage 9181 and 9183 have a narrow *E. faecium* host range, while phage 9184 shows a broader host range. Bacteria were susceptible if less than 1 x 10^5^ CFU/mL of bacteria were recovered from a phage susceptibility assay. Bacteria were resistant if greater than 1 x 10^5^ CFU/mL of bacteria were recovered from a phage susceptibility assay. (A) Indicates host range for a collection of laboratory strains. (B) Indicates the host range for a collection of clinical isolates provided by the clinical microbiology lab at the University of Colorado, Anschutz Medical Campus. A white star signifies the *E. faecium* host strain utilized for phage propagation. Efficiency of plating assay shows that phage 9181 (C) and 9184 (D) plaque most efficiently on their laboratory host strains. Data represent the average of three replicates ± the standard deviation. *, *P*< 0.05 and **, *P* < 0.01 by unpaired Student’s t-test.

### Phage predation elicits spontaneous and stable phage resistance in *E. faecium*

To identify *E. faecium* genes that are involved in phage infection, we isolated spontaneous phage-resistant *E. faecium* strains following exposure to phages 9181, 9183 and 9184. Phage-resistant isolates were identified by plating stationary phase cultures of *E. faecium* Com12 and 1,141,733 on THB agar embedded with phages 9181, 9183, or 9184. Colonies that arose on these plates represented potential phage-resistant colonies. To confirm the stability of the phage-resistant phenotype, a colony was serially passaged daily for 3 days on THB agar before re-streaking again on phage embedded THB agar. The growth of a strain in the presence of phage following serial passage suggested a stable phage-resistant phenotype (Fig. 4A-C). Six to eight independent phage-resistant strains were further characterized for phages 9181, 9183 and 9183 (Table 1 and Table S2). For phages 9181 and 9183 resistant *E. faecium* strains (denoted 81R3-8 and 83R1-8, respectively) we observed bacterial growth in the presence of phages to levels that were similar to bacterial growth in the absence of phages indicating a strong resistance phenotype (Fig. 4A, 4B and Fig. S3A, S3B, S3D, S3E). However, for phage 9184 we observed limited phage resistance in all but one presumed *E. faecium* phage resistant isolate (Fig. 4C and Fig. S3C, S3F) suggesting that robust resistance to phage 9184 may be multifactorial.

**Figure 4.**
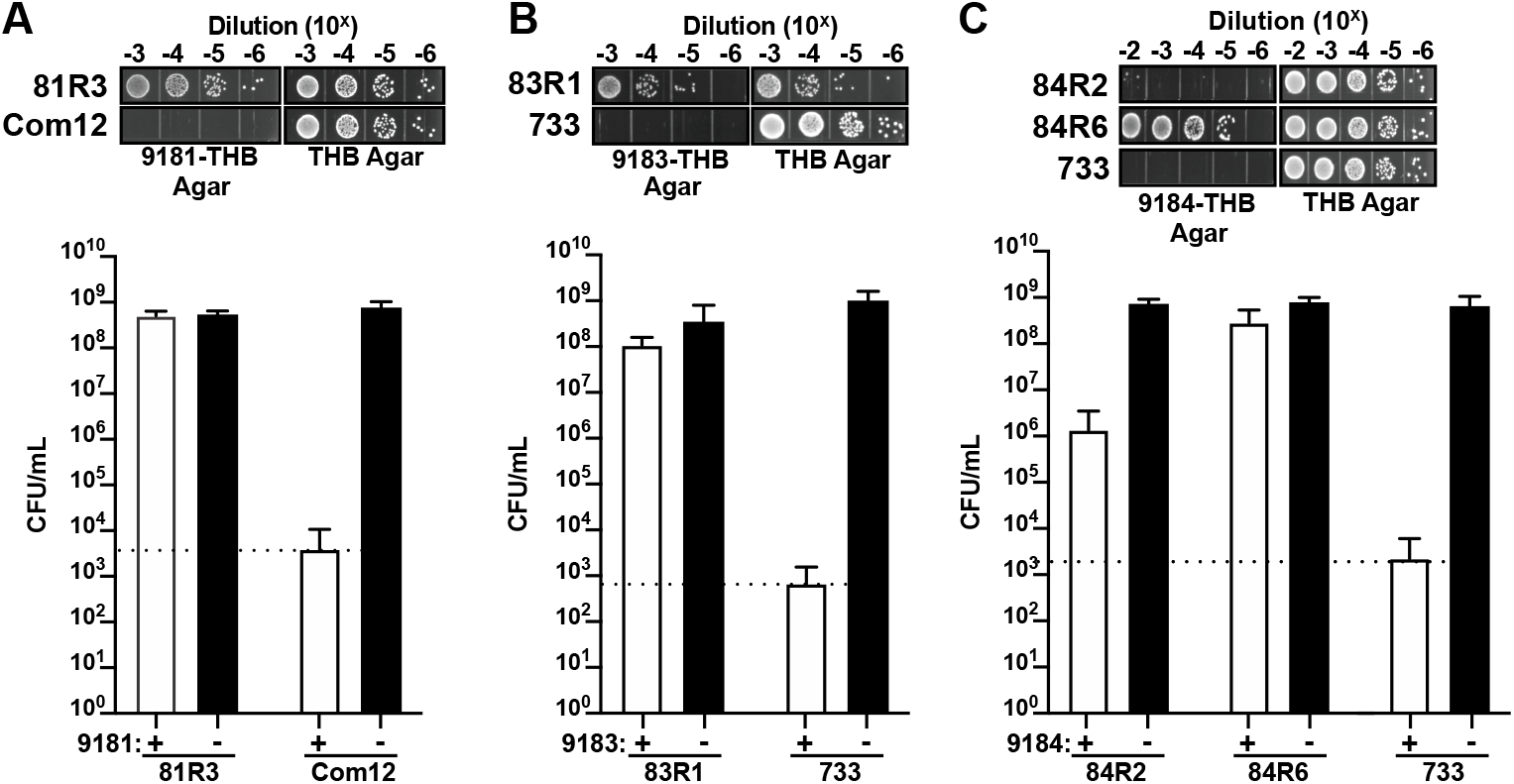
*E. faecium* elicits a robust resistance phenotype to phage 9181 and 9183, but variable resistance to phage 9184. Representative phage resistant strains raised against phages 9181 (A), 9183 (B), and 9184 (C). Data show phage susceptibility assays and associated bacterial enumeration of wild type and phage resistant mutants in the presence (white bars) or absence (black bars) of phage from three independent experiments. Error bars indicate standard deviation. Phage 9181 resistant (A) and phage 9183 resistant (B) strains exhibit ≥ 4-log of survival in the presence of phages compared to the parental *E. faecium* Com12 and 1,141,733 (733) strains, respectively. Phage 9184 resistant strains (C) exhibit diverse resistance strength characterized by weak (84R2) and strong (84R6) resistance phenotypes. The dotted line indicates the spontaneous mutation threshold of wild type *E. faecium*, which is defined as the mean CFU per ml at which spontaneous phage resistance is observed for the wild type host strain of each phage.

### *E. faecium* phage resistance mutations occur in cell wall biosynthesis and architecture genes and a gene encoding a transmembrane protein

To identify genetic changes conferring a phage resistance phenotype, we performed whole genome DNA sequencing of phage resistant and parental *E. faecium* strains. We observed unique and conserved genome mutations in strains that had developed phage resistance (Fig. 5A-D and Table S2A-C).

**Figure 5.**
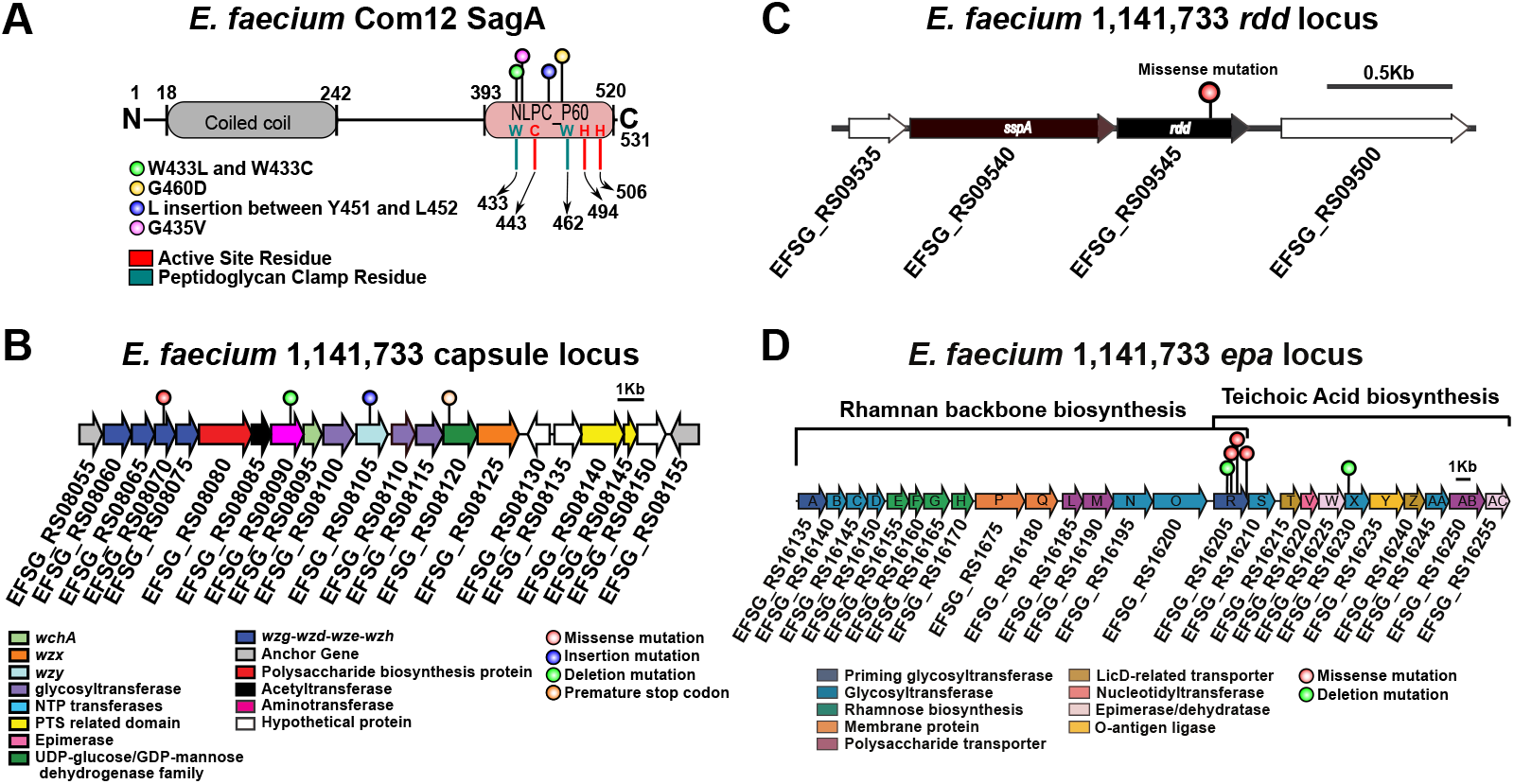
A diverse assortment of mutations confers phage resistance in *E. faecium*. (A) Protein secondary structure of *E. faecium* Com12 SagA, consisting of an N-terminus coiled-coil domain (residues 18-242) and C-terminus NlpC_P60 peptidoglycan hydrolase domain (residues 393-520). Displayed above the protein structure are colored lollipops denoting the site of mutations within NlpC_P60 domain of phage 9181-resistant mutants. Inside and below the protein structure are colored one letter amino acid abbreviations and lines, respectively, corresponding to key active site (red) and peptidoglycan clamp residues (teal) of the NlpC_P60 domain. Abbreviations: W, tryptophan; C, Cysteine; H, Histidine; G, Glycine; D, Aspartate; L, Leucine; Y, Tyrosine; V, Valine. (B) Capsule locus mutations are detected in a tyrosine kinase (*wze*), aminotransferase (*efsg_rs08090*), *wzy* (*efsg_rs08105*), and nucleotide sugar dehydrogenase (*efsg_rs08120*) of phage 9184-resistant mutants. Arrows indicate open reading frames. Arrow colors correspond to colored boxes (figure bottom left) indicate predicted open reading frame function (17). Colored lollipops above the arrows corresponding to colored dots (figure bottom right) indicate the mutational type. *E. faecium* 1,141,733 locus tags are angled below the arrows. (C) A missense mutation is found within a predicted arginine-aspartate-aspartate protein (*rdd*; black arrow) of one phage 9184-resistant mutant (84R6) of *E. faecium* 1,141,733. *rdd* is flanked upstream by a predicted hypothetical protein (white arrow) and signal sequence peptidase A (*sspA*; black arrow) and downstream by another hypothetical protein (white arrow). *E. faecium* 1,141,733 locus tags are angled below the arrows. (D) Mutations in predicted teichoic acid biosynthesis genes (*epaR* and *epaX*) are identified in phage 9183-resistant mutants of *E. faecium* 1,141,733 (25). Arrow colors correspond to colored boxes (figure bottom left) indicate predicted open reading frame function. Colored lollipops above the arrows corresponding to colored dots (figure bottom right) indicate the mutational type. *E. faecium* 1,141,733 locus tags are angled below the arrows. The brackets above the locus correspond the conserved (left) and variable (right) portions of the *epa* locus proposed to by Gueredal et al. to encode the machinery necessary for rhamnopolysaccharide synthesis and wall teichoic acid biosynthesis, respectively (25).

Five of six mutations identified in phage 9181-resistant strains were detected in *efvg_rs16270*, which in the *E. faecium* Com12 reference genome is annotated as a hypothetical protein and was flanked by a 5’ sequencing gap. Closure of this sequencing gap by PCR and amplicon sequencing revealed that *efvg_rs16270* encodes the *E. faecium* secreted antigen A (SagA) protein. Whole genome sequencing showed that all *sagA* mutations localized at or near the peptidoglycan clamp or active site residues of the NlpC_P60 hydrolase domain of SagA, which was recently shown to function as an endopeptidase that cleaves crosslinked Lys-type peptidoglycan fragments (Fig. 5A and Table S2A) (22). To determine the impact of *sagA* mutations on protein structure and function, each single nucleotide polymorphism-associated *sagA* mutant was assessed by Missense 3D analysis (23). BLASTp alignment of SagA from *E. faecium* Com12 and Com15 showed 95% identity along the entire length of the protein and *E. faecium* Com12 and Com15 exhibit identical protein homology in the NlpC_P60 hydrolase domain (Fig. S4A), suggesting that SagA should be functionally conserved between these two stains. Therefore, we used the *E. faecium* Com15 NlpC_P60 crystal structure (PDB 6B8C) in Missense 3D to assess the impact of residue changes on the structure and function of NlpC_P60 hydrolase in our *sagA* mutant strains (22). Except for one SagA mutant (81R8; G435V), no structural damaging mutations were found. Using the supernatants and cell pellets of exponentially growing (OD_600_ ~0.8) wild type and *sagA* mutants, we performed Western blots for SagA expression. All *sagA* mutants produced similar levels of both intracellular and secreted SagA suggesting that these *sagA* mutants are likely catalytically inactive or dampened because of mutations in the NlpC_P60 hydrolase domain (Fig. S4B, S4C, and S4D). We then complemented the *sagA* mutations in phage 9181 resistant strains using a construct previously generated, pAM401-*sagA*, which carries the *sagA* gene and its native promoter from *E. faecium* Com15 (24). For all *sagA* mutants, complementation with pAM401-*sagA* restored phage susceptibility (Fig. S5A). These results suggest that SagA hydrolase activity may be dispensable for *E. faecium* viability and that non-crosslinked peptidoglycan in *E. faecium* Com12 is important for phage 9181 infection.

One phage 9181-resistant strain (81R7) harbored mutations in capsule tyrosine kinase (*wze*) and topoisomerase III (*topB*) genes and lacked a *sagA* mutation (Table S2A). Similarly, sequencing analysis of all 9184 resistant strains (84R1-6) revealed an assortment of mutations in the capsule biosynthesis locus. Nonsense, insertion and deletion mutations were detected in *wze*, capsule aminotransferase (*efsg_rs08090*), capsule polymerase (*wzy*), and capsule nucleotide sugar dehydrogenase (*efsg_rs08120*) genes (Fig. 5B and Table S2C). Prior co-evolution experiments between the *Myoviridae* phage 1 and *E. faecium* TX1330 revealed a propensity for *wze* mutations within an evolved phage resistant *E. faecium* population (12). Our data is consistent with this observation and suggests that *E. faecium* capsule might serve as a possible receptor and/or adsorption factor for phage 9181 and 9184. We found that complementation of certain capsule mutants, using the constitutive expression vector pLZ12A (i.e. 84R2 with *efsg_rs08120* and 84R5 with *efsg_rs08090*), partially restored phage 9184 susceptibility, while complementation of other capsule mutants (i.e. 84R6 and 81R7 each with *wze*) failed to restore phage susceptibility (Fig. S5B-C). This result suggests that capsule is not a major factor mediating phage resistance to phage 9181 (Fig. S5B) and only weakly promotes phage 9184 resistance when select capsule genes are mutated (Fig. S5C). These results emphasize the importance of other non-capsule associated mutations in conferring phage-resistance to phage 9181 (*sagA*) and phage 9184 (*rdd*). We attempted to address the non-capsule associated mutation in strain 81R7 (*topB*) and its involvement in phage 9181 resistance, however, all attempts to clone *topB* into the pLZ12A resulted in truncated *topB* inserts following transformation into *Escherichia coli*, suggesting that constitutive expression of *E. faecium topB* may be toxic to *E. coli*. Similarly, to address the role of the non-capsule mutation detected in 84R6, which exhibited a robust phage 9184-resistance phenotype, a predicted arginine-aspartate-aspartate gene (*rdd*), this gene was successfully cloned into pLZ12A yet transformation of this construct into *E. faecium* 84R6 was unsuccessful despite repeated attempts. Given the ease with which *pLZ12A-wze* and empty pLZ12A vector were transformed into *E. faecium* 84R6 and our repeated failure to successfully recover transformants harboring pLZ12A-*rdd* suggests that over-expression of *rdd* in *E. faecium* 84R6 may be lethal.

Analysis of *E. faecium* phage 9183 resistant strains (83R1-8) identified mutations in *epa* genes, *epaR* and *epaX* (Fig. 5D and Table S2B). Mutation of *epaR* and *epaX* results in *E. faecalis* phage resistance (9, 10, 14) and recently it was determined that the *epaR* and *epaX* genes of *E. faecalis* V583 participate in wall teichoic acid biosynthesis (25). Considering that mutation of the *epaX* homologs *epaOX* and *epaOX2* from *E. faecalis* OG1RF conferred phage VPE25-resistance by limiting phage adsorption (10, 11), we suspect that teichoic acids also mediate adsorption of phage 9183 to *E. faecium* 1,141,733. We were surprised that we did not find any phage 9183 resistant strains with mutations in PIP_EF_, given the high protein homology and similar genome organization observed between phages 9183, VPE25 and VFW, the latter two which use PIP_EF_ as a receptor (11) (Fig. 2 and Fig. S2A). To confirm that mutations in the *epa* locus confer phage resistance in *E. faecium*, we pursued a similar complementation strategy as above with the *epaR* and *epaX* mutants identified in the phage 9183-resistant mutants. All phage 9183-resistant mutants complemented with either the *epaR* or *epaX* were restored for phage susceptibility (Fig. S5D). Given the importance of D-alanylation in teichoic acid biosynthesis, we performed complementation with pLZ12A-*dltA* in the *epaX* and *dltA* double mutant (83R7). We observed that only pLZ12A-*epaX*, not pLZ12A-*dltA*, was capable of restoring phage susceptibility in 83R7 (Fig. S5D). Considering that EpaX acts upstream of DltA in the biosynthesis of teichoic acids (25), these data suggest that *dltA* is dispensable during phage infection, lending further support to the notion that the *epa* variable locus involved in teichoic acid biosynthesis is a driver of *E. faecium* infection by phage 9183.

### *E. faecium* phage resistant mutants have phage adsorption defects

To determine if phage adsorption defects occur due to phage resistance, we sought to quantify phage 9181, 9183, and 9184 adsorption to wild type and phage resistant *E. faecium* strains using a phage adsorption assay (9, 10, 14). For phage 9181 resistant strains, we observed no significant change in percentage adsorption to *sagA* mutant strain 81R5, nor phage resistant strain 81R7 harboring a *wze* and *topB* mutations (Fig. 6A). These results are consistent with the inability of *wze* complementation to enhance phage 9181 adsorption in 81R7 (Fig. S6A). This result suggests that mutation of *wze* in *E. faecium* Com12 has little to no effect on phage 9181 adsorption. Given that SagA is expressed into supernatants of phage 9181 resistant *sagA* mutants, it remains possible that phage 9181 adsorbs to SagA or non-crosslinked peptidoglycan at the surface of *E. faecium* Com12. Furthermore, it is possible that complementation of *topB* in the 81R7 background might cause transcriptional or translational changes in surface expressed molecules enabling enhanced phage 9181 adsorption phenotype (Fig. 6A). Unfortunately, the lack of a *sagA* knock-out mutant and *topB* complementation vector prevented us from addressing these questions.

**Figure 6.**
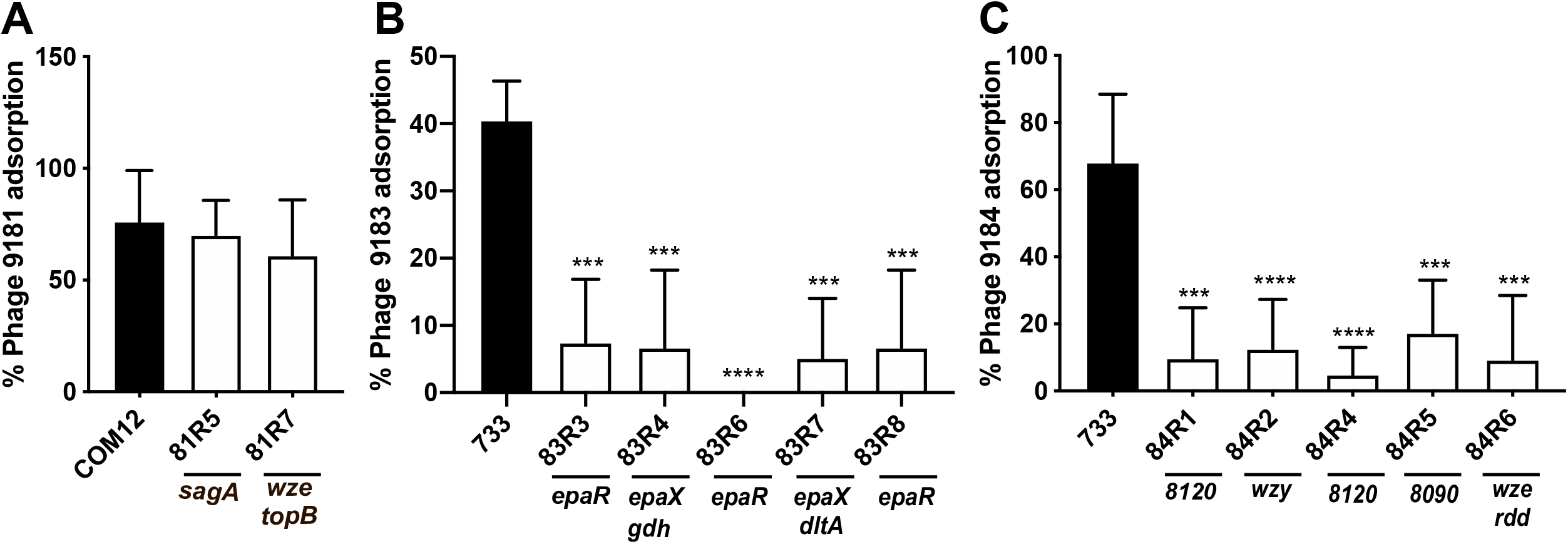
Mutation in the capsule and exopolysaccharide loci limit phage adsorption in *E. faecium*. Shown is the percentage phage adsorption in phage 9181 (A), 9183 (B), and 9184 (C) compared to parental strains, *E. faecium* Com12 and *E. faecium* 1,141,733, respectively. Results represent average percent adsorption and standard deviation from three independent experiments. ***, *P* < 0.001; ****, *P* < 0.0001 by unpaired Student’s *t* test.

Previous work has demonstrated that *epa* mutants exhibit phage adsorption defects in *E. faecalis* (9, 10, 13, 14). Since we observed *epa* mutations that conferred phage 9183 resistance, we sought to determine if *epa* mutations might promote a similar phenotype in *E. faecium*. We observed a reduction in phage 9183 adsorption to mutants possessing *epaR* (83R6 and 83R8) and *epaX* (83R4 and 83R7) mutations compared with the parental strain (Fig. 6B). Although *epaX* mutants 83R4 and 83R7 were noted to also have mutations in *gdh* and *dltA*, respectively, we suspect that EpaX was the driver of this phenotype because of the known role of *epaX* homolog mutations to inhibit phage VPE25 adsorption to *E. faecalis* and that EpaX functions upstream of DltA in the biosynthesis of teichoic acid (10, 25). Consistent with this notion, we observed that over-expressing *epaR* and *epaX* in phage 9183 resistant mutants, 83R3 and 83R4, harboring mutations in *epaR* and *epaX* regained the ability to adsorb phage 9183 (Fig. S6B). Taken together, these results suggest that mutations in the *epa* locus of *E. faecium* lessen phage 9183 adsorption to the surface of its host strain.

To determine if mutations in the capsule locus facilitated phage 9184 adsorption defects, we performed phage 9184 adsorption assays using wild type and phage 9184 resistant mutants. We observed significant deficits in phage adsorption to strains harboring mutations in capsule polymerase (*wzy*; 84R1), nucleotide sugar dehydrogenase (*efsg_rs08120*; 84R4), aminotransferase (*efsg_rs08090*; 84R5) and tyrosine kinase (*wze*; 84R6) in comparison to the parental strain (Fig. 6C).

Complementation of 84R2 with *efsg_rs08120* restored phage 9184 adsorption, while complementation of 84R6 with *wze* partially restored phage 9184 absorption, albeit the change was statistically non-significant compared to the empty vector control strain (Fig. S6C). Given that 84R6 also harbors an *rdd* mutation which encodes a putative transmembrane protein, we cannot definitively conclude that the adsorption deficit was related to the *wze* mutation, as this *wze* mutation did not cause an adsorption defect for phage 9181 (Fig. 6A and Fig. S6A). Considering the adsorption defect is greater for the capsule mutants raised against phages 9184 compared to the phage 9181 capsule mutant 81R7, it is possible that additional surface associated molecules mediate the attachment of phage 9181 to *E. faecium* cells. Together, these data indicate that *E. faecium* capsule contributes to phage 9184 adsorption and may be phage specific.

### *E. faecium* phage resistance enhances β-lactam and lipopeptide susceptibility

With renewed interest focused on utilizing lytic phages for the treatment of bacterial infections and the observation that phage resistance can be a fitness tradeoff under antibiotic pressure (26, 27), we sought to determine the impact of *E. faecium* phage resistance on antimicrobial susceptibility. We performed antimicrobial susceptibility screening using E-test strips for the phage 9181, 9183, and 9184 resistant mutants compared to their parental strains to determine if phage resistance altered *E. faecium* antimicrobial susceptibility. For phage 9181 resistant mutants, we observed a ~2-5 fold reduction in the minimum inhibitory concentration (MIC) of ampicillin and an overall reduction in the MIC of ceftriaxone (Table S3A). Interestingly, the enhancement of ampicillin and ceftriaxone susceptibility correlated with phage 9181 resistant mutants harboring mutations in *sagA*, and not *wze* or *topB*. For phage 9183 resistant mutants, we also observed a 3-5 fold reduction in the MIC of ampicillin and an overall reduction in the MIC of ceftriaxone (Table S3B). Additionally, we noted a 2.5-5 fold reduction in the MIC of daptomycin, a lipopeptide class antimicrobial, which was not observed for the phage 9181 or 9184 resistant mutants. These results suggest that the acquisition of phage resistance via mutation of *sagA* and *epa* genes in *E. faecium* is a fitness defect that manifests as enhanced β-lactam susceptibility.

No phage capsule mutants showed a significant difference in antimicrobial susceptibility to β-lactams or lipopeptides, suggesting that mutations to the *E. faecium* capsule locus and *rdd* avoid the cost of increased antimicrobial susceptibility to β-lactams and daptomycin (Table S3C).

### Lytic phages synergize with β-lactam and lipopeptide antimicrobials to inhibit the growth of *E. faecium*

Considering the antibiotic fitness cost associated with phage resistance in *E. faecium*, we hypothesized that phages 9181 and 9183 would be capable of synergizing with ampicillin, ceftriaxone, and daptomycin to inhibit the growth of *E. faecium*. To address this question, we performed phage-antibiotic synergy assays where *E. faecium* was grown in the presence of phages alone, sub-inhibitory concentrations of ampicillin, ceftriaxone, or daptomycin alone, or a combination of phage and a sub-inhibitory concentration of antibiotics (Fig. 7A-E). For all three antibiotics, we observed that the combination of phage and sub-inhibitory concentrations of antibiotics were able to inhibit the growth of *E. faecium* better than phage or antibiotic alone. Given the absence of growth inhibition of *E. faecium* in the presence of sub-inhibitory concentrations of antibiotics alone, this result is consistent with a synergistic antimicrobial interaction between phages and antibiotics. Interestingly, the synergy observed between phages 9181 and 9183 and ceftriaxone appeared more potent than the synergy observed between these phages and ampicillin (Fig. 7A-D). A dose-response relationship emerged when ampicillin was combined with phages 9181 and 9183 where decreasing concentrations of ampicillin enabled varying degrees of bacterial population recovery (Fig. 7A-B). These data suggest that phages 9181 and 9183 could serve as useful adjuvants in combination with β-lactams for the treatment of *E. faecium* infections by restoring the susceptibility to *E. faecium* strains harboring intrinsic β-lactam resistance. We also observed that the combination of phage 9183 and daptomycin slowed the growth of *E. faecium* 1,141,733 more than phage 9183 alone or daptomycin alone (Fig. 7E). This suggests that phage 9183 also synergizes with daptomycin to inhibit *E. faecium* 1,141,733. These results are consistent with those observed by Morrisette et al. who observed synergy between the *Myoviridae* phage 113 and β-lactam (ampicillin, ertapenem and ceftaroline) and lipopeptide antimicrobials against daptomycin-resistant and tolerant strains of *E. faecium* (28).

**Figure 7.**
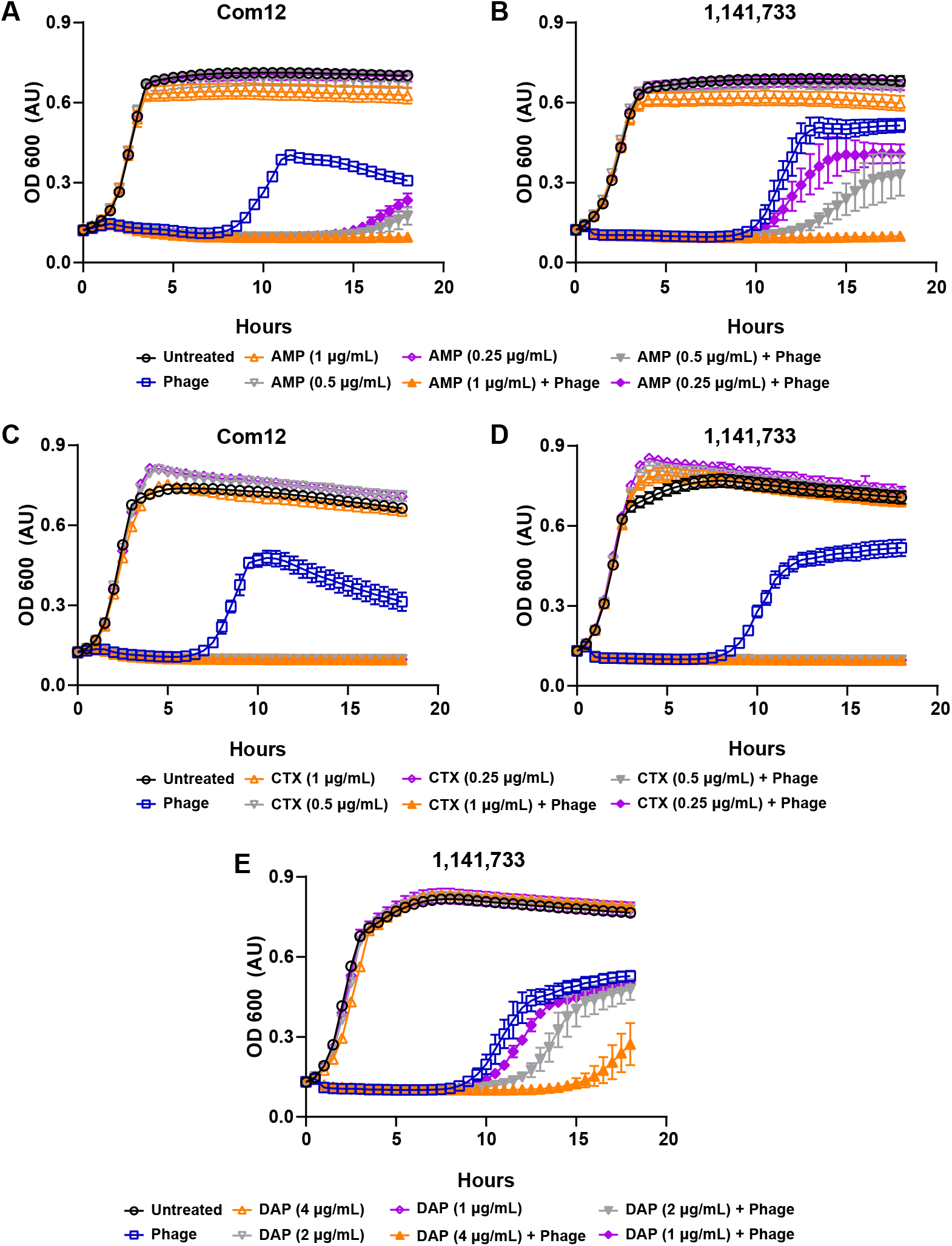
Phage 9181 and phage 9183 synergize with antibiotics to inhibit the growth of *E. faecium*. (A-E) *E. faecium* growth was monitored over 18 hours in the presence of phage (open blue squares), sub-inhibitory concentrations of antibiotics (open orange, grey, purple triangles or diamonds), both phage and sub-inhibitory concentration of antibiotics (filled orange, grey and purple triangles or diamonds), or media alone (open black circles). Phage 9181 was used in experiments with *E. faecium* Com12, while phage 9183 was employed for experiments with 1,141,733. Phages 9181 (A) and 9183 (B) synergize with sub-inhibitory concentration of ampicillin (AMP) in a dose responsive manner to slow the growth of *E. faecium* Com12 and 1,141,733, respectively. Phage 9181 (C) and 9183 (D) synergize with sub-inhibitory concentrations of ceftriaxone (CTX) to inhibit the growth of *E. faecium* Com12 and 1,141,733, respectively. Phage 9183 (E) synergizes with sub-inhibitory concentrations of daptomycin (DAP) in a dose-responsive manner to inhibit *E. faecium* 1,141,733. Three technical replicates were performed for each condition tested and the averages plotted. Error bars indicate standard deviation. Shown are the results from one experiment that was replicated in triplicate.

## Discussion

Considering the treatment pitfalls due to worsening drug resistance in *E. faecium* and other bacterial pathogens, the biomedical community is revisiting the use of phage therapy. Since phage therapy’s departure from 20^th^ century Western Medicine, new technologies have emerged that have facilitated fine-scale resolution of phage-bacterial molecular interactions. Despite these advancements, for many bacteria, including *E. faecium*, the molecular factors exploited by phages for infection remain largely understudied (12). We believe that studying the molecular interactions of phages with their *E. faecium* hosts will inform rational approaches for future phage therapies against this pathogen.

In this work, we describe three novel lytic phages of *E. faecium*. Using protein coding orthology, we show that one of these phages, phage 9181, forms a new orthocluster from the ten previously described enterococcal phage orthoclusters (16). We show that these phages are specific for *E. faecium* and exhibit broad and narrow strain tropism. Using whole genome sequencing and comparative genomics, we provide evidence that *sagA, epa*, and capsule biosynthesis genes are important for phage infection of *E. faecium*. We were unable to fully assess if the genes *topB* and *rdd* are important in conferring phage 9181 and phage 9184 resistance, respectively. We suspect that these genes aid in phage-*E. faecium* interactions. Consistent with previous observations in *E. faecalis* (9, 10, 13, 14), we show that mutations in *epaR* and *epaX* limit phage 9183 adsorption to *E. faecium*, albeit to a lesser extent than that observed for similar mutations in *E. faecalis*. For example, the difference in phage VPE25 adsorption to wild type *E. faecalis* versus an *epaOX* mutant was ~80% (10), compared with the ~35% reduction in phage 9183 adsorption to an *epaX* (*epaOX* homolog) mutant (Fig. 6B). This weaker phage 9183 adsorption despite the inability to infect the host closely resembles the ~50% reduction in phage SHEF2 adherence to an *epaB* and OPDV_11720 (encoding an *epaX*-like glycosyltransferase) mutants compared to wild type *E. faecalis* (29, 30). The partial adsorption of phage 9183 to *epaR* and *epaX* mutants despite phage resistance suggests that phage 9183 adherence might also depend on the core Epa rhamnopolysaccharide, in addition to Epa teichoic acid decorations, similar to phage SHEF2 (29). We show for the first time that mutations in the capsule locus, which is absent in *E. faecalis* (17), limits phage 9184 adsorption to *E. faecium* 1,141,733, but not phage 9181 adsorption to *E. faecium* Com12. The ~50% reduction in phage 9184 adsorption in capsule mutants (Fig. 6C), is comparable to the ~60-80% reduction of phages Ycsa and 8 against acapsular *Streptococcus thermophilus* (31).

Our investigation into fitness tradeoffs associated with *E. faecium* phage resistance revealed enhanced susceptibility to cell wall and membrane-acting antibiotics. We demonstrated that phages 9181 and 9183 synergize with cell wall and membrane-targeting antibiotics to more potently inhibit *E. faecium*. Importantly, this analysis revealed that phages 9181 and 9183 could sensitize *E. faecium* to ceftriaxone, an antibiotic that normally promotes enterococcal colonization of the intestine due to intrinsic resistance (32). Phage synergy with ceftriaxone is an important discovery as it suggests a strategy to re-sensitize enterococci to a third-generation cephalosporin. Exposure to cell wall-acting agents is recognized as a key event prior to hospital-acquired enterococcal infection in susceptible patients and cephalosporin re-sensitization could have a broad impact on anti-enterococcal therapy (2, 33). Cephalosporin activity pressures the native intestinal microbiota altering its ecology and related mucosal immunity, creating a scenario for enterococci to thrive and become dominant members of the microbiota (33-36). In patients with weakened immune systems or made vulnerable from hospital procedures such as surgeries, bone marrow ablative chemotherapy, or pre-existing alcoholic hepatitis/cirrhosis, these ceftriaxone-associated conditions can tip the scale in favor of infection (33-35, 37, 38). Even in *E. faecium* strains with ampicillin susceptibility, synergy with ceftriaxone for the treatment of endocarditis was demonstrated to be not absolute, suggesting that current Infectious Disease Society of America guidelines for the treatment of *E. faecium* endocarditis may lead to sub-optimal results (39, 40). Combination therapy with phage and cell wall or membrane-acting antimicrobials may offer a potential solution to circumvent this issue, while avoiding the risk associated with exposing patients to combination β-lactam agents.

The underlying molecular mechanisms conferring enhanced susceptibility to beta-lactams in *sagA, epaR* and *epaX* mutants remain unclear. Given that intrinsic resistance of *E. faecium* to ceftriaxone is derived, in part, from class A and B penicillin binding proteins (Pbps) (41-44), we hypothesize that modification of the surface architectural display of Pbps in *epaR, epaX*, and *sagA* mutants might facilitate this phenotype. Parallels to *E. faecium sagA* mutants can be drawn from mutation of a secreted peptidoglycan hydrolase in *E. faecalis*, SalB, which also demonstrates enhanced susceptibility to cephalosporins (45). Pairwise amino acid alignment of *E. faecium* Com12 SagA and *E. faecalis* SalB revealed 51% identity over the N-terminal coiled-coil domain region, which is expected given their different C-terminal hydrolase domains (SCP in SalB; NlpC_P60 in SagA). Contrary to *sagA* in *E. faecium, salB* was shown to be non-essential in *E. faecalis*, and has a homolog (*salA*) which may partly compensate for the function of *salB* to maintain cell viability (45). Staining of an *E. faecalis salB* mutant with a non-specific, fluorescent penicillin (Bocillin FL) revealed no difference from wild type. However, this analysis was performed in the absence of ceftriaxone pre-treatment, potentially masking subtle changes in the abundance of Pbps in the *salB* mutant at the cell wall (41). Therefore, it remains unclear if SalB partners with or coordinates the activity of Pbps to induce cephalosporin resistance. A *sagA* mutant described in our study (81R5; G460D) has a mutation residing two residues upstream from a peptidoglycan clamp residue (W462) and lacked enhanced susceptibility to ampicillin and ceftriaxone. The reason for this exception and why this mutation confers phage resistance is unclear.

The enhanced susceptibility to β-lactams in *epaR* and *epaX* in *E. faecium* mutants was surprising given prior reports of increased β-lactam resistance in *epa* mutants in *E. faecalis* (46). However, we note that all *epa* mutants tested in that analysis harbored mutations in genes from the core region of *epa* locus (i.e. *epaA, epaE, epaL, epaN, epaB*). To the best of our knowledge, this is the first report demonstrating enhanced β-lactam sensitivity to *epa* variable region mutants in enterococci. We observed enhanced susceptibility to daptomycin in *E. faecium epaR* and *epaX* mutants, consistent with data from *E. faecalis epaR* and *epaX* mutants (9, 14, 47). Given that the *epa* variable genes have recently been discovered to be involved in teichoic acid biosynthesis (25), we hypothesize that altered display of teichoic acids at the cell surface enables the differential β-lactam and daptomycin susceptibility observed in *epa* core versus variable region mutants in enterococci. This hypothesis is supported by observations in *Staphylococcus aureus*, where metabolic perturbations leading to enhanced teichoic acid output or teichoic acid D-alanylation correlate with daptomycin tolerance (48-50). Similarly, mutation of *lafB*, a gene encoding lipoteichoic acid glycosyltransferase, induces a daptomycin hypersusceptible phenotype in *E. faecium* (51). Mutation of *bgsB* in *E. faecalis*, which functions with a *lafB* homolog (*bgsA*) in lipoteichoic acid anchor biosynthesis, results in enhanced susceptibility to daptomycin (14). A reduction in susceptibility to the β-lactam piperacillin in *lafB* (*E. faecium*) or *bgsB* (*E. faecalis*) mutants is reminiscent of the effect of *epa* core region mutations in enterococci (46). A similar pattern of enhanced daptomycin susceptibility at the cost of reduced β-lactam susceptibility, known as the see-saw effect (52), suggests that the altered display or abundance of the wall teichoic acids at the cell surface may occur in response to the modification of rhamnopolysaccharide or lipoteichoic acid. Mutation of *epaR* or *epaX* in *E. faecium* would potentially avoid the daptomycin-β-lactam see-saw effect, making phages that induce these mutations in enterococci attractive antimicrobial candidates. Collectively, these observations suggest that the location of *epa* mutations, core versus variable region, as well mutations in genes participating in teichoic acid biosynthesis, are likely to impact the trajectory of β-lactam and daptomycin susceptibility in enterococci.

*E. faecalis epa* mutations are detrimental during intestinal colonization and show reduced virulence in a mouse peritonitis infection model (9, 53, 54). *epa* mutants are more susceptible to bile salts, neutrophils, exhibit reduced biofilm formation, and are unable to invade biotic and abiotic surfaces (47, 54-56). Therefore, we predict that *epaR* and *epaX* mutants in *E. faecium* will show a similar intestinal colonization dysfunction. Hydrolase-domain mutations in SagA are also likely to induce fitness costs *in vivo*. SagA was shown to promote *E. faecium* attachment to multiple connective tissue molecules, including fibrinogen, fibronectin, and collagen (57). Interestingly, peptidoglycan fragments released following SagA hydrolytic activity activates NOD2-mediated mucosal immunity in the intestine, providing protection from *Salmonella enterica* infection and *Clostridioides difficile* pathogenesis (22, 24). In *E. faecalis*, mutation of the *sagA*-like gene *salB* altered cell morphology, increased biofilm formation, impacted autolysis, and increased susceptibility to bile salts, detergent, ethanol, peroxide, and heat (58-61). Contrary to SagA, cells expressing SalB were limited in binding fibronectin and collagen type I, suggesting that these proteins exhibit different adherence capacities to host tissue. Considering these observations together, it is possible that phage predation that promotes the formation of *sagA* mutants would result in *E. faecium* cells that are compromised for adherence and/or invasion of host tissues, and potentially less immunostimulatory during infection.

The absence of phage-antibiotic synergy for phage resistant strains harboring capsule, topoisomerase 3 (*topB*), and the *rdd* gene does not imply that these mutations do not come with a fitness cost for other antimicrobial agents. We selectively chose to examine beta-lactams and daptomycin in this study given their clinical relevance for treating enterococcal infections. Sensitization to other antibiotics that target pathways other than cell wall biogenesis or membrane stability may exist and remain to be tested. The presence of phage-antibiotic synergy might be a function of how a mutation inhibits phage infection. For instance, bacterial mutations that limit phage adsorption and/or genome ejection (i.e. *epa, sagA*) into the host cell might result in sensitivity to agents acting at the cell wall or membrane, while mutations that might inhibit phage genome replication (i.e. *topB*) could sensitize cells to agents that block bacterial DNA replication. Sensitization of *E. coli* to novobiocin, a topoisomerase inhibitor, following mutation of *topB* (a type I topoisomerase) supports this theory (62). Additionally, purified capsule from *Streptococcus pneumoniae* was shown to protect an acapsular mutant of *Klebsiella pneumoniae* from polymyxin B (63), a lipopeptide antibiotic that normally binds lipopolysaccharide in the outer membrane of Gram-negative bacteria (64). *E. faecalis epa* variable locus mutants are also sensitized to polymyxin B (30), suggesting phage resistance sensitizes enterococci to other antibiotics for which they exhibit intrinsic resistance (64).

Although phage-antibiotic synergy represents an enticing approach for treatment of multi-drug resistant *E. faecium* infections, the narrow host range observed for phages 9181 and 9183 (Fig. 3A-B) will need to be addressed in future studies. Ideal phages would exhibit broader host range activity while retaining synergy with antibiotics. Whether such phages exist in the natural environment is unclear. If not, phage recombineering methods offer promise for precisely broadening the host range of phages (65-67).

Despite their narrow host range, phages 9181 and 9183 can discriminate between members of *E. faecium* clade B. The inability of phage 9183 to infect *E. faecium* Com12 despite this strain exhibiting a near identical *epa* locus compared to 1,141,733 (i.e. *epa* variant 2) (17, 18), suggests phage 9183 might adsorb to Com12 but is unable to infect the cell either due to an inability to bind a secondary receptor, intracellular restriction, or failure to effectively lyse the cell following intracellular phage replication and assembly. What drives phage 9181 specificity for *E. faecium* Com12 and Com15, but not 1,141,733 is not clear. Given the near identical SagA amino acid sequences between *E. faecium* Com12 and 1,141,733 (Identity 97%, Positives 97%, Gaps 2%) suggests that architecture of the peptidoglycan or a mechanism highlighted above for phage 9183 resistance enable *E. faecium* 1,141,733 resistance to phage 9181 infection. Future studies will seek to identify further host factors that constrain the host range of these phages.

Phage 9184 infects both clade A and B strains. Given the heterogenous nature of the capsule loci between the strains infected by phage 9184, it is difficult to ascertain an attribute of this loci that might serve as a determinant of phage 9184 adsorption and infectivity. Acknowledging that the arginine-aspartate-aspartate (RDD) protein has yet to be proven as a receptor for phage 9184, the conservation of this protein in *E. faecium* Com12, which is not infected by phage 9184, suggests that RDD is not the factor limiting infection of this strain.

In conclusion, we have identified three previously undescribed phages that infect *E. faecium*. The study of *E. faecium* resistance to these phages identified multiple components of the *E. faecium* cell surface to be critical for productive phage infection. The enhanced sensitivity of *sagA, epaR* and *epaX* mutants to cell wall and membrane acting antimicrobials suggests that these proteins represent intriguing antimicrobial targets to be considered for future drug discovery efforts against *E. faecium*, and potentially other Gram-positive pathogens harboring homologs of these genes. The finding that *E. faecium* phages synergize with β-lactam and lipopeptide antibiotics provides encouragement that phages could be used in combination with these antibiotics to increase their efficacy and possibly repurpose such antibiotics that are currently deemed ineffective against enterococci.

## Materials and Methods

### Bacteria and bacteriophages

A complete list of the bacterial strains and bacteriophages used in this study can be found in Table S4. *E. faecium* Com12 was cultured in Todd-Hewitt broth (THB) and *E. faecium* 1,141,733 was cultured in brain heart infusion (BHI) broth at 37°C with rotation at 250 rpm. *E. coli* strains were cultured in Lennox L broth (LB) at 37°C with rotation at 250 rpm. Semi-solid media in petri plates were made by adding 1.5% agar to broth prior to autoclaving. For antibiotic susceptibility testing, Mueller Hinton Broth (MHB) was used. When needed, chloramphenicol was added to media at 20 μg/ml or 10 μg/ml for selection of *E. coli* or *E. faecium,* respectively. Phage susceptibility assays were performed on THB agar supplemented with 10 mM MgSO_4_.

### Bacteriophage isolation and purification

Phages 9181, 9183, and 9184 were isolated from wastewater obtained from a water treatment facility located near Denver, Colorado. Fifty milliliters of raw sewage was centrifuged at 3220 x *g* for 10 minutes at room temperature to remove debris. The supernatant was decanted and passed through a 0.45 μm filter. A 100 μl aliquot of filtered wastewater was mixed with 130 μl of *E. faecium* 1,141,733 or Com12 diluted 1:10 from an overnight culture and incubated at room temperature for 15 min. Molten THB top agar (0.35%), supplemented with 10 mM MgSO_4_, was added to the bacteria-wastewater suspension and poured over a 1.5% THB agar plate supplemented with 10 mM MgSO_4_. Following overnight growth at 37°C, plaques were picked with a sterile Pasteur pipette and phages were eluted from the plaque in 500 μl SM-plus buffer (100 mM NaCl, 50 mM Tris-HCl, 8 mM MgSO_4_, 5 mM CaCl_2_ [pH 7.4]) overnight (O/N) at 4°C. After O/N elution, the phages were filter sterilized (0.45 μm). This procedure was repeated two more times to ensure clonal phage isolates. To amplify phages to high titer stocks, 10-fold serially diluted clonal phage isolates were mixed with their appropriate host strain diluted 1:10 from an O/N culture, incubated at room temperature and then poured over 1.5% THB agar supplemented with 10 mM MgSO_4_. Top agar from multiple near confluent lysed bacterial lawns were scraped into a 15 ml conical tube and centrifuged at 18000 x *g* for 10 minutes prior to decanting and 0.45 μm filter sterilization. Using these recovered phages, high-titer phage stocks were generated by infecting 500 mL of early logarithmically (2-3 x 10^8^ CFU/mL) growing *E. faecium* with phage at a multiplicity of infection of 0.5 following supplementation of media with 10 mM MgSO_4_. The phage-cell suspension was incubated at room temperature for 15 min and then incubated at 37°C with rotation (200 rpm) for 4-6 hours. The cultures were centrifuged at 3220 x *g* for 10 minutes at 4°C and the supernatants filtered (0.45 μm). Clarified and filtered lysates were treated with 5 μg/ml each of DNase and RNase at room temperature for 1 hour and phages were precipitated with 1 M NaCl and 10% (wt/vol) polyethylene glycol 8000 (PEG 8000) on ice at 4°C overnight. Phage precipitates were pelleted by centrifugation at 11,270 x *g* for 20 minutes and resuspended in 2 mL of SM-plus buffer. One-third volume chloroform was mixed by inversion into the phage precipitates and centrifuged at 16,300 x *g* to separate out residual PEG 8000 into the organic phase. Phages in the aqueous phase were further purified using a cesium chloride gradient as described previously (11). The final titer was confirmed by plaque assay. Crude phage lysates were used for all phage susceptibility and adsorption assays, while cesium chloride gradient purified phages were used for phage genomic DNA isolation and transmission electron microscopy.

### Transmission electron microscopy

8 μl of 1 x 10^10^ pfu/mL of phages was applied to a copper mesh grid coated with formvar and carbon (Electron Microscopy Sciences) for 2 minutes and then gently blotted off with a piece of Whatman filter paper. The grids were rinsed by transferring between two drops of MilliQ water, blotting with Whatman filter paper between each transfer. Finally, the grids were stained using two drops of a 0.75% uranyl formate solution (a quick rinse with MilliQ water following the first drop followed by an additional 20 seconds of staining). After rinsing and blotting, the grids were allowed to dry for at least 10 minutes. Samples were imaged on a FEI Tecnai G2 Biotwin TEM at 80kV with an AMT side-mount digital camera.

### Whole-genome sequence analysis of phages and phage-resistant bacteria

Phage DNA was isolated by incubating phages with 50 μg/mL proteinase K and 0.5% sodium dodecyl sulfate at 56°C for 1 hour followed by extraction with an equal volume of phenol/chloroform. The aqueous phase was extracted a second time with an equal volume of chloroform and the DNA was precipitated using isopropanol. Bacterial DNA was isolated using a ZymoBIOMICS DNA miniprep kit (Zymo Research), following the manufacturers protocol. Phage and bacterial DNA samples were sequenced at the Microbial Genome Sequencing Center, University of Pittsburgh, using an Illumina NextSeq 550 platform and paired end chemistry (2 x 150bp). Paired-end reads were trimmed and assembled into contigs using CLC genomics workbench (Qiagen). Open reading frames (ORFs) were detected and annotated using rapid annotation subsystem technology (RAST) and the Phage Galaxy structural annotation (version 2020.1) and functional workflows (version 2020.3) (68, 69). Trimmed bacterial genomic reads for *E. faecium* Com12, 1,141,733, and phage resistant derivatives were mapped to reference genomes (GCF_000157635.1 (Com12); GCA_000157575.1 (1,141,733)), downloaded from the National Center for Biotechnology Information (NCBI) website. To identify mutations conferring phage resistance the basic variant detection tool from CLC genomics workbench was used to identify polymorphisms (similarity fraction = 0.5 and length fraction = 0.8).

### PCR screen for phage lysogeny

PCR primers to screen for phage 9181, 9183, and 9184 lysogeny were designed to target the phage lysin (phage 9181 and 9184) or integrase (phage 9183) genes (Table S4). PCR was performed using GoTaq Green master mix (Promega), per the manufacturer’s instructions. Bands were visualized from PCR reactions following electrophoresis of 10 μL of each reaction loaded onto 1% (phage 9184) or 1.5% (phage 9181 and 9183) agarose gels embedded with ethidium bromide. Predicted PCR product sizes were as follows: phage 9181 lysin gene (phi9181_ORF001), 432bp; phage 9183 integrase gene (phi9183_ORF077), 514bp; phage 9184 lysin gene (phi9184_ORF022), 801bp.

### Enterococcal phage orthology analysis

Enterococcal phage orthology was performed according to a method described by Bolocan et al. (16). Briefly, publicly available enterococcal genomes were downloaded from the Millard Lab phage genome database (http://millardlab.org/bioinformatics/). As of May 15, 2020, there were 99 complete enterococcal phage genomes. Open reading frames for each enterococcal phage genome were called using Prodigal and bacteriophage protein Orthologous Groups were identified by OrthoMCL (20, 70). The resulting OrthoMCL matrix was used to generate an orthology tree using the ggplot2 and ggdendro packages in R. Nearest neighbor phages to phages 9181, 9183, and 9184 from the OrthoMCL analysis were compared using the genome alignment feature of ViP Tree using normalized tBLASTx scores between viral genomes to calculate genomic distance for phylogenetic proteomic tree analysis (71).

### Routine molecular techniques, DNA sequencing, and complementation

Confirmation PCRs were performed using GoTaq Green master mix (Promega), per the manufacturer’s instructions. Q5 DNA polymerase master mix (New England Biolabs) was used for PCR reactions intended for cloning, per the manufacturer’s instructions. Plasmid DNA was purified using a QIAprep Miniprep kit (Qiagen) or a ZymoPURE II Plasmid Midiprep kit (Zymo Research). Restriction enzymes and T4 ligase were purchased from New England Biolabs. Sanger DNA sequencing was performed by Quintara Biosciences (San Francisco, CA). A complete list of primers can be found in Table S4. Complementation was performed using plasmid pLZ12A, a derivative of pLZ12 (72) carrying the *bacA* promoter upstream of the multiple cloning site (9). *wze, epaX, dltA*, and *efsg_rs08090* were cloned into pLZ12A as BamHI and EcoRI fragments. *epaR* and *efsg_rs08120* were cloned into pLZ12A as BamHI and PstI fragments. Plasmids were transformed into *E. faecium* using a previously described glycine-sucrose method (73, 74). Briefly, 1 ml of overnight culture was inoculated into 50 ml of BHI supplemented with 2% glycine and 0.5 M sucrose and grown overnight at 37°C with rotation (250 rpm). The following day the cells were pelleted at 7200 x *g* and re-suspended in an equal volume of pre-warmed BHI supplemented with 2% glycine and 0.5 M sucrose and incubated for 1h at 37°C statically. The cells were pelleted at 7200 x *g* and washed three times in ice cold electroporation buffer (0.5 M sucrose and 10% glycerol). 1-2 μg of plasmid DNA was electroporated into *E. faecium* using a Gene Pulser (Bio-Rad) with a 0.2 mm cuvette at 1.7kV, 200 Ω and 25 μF.

### Phage susceptibility assay

Overnight bacterial cultures were pelleted, re-suspended in SM-plus buffer, and normalized to OD_600_ of 1.0. 10-fold serial dilutions of bacteria were spotted on THB agar embedded with phage or THB agar alone, supplemented with 10 mM MgSO_4_. Phages were embedded at the following concentrations within THB agar: phage 9181 (10^8^ PFU/ml), phage 9183 (10^7^ PFU/ml), and phage 9184 (10^7^ PFU/ml). Plates were incubated overnight at 37°C and viable CFU was determined by colony counting.

### Isolation of phage-resistant *E. faecium* strains

130 μl of a 1:10 dilution of *E. faecium* grown O/N was mixed with 10 μl of 10-fold serially diluted phages and added to 5 ml of pre-warmed THB top agar (0.35% wt/vol). Phage-bacterium mixtures were poured onto the surface of THB agar plates (1.5% wt/vol). The plates were incubated at 37°C until phage-resistant colonies appeared in the zones of clearing. The presumptive resistant colonies were passaged four times by streaking single colonies onto THB agar.

### Determination of phage host range

The host range of phages 9181, 9183, and 9184 were determined using a panel of laboratory and contemporary clinical *E. faecium* and *E. faecalis* isolates (Table S4). Overnight bacterial cultures were suspended in SM-plus buffer to an OD_600_ of 1.0 and 10-fold serially diluted and spotted on to THB agar containing phages. Plates were incubated O/N at 37°C and viable CFU was determined. Strains that exhibited greater than 4-log killing in the presence of phage were termed phage susceptible, while those that grew beyond this threshold were considered phage resistant.

### Efficiency of plating

Bacterial strains of interest were selected from a single colony to inoculate 3 mL THB and incubated O/N at 37°C with agitation. Bacteria from O/N cultures were 1:10 diluted in SM-plus buffer and 130 μL was mixed gently with 10 μL of phage 9181 and phage 9184 serially 1:10 diluted in SM-plus buffer using starting titers of 5 x 10^6^ pfu and 1 x 10^7^ pfu, respectively. This phage and bacterial mixture was incubated at room temperature for 15 minutes to enable phage attachment before 5 mL of pre-warmed THB soft agar (0.35% wt/vol) supplemented with 10 mM magnesium sulfate was mixed in and spread over the surface of THB agar (1.5% wt/vol) supplemented with 10 mM magnesium sulfate. The soft agar was allowed to solidify and then incubated O/N at 37°C with the petri dish upright to prevent dislodgement of the soft agar. Phage plaques were enumerated following O/N incubation. Given our difficulties forming an evenly spread plaque layer of phage 9184 on *E. faecium* U37 (75), 6 μL of 10-fold serial dilutions of phage 9184 were spotted onto bacterial lawns of *E. faecium* U37 or 1,141,733 embedded in THB soft agar (0.35 wt/vol) supplemented with 10 mM magnesium sulfate containing either. Following drying of spots, plates were incubated upright O/N at 37°C. Plaques were enumerated following O/N incubation.

### Western blot analysis for SagA from bacterial supernatants and pellets

Western blot was performed as described previously (22). Briefly, bacteria were grown to exponential phase (OD_600_ ~0.8) in BHI. 1 mL exponential phase culture samples were centrifuged at ≥ 18,000 x g and the supernatants were transferred to a new microcentrifuge tube for preparation. Supernatants were prepared as follows: 10% trichloroacetic acid (final; vol/vol) was added and tubes placed at −20°C for 15 minutes for protein precipitation. Tubes were then spun at max speed for 15 minutes, supernatant discarded, and the protein pellet was washed twice with 500 μL cold acetone. Tubes were transferred to a 95°C heat block with caps open to evaporate acetone and dry protein pellets. 60 μL 4% sodium dodecyl sulfate (SDS) buffer (4% SDS, 50 mM Bis-Tris pH 7.5, 150 mM sodium chloride, 1X Laemmli Buffer, 2.5% β-mercaptoethanol) was added to the protein pellets, which were sonicated for 5 minutes to solubilize. The samples were then placed at 95°C for 5 minutes to denature proteins. Cell pellets were prepared as follows: 1 mL cell pellets were resuspended with 1 mL phosphate buffered saline, transferred to 2 mL cryovials, and centrifuged at 5000 x g for 5 minutes to wash. After discarding supernatant, 50 μL 0.1 mm beads were added followed by 250 μL 4% SDS buffer (see above). Cryovial tubes were placed in a FastPrep FP120 cell disruptor at max speed for 20 seconds on and 10 seconds off, and this process was repeated twice more. Tubes were then centrifuged at 5000 x g for one minute and then placed at 95°C for 10 minutes to remove bubbles & denature proteins. 15 μL of 60 μL supernatant or cell pellet sample was loaded for SDS-PAGE. Proteins were separated by SDS-PAGE on 4-20% Criterion TGX precast gels (Bio-Rad), then transferred to nitrocellulose membrane (0.2 mM, BioTrace NT Nitrocellulose Transfer Membranes, Pall Laboratory). For SagA blots, polyclonal SagA serum and HRP conjugated anti-Rabbit IgG (GE Healthcare, NA 934V) served as primary and secondary antibodies, respectively. Polyclonal SagA primary antibodies and secondary antibody were used at a dilution of 1:25000 and 1:10000 (supernatants) or 1:10000 and 1:5000 (cell pellets), respectively. Membranes were blocked for one hour in TBS-T (Tris-buffered saline, 0.1% Tween 20) containing 5% non-fat milk, incubated with blocking buffer containing primary antibody for one hour, washed five times with TBS-T, incubated with blocking buffer containing secondary antibody, and washed four times with TBS-T. Protein detection was performed with ECL detection reagent (GE Healthcare) on a Bio-Rad ChemiDoc MP Imaging System.

### Bacterial growth curves

250 mL BHI was inoculated with 2.5 mL (1:100) overnight cultures and incubated at 37°C with agitation until OD_600_ ~0.8.

### Phage adsorption assay

This assay was performed as described previously (10, 11). O/N bacterial cultures were pelleted at 3220 × *g* for 10 min and resuspended to 10^8^ CFU/mL in SM-plus buffer. Phage adsorption was determined by mixing 5 x 10^6^ pfu of phage and to 5 x 10^7^ cfu of the appropriate bacterial strain in 500 μL and incubating statically at room temperature for 10 min. The bacteria-phage suspensions were centrifuged at 24,000 × *g* for 1 min, the supernatant was collected, and remaining phages enumerated by a plaque assay. SM-plus buffer with phage only (no bacteria) served as a control. Percent adsorption was determined as follows: [(PFU_control_ - PFU_test supernatant_)/PFU_control_] × 100. The fold change was calculated by dividing the percent adsorption of phage resistant mutants by those of parental strain.

### Antibiotic MIC assay

Antibiotic MIC was determined for each strain using Etest strips (bioMérieux). Single colonies were grown O/N in 3 mL of MHB broth at 37°C with rotation (250 rpm). The following day overnight cultures were diluted to McFarland 0.5 in MHB broth and 100 μL of the cell suspension was spread over the surface of MHB agar plates. One Etest strip was placed on the surface of the agar using sterile forceps. The plates were incubated for 18 hours at 37°C. The MIC was determined to be the number closest to the zone of inhibition. The mean and standard deviation of the MIC from three independent experiments is reported for each strain.

### Phage-antibiotic synergy assay

O/N cultures of *E. faecium* Com12 and *E. faecium* 1,141,733 were normalized to 10^8^ CFU/mL. 100 μl (10^7^ CFU/mL) of bacteria was added into a sterile 96-well plate in triplicate. Antibiotics were diluted 1:100 into desired wells to achieve the appropriate final concentration. Phages were added to desired wells at 10^6^ PFU/mL, achieving a multiplicity of infection of 0.1. The 96 well plate was loaded on to BioTek Synergy Plate reader pre-warmed to 37°C, and agitated continuously for 18h, allowing for OD_600_ reading every 30 minutes.

## Data availability

The Illumina reads for phage 9181, 9183, and 9184 and phage-resistant *E. faecium* mutants have been deposited in the European Nucleotide Archive under the accession number PRJEB39873. Assembled phage genomes were submitted to Genbank and were assigned the following accession numbers: MT939240 (phage 9181), MT939241 (phage 9183), and MT939242 (phage 9184).

## Acknowledgements

This work was supported by National Institutes of Health grants R01AI141479 (B.A.D.), R01GM103593 (H.C.H), R01CA245292 (H.C.H.), T32AI070084 (J.E.), and T32AR007534 (M.R.M). J.E. acknowledges support from The Rockefeller University Graduate Program and H.C.H. acknowledges support from a Kenneth Rainin Foundation Synergy Award. We thank Dr. Jennifer Bourne at the University of Colorado School of Medicine Electron Microscopy Center for preparing and visualizing electron micrographs of phages. We thank the staff at the Microbial Genome Sequencing Center (MiGS) at the University of Pittsburgh for assistance with bacterial and phage whole genome DNA sequencing.

## Supplemental Figure Legends

**Figure S1. PCR screen for phage lysogeny in phage resistant mutants.** A molecular weight marker with corresponding band sizes in base pairs (i.e. bp) is shown at the far left and right of each gel image. (A) PCR screen for phage 9181 lysin gene in *E. faecium* or phage 9181 genomic DNA. Lane numbers correspond to the following genomic DNA samples: 1) 81R3, 2) 81R4, 3) 81R5, 4) 81R6, 5) 81R7, 6) 81R8, 7) *E. faecium* Com12, 8) Phage 9181, 9) negative control. (B) PCR screen for phage 9183 integrase gene in *E. faecium* or phage 9183 genomic DNA. Lane numbers correspond to the following genomic DNA samples: 1) 83R1, 2) 83R2, 3) 83R3, 4) 83R4, 5) 83R5, 6) 83R6, 7) 83R7, 8) 83R8, 9) *E. faecium* 1,141,733, 10) Phage 9183, 11) negative control. (C) PCR screen for phage 9184 lysin gene in *E. faecium* or phage 9184 genomic DNA. Lane numbers correspond to the following genomic DNA samples: 1) 84R1,2) 84R2, 3) 84R3, 4) 84R4, 5) 84R5, 6) 84R6, 7) 84R8, 8) *E. faecium* 1,141,733, 9) phage 9184, 10) negative control.

**Figure S2. *Enterococcus faecium* phage orthoclusters.** Phage protein coding sequence alignments were performed with nearest neighbors in VIP Tree (71, 76). Colored lines connecting genomes indicate percent protein identity along the length of each genome. (A) Phage 9183 demonstrates protein homology and similar genome organization to its nearest neighbor intra-orthocluster phages (phages VFW and VPE25). (B) Phage 9184 demonstrates proteome homology and similar genome organization to its nearest neighbor intra-orthocluster phages (phages vB EfaS-DELF1 and IME-EFm5). (C) Phage 9181 shows little to no protein homology to its nearest neighbor extra-orthocluster phages (phages EFC-1 and FLA4).

**Figure S3. Phage resistant mutants of *E. faecium* following exposure to phages 9181, 9183 and 9184.** Phage 9181 (A), 9183 (B), and 9184 (C) susceptibility assays and associated bacterial enumeration of wild type and phage resistant mutants in the presence (white bars) or absence (black bars) of phage (A-F) from three independent experiments. Phage 9181 (A, D) and phage 9183 (B, E) resistant strains exhibit ≥ 5-logs of survival versus *E. faecium* Com12 and 1,141,733 (i.e. 733), respectively. Phage 9184 (C, F) resistant strains exhibit a weak resistance phenotypes. The dotted line indicates the spontaneous mutation threshold conferring phage resistance observed in the respective wild type host strain of each phage. The threshold was placed to aid in discriminating weak phage resistance phenotypes versus the parental strain.

**Figure S4. SagA is conserved in *E. faecium* Com12 and Com15 and SagA is expressed in *sagA* mutants.** (A) Displayed is the BLASTP alignment of SagA between Com12 and Com15, showing 95% similarity and strict conservation of peptidoglycan clamp (orange lettering) and active site residues (red lettering). Colored highlights indicate the location of amino acid changes detected in phage 9181 resistant mutants (81R3 and 81R4 - green highlight, 81R5 - yellow highlight, 81R6 - blue highlight, 81R8 - magenta highlight). Specific amino acid changes are noted below the alignment in parentheses next to their respective phage resistant mutant. (B) Growth of *E. faecalis* OG1RF, *E. faecium* Com15, Com12 (WT) and *sagA* mutants (81R3-6; 81R8) are similar in BHI, except for 81R6. (C,D) Displayed is the whole protein fraction (upper panel; Stain-Free) and Western Blot of SagA (lower panel; α-SagA) taken from the exponential phase (OD_600_ ~0.8) supernatants (C) or cell pellets (D) of *E. faecalis* OG1RF, *E. faecium* Com15, Com12 (WT), and *sagA* mutants (81R3-6; 81R8). Protein band sizes are demonstrated to the left of each panel in kilodaltons (kDa).

**Figure S5. Complementation restores phage susceptibility in phage resistant mutants.** Bacterial enumeration from Phage 9181 (A and B), 9184 (C), and 9183 (D) phage susceptibility assays of wild type and phage resistant mutants complemented with their respective wild type allele or empty vector. Assays were performed in the presence (white bars) or absence (black bars) of phages from two independent experiments. The bars and error bars indicate the average and standard deviation from two independent experiments. The dotted line indicates the spontaneous mutation threshold conferring phage resistance observed in the respective wild type host strain of each phage.

**Figure S6. Complementation restores phage adsorption in phage resistant mutants.** Percentage phage adsorption of phage resistant mutants, complemented phage resistant mutants, or their parental strains to Phage 9181 (A), 9183 (B), and 9184 (C). Parental and phage resistant mutants were complemented with the empty vector (E; pLZ12a) and compared to their complemented phage resistant mutant strain. Data represent the mean percent adsorption and standard deviation from three independent experiments. *, *P* < 0.05; ***, *P* < 0.001; ****, *P* < 0.0001; ns, non-significant by unpaired Student’s *t* test.

## References

1. Arias CA, Murray BE. 2012. The rise of the Enterococcus: beyond vancomycin resistance. Nat Rev Microbiol 10:266–78. doi:10.1038/nrmicro2761.

2. Kristich CJ, Rice LB, Arias CA. 2014. Enterococcal infection—treatment and antibiotic resistance, p 1–62. In Gilmore MS, Clewell DB, Ike Y, Shakar N (ed), Enterococci: from commensals to leading causes of drug resistant infection [Internet]. Massachusetts Eye and Ear Infirmary, Boston, MA.

3. Beganovic M, Luther MK, Rice LB, Arias CA, Rybak MJ, LaPlante KL. 2018. A review of combination antimicrobial therapy for *Enterococcus faecalis* bloodstream infections and infective endocarditis. Clin Infect Dis 67:303–309. doi:10.1093/cid/ciy064.

4. Rybak MJ, McGrath BJ. 1996. Combination antimicrobial therapy for bacterial infections. Guidelines for the clinician. Drugs 52:390–405. doi:10.2165/00003495-199652030-00005.

5. Fish R, Kutter E, Bryan D, Wheat G, Kuhl S. 2018. Resolving digital staphylococcal osteomyelitis using bacteriophage-a case report. Antibiotics (Basel) 7:87. doi:10.3390/antibiotics7040087.

6. Cano EJ, Caflisch KM, Bollyky PL, Van Belleghem JD, Patel R, Fackler J, Brownstein MJ, Horne B, Biswas B, Henry M, Malagon F, Lewallen DG, Suh GA. 2020. Phage therapy for limb-threatening prosthetic knee *Klebsiella pneumoniae* infection: case report and in vitro characterization of anti-biofilm activity. Clin Infect Dis doi:10.1093/cid/ciaa705. doi:10.1093/cid/ciaa705.

7. Schooley RT, Biswas B, Gill JJ, Hernandez-Morales A, Lancaster J, Lessor L, Barr JJ, Reed SL, Rohwer F, Benler S, Segall AM, Taplitz R, Smith DM, Kerr K, Kumaraswamy M, Nizet V, Lin L, McCauley MD, Strathdee SA, Benson CA, Pope RK, Leroux BM, Picel AC, Mateczun AJ, Cilwa KE, Regeimbal JM, Estrella LA, Wolfe DM, Henry MS, Quinones J, Salka S, Bishop-Lilly KA, Young R, Hamilton T. 2017. Development and use of personalized bacteriophage-based therapeutic cocktails to treat a patient with a disseminated resistant *Acinetobacter baumannii* infection. Antimicrob Agents Chemother 61:e00954–17. doi:10.1128/AAC.00954-17.

8. Chan BK, Turner PE, Kim S, Mojibian HR, Elefteriades JA, Narayan D. 2018. Phage treatment of an aortic graft infected with *Pseudomonas aeruginosa*. Evol Med Public Health 2018:60–66. doi:10.1093/emph/eoy005.

9. Chatterjee A, Johnson CN, Luong P, Hullahalli K, McBride SW, Schubert AM, Palmer KL, Carlson PE, Jr., Duerkop BA. 2019. Bacteriophage resistance alters antibiotic-mediated intestinal expansion of enterococci. Infect Immun 87:e00085–19. doi:10.1128/iai.00085-19.

10. Chatterjee A, Willett JLE, Nguyen UT, Monogue B, Palmer KL, Dunny GM, Duerkop BA. 2020. Parallel genomics uncover novel enterococcal-bacteriophage interactions. mBio 11:e03120–19. doi:10.1128/mBio.03120-19.

11. Duerkop BA, Huo W, Bhardwaj P, Palmer KL, Hooper LV. 2016. Molecular basis for lytic bacteriophage resistance in enterococci. mBio 7:e01304–16. doi:10.1128/mBio.01304-16.

12. Wandro S, Oliver A, Gallagher T, Weihe C, England W, Martiny JBH, Whiteson K. 2018. Predictable molecular adaptation of coevolving *Enterococcus faecium* and lytic phage EfV12-phi1. Front Microbiol 9:3192. doi:10.3389/fmicb.2018.03192.

13. Lossouarn J, Briet A, Moncaut E, Furlan S, Bouteau A, Son O, Leroy M, DuBow MS, Lecointe F, Serror P, Petit MA. 2019. *Enterococcus faecalis* countermeasures defeat a virulent Picovirinae bacteriophage. Viruses 11:48. doi:10.3390/v11010048.

14. Ho K, Huo W, Pas S, Dao R, Palmer KL. 2018. Loss-of-function mutations in *epaR* confer resistance to phiNPV1 Infection in *Enterococcus faecalis* OG1RF. Antimicrob Agents Chemother 62:e00758–18. doi:10.1128/AAC.00758-18.

15. Duerkop BA, Palmer KL, Horsburgh MJ. 2014. Enterococcal bacteriophages and genome defense, p 1–44. In Gilmore MS, Clewell DB, Ike Y, Shankar N (ed), Enterococci: from commensals to leading causes of drug resistant infection. Massachusetts Eye and Ear Infirmary, Boston.

16. Bolocan AS, Upadrasta A, Bettio PHA, Clooney AG, Draper LA, Ross RP, Hill C. 2019. Evaluation of phage therapy in the context of *Enterococcus faecalis* and its associated diseases. Viruses 11:366. doi:10.3390/v11040366.

17. Palmer KL, Godfrey P, Griggs A, Kos VN, Zucker J, Desjardins C, Cerqueira G, Gevers D, Walker S, Wortman J, Feldgarden M, Haas B, Birren B, Gilmore MS. 2012. Comparative genomics of enterococci: variation in *Enterococcus faecalis*, clade structure in *E. faecium*, and defining characteristics of *E. gallinarum* and *E. casseliflavus*. mBio 3:e00318–11. doi:10.1128/mBio.00318-11.

18. de Been M, van Schaik W, Cheng L, Corander J, Willems RJ. 2013. Recent recombination events in the core genome are associated with adaptive evolution in *Enterococcus faecium*. Genome Biol Evol 5:1524–35. doi:10.1093/gbe/evt111.

19. Ackermann HW. 2007. 5500 Phages examined in the electron microscope. Arch Virol 152:227–43. doi:10.1007/s00705-006-0849-1.

20. Li L, Stoeckert CJ, Jr., Roos DS. 2003. OrthoMCL: identification of ortholog groups for eukaryotic genomes. Genome Res 13:2178–89. doi:10.1101/gr.1224503.

21. Holtzman T, Globus R, Molshanski-Mor S, Ben-Shem A, Yosef I, Qimron U. 2020. A continuous evolution system for contracting the host range of bacteriophage T7. Sci Rep 10:307. doi:10.1038/s41598-019-57221-0.

22. Kim B, Wang YC, Hespen CW, Espinosa J, Salje J, Rangan KJ, Oren DA, Kang JY, Pedicord VA, Hang HC. 2019. *Enterococcus faecium* secreted antigen A generates muropeptides to enhance host immunity and limit bacterial pathogenesis. Elife 8:e45343. doi:10.7554/eLife.45343.

23. Ittisoponpisan S, Islam SA, Khanna T, Alhuzimi E, David A, Sternberg MJE. 2019. Can predicted protein 3D structures provide reliable insights into whether missense variants are disease associated? J Mol Biol 431:2197–2212. doi:10.1016/j.jmb.2019.04.009.

24. Rangan KJ, Pedicord VA, Wang YC, Kim B, Lu Y, Shaham S, Mucida D, Hang HC. 2016. A secreted bacterial peptidoglycan hydrolase enhances tolerance to enteric pathogens. Science 353:1434–1437. doi:10.1126/science.aaf3552.

25. Guerardel Y, Sadovskaya I, Maes E, Furlan S, Chapot-Chartier MP, Mesnage S, Rigottier-Gois L, Serror P. 2020. Complete structure of the enterococcal polysaccharide antigen (EPA) of vancomycin-resistant *Enterococcus faecalis* V583 Reveals that EPA decorations are teichoic acids covalently linked to a rhamnopolysaccharide backbone. mBio 11:e00277–20. doi:10.1128/mBio.00277-20.

26. Kortright KE, Chan BK, Koff JL, Turner PE. 2019. Phage therapy: a renewed approach to combat antibiotic-resistant bacteria. Cell Host Microbe 25:219–232. doi:10.1016/j.chom.2019.01.014.

27. Mangalea MR, Duerkop BA. 2020. Fitness trade-offs resulting from bacteriophage resistance potentiate synergistic antibacterial strategies. Infect Immun 88:e00926–19. doi:10.1128/IAI.00926-19.

28. Morrisette T, Lev KL, Kebriaei R, Abdul-Mutakabbir J, Stamper KC, Morales S, Lehman SM, Canfield GS, Duerkop BA, Arias CA, Rybak MJ. 2020. Bacteriophage-antibiotic combinations for *Enterococcus faecium* with varying bacteriophage and daptomycin susceptibilities. Antimicrob Agents Chemother 64:e00993–20. doi:10.1128/AAC.00993-20.

29. Al-Zubidi M, Widziolek M, Court EK, Gains AF, Smith RE, Ansbro K, Alrafaie A, Evans C, Murdoch C, Mesnage S, Douglas CWI, Rawlinson A, Stafford GP. 2019. Identification of Novel Bacteriophages with Therapeutic Potential That Target *Enterococcus faecalis*. Infect Immun 87:e00512–19. doi:10.1128/iai.00512-19.

30. Smith RE, Salamaga B, Szkuta P, Hajdamowicz N, Prajsnar TK, Bulmer GS, Fontaine T, Kolodziejczyk J, Herry JM, Hounslow AM, Williamson MP, Serror P, Mesnage S. 2019. Decoration of the enterococcal polysaccharide antigen EPA is essential for virulence, cell surface charge and interaction with effectors of the innate immune system. PLoS Pathog 15:e1007730. doi:10.1371/journal.ppat.1007730.

31. Rodriguez C, Van der Meulen R, Vaningelgem F, Font de Valdez G, Raya R, De Vuyst L, Mozzi F. 2008. Sensitivity of capsular-producing *Streptococcus thermophilus* strains to bacteriophage adsorption. Lett Appl Microbiol 46:462–8. doi:10.1111/j.1472-765X.2008.02341.x.

32. Rice LB, Lakticová V, Helfand MS, Hutton-Thomas R. 2004. In vitro antienterococcal activity explains associations between exposures to antimicrobial agents and risk of colonization by multiresistant enterococci. J Infect Dis 190:2162–6. doi:10.1086/425580.

33. Ubeda C, Taur Y, Jenq RR, Equinda MJ, Son T, Samstein M, Viale A, Socci ND, van den Brink MR, Kamboj M, Pamer EG. 2010. Vancomycin-resistant *Enterococcus* domination of intestinal microbiota is enabled by antibiotic treatment in mice and precedes bloodstream invasion in humans. J Clin Invest 120:4332–41. doi:10.1172/JCI43918.

34. Brandl K, Plitas G, Mihu CN, Ubeda C, Jia T, Fleisher M, Schnabl B, DeMatteo RP, Pamer EG. 2008. Vancomycin-resistant enterococci exploit antibiotic-induced innate immune deficits. Nature 455:804–7. doi:10.1038/nature07250.

35. Donskey CJ, Chowdhry TK, Hecker MT, Hoyen CK, Hanrahan JA, Hujer AM, Hutton-Thomas RA, Whalen CC, Bonomo RA, Rice LB. 2000. Effect of antibiotic therapy on the density of vancomycin-resistant enterococci in the stool of colonized patients. N Engl J Med 343:1925–32. doi:10.1056/NEJM200012283432604.

36. Hendrickx AP, Top J, Bayjanov JR, Kemperman H, Rogers MR, Paganelli FL, Bonten MJ, Willems RJ. 2015. Antibiotic-driven dysbiosis mediates intraluminal agglutination and alternative segregation of *Enterococcus faecium* from the intestinal epithelium. mBio 6:e01346–15. doi:10.1128/mBio.01346-15.

37. Duan Y, Llorente C, Lang S, Brandl K, Chu H, Jiang L, White RC, Clarke TH, Nguyen K, Torralba M, Shao Y, Liu J, Hernandez-Morales A, Lessor L, Rahman IR, Miyamoto Y, Ly M, Gao B, Sun W, Kiesel R, Hutmacher F, Lee S, Ventura-Cots M, Bosques-Padilla F, Verna EC, Abraldes JG, Brown RS, Jr., Vargas V, Altamirano J, Caballeria J, Shawcross DL, Ho SB, Louvet A, Lucey MR, Mathurin P, Garcia-Tsao G, Bataller R, Tu XM, Eckmann L, van der Donk WA, Young R, Lawley TD, Starkel P, Pride D, Fouts DE, Schnabl B. 2019. Bacteriophage targeting of gut bacterium attenuates alcoholic liver disease. Nature 575:505–511. doi:10.1038/s41586-019-1742-x.

38. Stein-Thoeringer CK, Nichols KB, Lazrak A, Docampo MD, Slingerland AE, Slingerland JB, Clurman AG, Armijo G, Gomes ALC, Shono Y, Staffas A, Burgos da Silva M, Devlin SM, Markey KA, Bajic D, Pinedo R, Tsakmaklis A, Littmann ER, Pastore A, Taur Y, Monette S, Arcila ME, Pickard AJ, Maloy M, Wright RJ, Amoretti LA, Fontana E, Pham D, Jamal MA, Weber D, Sung AD, Hashimoto D, Scheid C, Xavier JB, Messina JA, Romero K, Lew M, Bush A, Bohannon L, Hayasaka K, Hasegawa Y, Vehreschild M, Cross JR, Ponce DM, Perales MA, Giralt SA, Jenq RR, Teshima T, Holler E, Chao NJ, et al. 2019. Lactose drives *Enterococcus* expansion to promote graft-versus-host disease. Science 366:1143–1149. doi:10.1126/science.aax3760.

39. Baddour LM, Wilson WR, Bayer AS, Fowler VG, Jr., Tleyjeh IM, Rybak MJ, Barsic B, Lockhart PB, Gewitz MH, Levison ME, Bolger AF, Steckelberg JM, Baltimore RS, Fink AM, O’Gara P, Taubert KA, American Heart Association Committee on Rheumatic Fever E, Kawasaki Disease of the Council on Cardiovascular Disease in the Young CoCCCoCS, Anesthesia, Stroke C. 2015. Infective endocarditis in adults: diagnosis, antimicrobial therapy, and management of complications: a scientific statement for healthcare professionals from the American Heart Association. Circulation 132:1435–86. doi:10.1161/CIR.0000000000000296.

40. Lorenzo MP, Kidd JM, Jenkins SG, Nicolau DP, Housman ST. 2019. In vitro activity of ampicillin and ceftriaxone against ampicillin-susceptible *Enterococcus faecium*. J Antimicrob Chemother 74:2269–2273. doi:10.1093/jac/dkz173.

41. Djoric D, Little JL, Kristich CJ. 2020. Multiple low-reactivity class B penicillin-binding proteins are required for cephalosporin resistance in enterococci. Antimicrob Agents Chemother 64:e02273–19. doi:10.1128/AAC.02273-19.

42. Arbeloa A, Segal H, Hugonnet JE, Josseaume N, Dubost L, Brouard JP, Gutmann L, Mengin-Lecreulx D, Arthur M. 2004. Role of class A penicillin-binding proteins in PBP5-mediated beta-lactam resistance in Enterococcus faecalis. J Bacteriol 186:1221–8. doi:10.1128/jb.186.5.1221-1228.2004.

43. Rice LB, Carias LL, Rudin S, Hutton R, Marshall S, Hassan M, Josseaume N, Dubost L, Marie A, Arthur M. 2009. Role of class A penicillin-binding proteins in the expression of beta-lactam resistance in *Enterococcus faecium*. J Bacteriol 191:3649–56. doi:10.1128/JB.01834-08.

44. Sifaoui F, Arthur M, Rice L, Gutmann L. 2001. Role of penicillin-binding protein 5 in expression of ampicillin resistance and peptidoglycan structure in *Enterococcus faecium*. Antimicrob Agents Chemother 45:2594–7. doi:10.1128/aac.45.9.2594-2597.2001.

45. Djoric D, Kristich CJ. 2017. Extracellular SalB contributes to intrinsic cephalosporin resistance and cell envelope integrity in *Enterococcus faecalis*. J Bacteriol 199:e00392–17. doi:10.1128/JB.00392-17.

46. Singh KV, Murray BE. 2019. Loss of a major enterococcal polysaccharide antigen (Epa) by *Enterococcus faecalis* is associated with increased resistance to ceftriaxone and carbapenems. Antimicrob Agents Chemother 63:e00481–19. doi:10.1128/aac.00481-19.

47. Dale JL, Cagnazzo J, Phan CQ, Barnes AM, Dunny GM. 2015. Multiple roles for *Enterococcus faecalis* glycosyltransferases in biofilm-associated antibiotic resistance, cell envelope integrity, and conjugative transfer. Antimicrob Agents Chemother 59:4094–105. doi:10.1128/AAC.00344-15.

48. Gaupp R, Lei S, Reed JM, Peisker H, Boyle-Vavra S, Bayer AS, Bischoff M, Herrmann M, Daum RS, Powers R, Somerville GA. 2015. *Staphylococcus aureus* metabolic adaptations during the transition from a daptomycin susceptibility phenotype to a daptomycin nonsusceptibility phenotype. Antimicrob Agents Chemother 59:4226–38. doi:10.1128/AAC.00160-15.

49. Mechler L, Bonetti EJ, Reichert S, Flotenmeyer M, Schrenzel J, Bertram R, Francois P, Gotz F. 2016. Daptomycin tolerance in the *Staphylococcus aureus pitA6* mutant is due to upregulation of the *dlt* operon. Antimicrob Agents Chemother 60:2684–91. doi:10.1128/AAC.03022-15.

50. Bertsche U, Weidenmaier C, Kuehner D, Yang SJ, Baur S, Wanner S, Francois P, Schrenzel J, Yeaman MR, Bayer AS. 2011. Correlation of daptomycin resistance in a clinical *Staphylococcus aureus* strain with increased cell wall teichoic acid production and D-alanylation. Antimicrob Agents Chemother 55:3922–8. doi:10.1128/AAC.01226-10.

51. Mello SS, Van Tyne D, Lebreton F, Silva SQ, Nogueira MCL, Gilmore MS, Camargo I. 2020. A mutation in the glycosyltransferase gene *lafB* causes daptomycin hypersusceptibility in *Enterococcus faecium*. J Antimicrob Chemother 75:36–45. doi:10.1093/jac/dkz403.

52. Sieradzki K, Tomasz A. 1997. Inhibition of cell wall turnover and autolysis by vancomycin in a highly vancomycin-resistant mutant of *Staphylococcus aureus*. Journal of Bacteriology 179:2557–2566. doi:10.1128/jb.179.8.2557-2566.1997.

53. Xu Y, Singh KV, Qin X, Murray BE, Weinstock GM. 2000. Analysis of a gene cluster of *Enterococcus faecalis* involved in polysaccharide biosynthesis. Infect Immun 68:815–23. doi:10.1128/iai.68.2.815-823.2000.

54. Rigottier-Gois L, Madec C, Navickas A, Matos RC, Akary-Lepage E, Mistou MY, Serror P. 2015. The surface rhamnopolysaccharide epa of *Enterococcus faecalis* is a key determinant of intestinal colonization. J Infect Dis 211:62–71. doi:10.1093/infdis/jiu402.

55. Mohamed JA, Huang W, Nallapareddy SR, Teng F, Murray BE. 2004. Influence of origin of isolates, especially endocarditis isolates, and various genes on biofilm formation by *Enterococcus faecalis*. Infect Immun 72:3658–63. doi:10.1128/iai.72.6.3658-3663.2004.

56. Ramos Y, Rocha J, Hael AL, van Gestel J, Vlamakis H, Cywes-Bentley C, Cubillos-Ruiz JR, Pier GB, Gilmore MS, Kolter R, Morales DK. 2019. PolyGlcNAc-containing exopolymers enable surface penetration by non-motile *Enterococcus faecalis*. PLoS Pathog 15:e1007571. doi:10.1371/journal.ppat.1007571.

57. Teng F, Kawalec M, Weinstock GM, Hryniewicz W, Murray BE. 2003. An *Enterococcus faecium* secreted antigen, SagA, exhibits broad-spectrum binding to extracellular matrix proteins and appears essential for *E. faecium* growth. Infect Immun 71:5033–41. doi:10.1128/iai.71.9.5033-5041.2003.

58. Breton YL, Mazé A, Hartke A, Lemarinier S, Auffray Y, Rincé A. 2002. Isolation and characterization of bile salts-sensitive mutants of *Enterococcus faecalis*. Current Microbiology 45:0434–0439. doi:10.1007/s00284-002-3714-3.

59. Rincé A, Le Breton Y, Verneuil N, Giard J-C, Hartke A, Auffray Y. 2003. Physiological and molecular aspects of bile salt response in *Enterococcus faecalis*. International Journal of Food Microbiology 88:207–213. doi:10.1016/s0168-1605(03)00182-x.

60. Mohamed JA, Teng F, Nallapareddy SR, Murray BE. 2006. Pleiotrophic effects of 2 *Enterococcus faecalis sagA*-like genes, *salA* and *salB*, which encode proteins that are antigenic during human infection, on biofilm formation and binding to collagen type i and fibronectin. J Infect Dis 193:231–40. doi:10.1086/498871.

61. Shankar J, Walker RG, Wilkinson MC, Ward D, Horsburgh MJ. 2012. SalB inactivation modulates culture supernatant exoproteins and affects autolysis and viability in *Enterococcus faecalis* OG1RF. J Bacteriol 194:3569–78. doi:10.1128/JB.00376-12.

62. Perez-Cheeks BA, Lee C, Hayama R, Marians KJ. 2012. A role for topoisomerase III in Escherichia coli chromosome segregation. Mol Microbiol 86:1007–22. doi:10.1111/mmi.12039.

63. Llobet E, Tomas JM, Bengoechea JA. 2008. Capsule polysaccharide is a bacterial decoy for antimicrobial peptides. Microbiology (Reading) 154:3877–3886. doi:10.1099/mic.0.2008/022301-0.

64. Zavascki AP, Goldani LZ, Li J, Nation RL. 2007. Polymyxin B for the treatment of multidrug-resistant pathogens: a critical review. Journal of Antimicrobial Chemotherapy 60:1206–1215. doi:10.1093/jac/dkm357.

65. Dunne M, Rupf B, Tala M, Qabrati X, Ernst P, Shen Y, Sumrall E, Heeb L, Pluckthun A, Loessner MJ, Kilcher S. 2019. Reprogramming bacteriophage host range through structure-guided design of chimeric receptor binding proteins. Cell Rep 29:1336–1350 e4. doi:10.1016/j.celrep.2019.09.062.

66. Burrowes BH, Molineux IJ, Fralick JA. 2019. Directed in vitro evolution of therapeutic bacteriophages: the appelmans protocol. Viruses 11:241. doi:10.3390/v11030241.

67. Yehl K, Lemire S, Yang AC, Ando H, Mimee M, Torres MT, de la Fuente-Nunez C, Lu TK. 2019. Engineering phage host-range and suppressing bacterial resistance through phage tail fiber mutagenesis. Cell 179:459–469 e9. doi:10.1016/j.cell.2019.09.015.

68. Aziz RK, Bartels D, Best AA, DeJongh M, Disz T, Edwards RA, Formsma K, Gerdes S, Glass EM, Kubal M, Meyer F, Olsen GJ, Olson R, Osterman AL, Overbeek RA, McNeil LK, Paarmann D, Paczian T, Parrello B, Pusch GD, Reich C, Stevens R, Vassieva O, Vonstein V, Wilke A, Zagnitko O. 2008. The RAST Server: rapid annotations using subsystems technology. BMC Genomics 9:75. doi:10.1186/1471-2164-9-75.

69. Mijalis E, Rasche H. 2013-2017. CPT galaxy tools. https://github.com/tamu-cpt/galaxy-tools/ Accessed 1 May 2020.

70. Hyatt D, Chen GL, Locascio PF, Land ML, Larimer FW, Hauser LJ. 2010. Prodigal: prokaryotic gene recognition and translation initiation site identification. BMC Bioinformatics 11:119. doi:10.1186/1471-2105-11-119.

71. Nishimura Y, Yoshida T, Kuronishi M, Uehara H, Ogata H, Goto S. 2017. ViPTree: the viral proteomic tree server. Bioinformatics 33:2379–2380. doi:10.1093/bioinformatics/btx157.

72. Perez-Casal J, Caparon MG, Scott JR. 1991. Mry, a trans-acting positive regulator of the M protein gene of *Streptococcus pyogenes* with similarity to the receptor proteins of two-component regulatory systems. J Bacteriol 173:2617–24. doi:10.1128/jb.173.8.2617-2624.1991.

73. Shepard BD, Gilmore MS. 1995. Electroporation and efficient transformation of *Enterococcus faecalis* grown in high concentrations of glycine. Methods Mol Biol 47:217–26. doi:10.1385/0-89603-310-4:217.

74. Zhang X, Paganelli FL, Bierschenk D, Kuipers A, Bonten MJ, Willems RJ, van Schaik W. 2012. Genome-wide identification of ampicillin resistance determinants in *Enterococcus faecium*. PLoS Genet 8:e1002804. doi:10.1371/journal.pgen.1002804.

75. Rice LB, Carias LL, Donskey CL, Rudin SD. 1998. Transferable, plasmid-mediated VanB-type glycopeptide resistance in *Enterococcus faecium*. Antimicrob Agents Chemother 42:963–4.

76. Mihara T, Nishimura Y, Shimizu Y, Nishiyama H, Yoshikawa G, Uehara H, Hingamp P, Goto S, Ogata H. 2016. Linking virus genomes with host taxonomy. Viruses 8:66. doi:10.3390/v8030066.

