## Supplementary figures and images for "Lytic bacteriophages facilitate antibiotic sensitization of *Enterococcus faecium*"

### Supplemental Figure 1

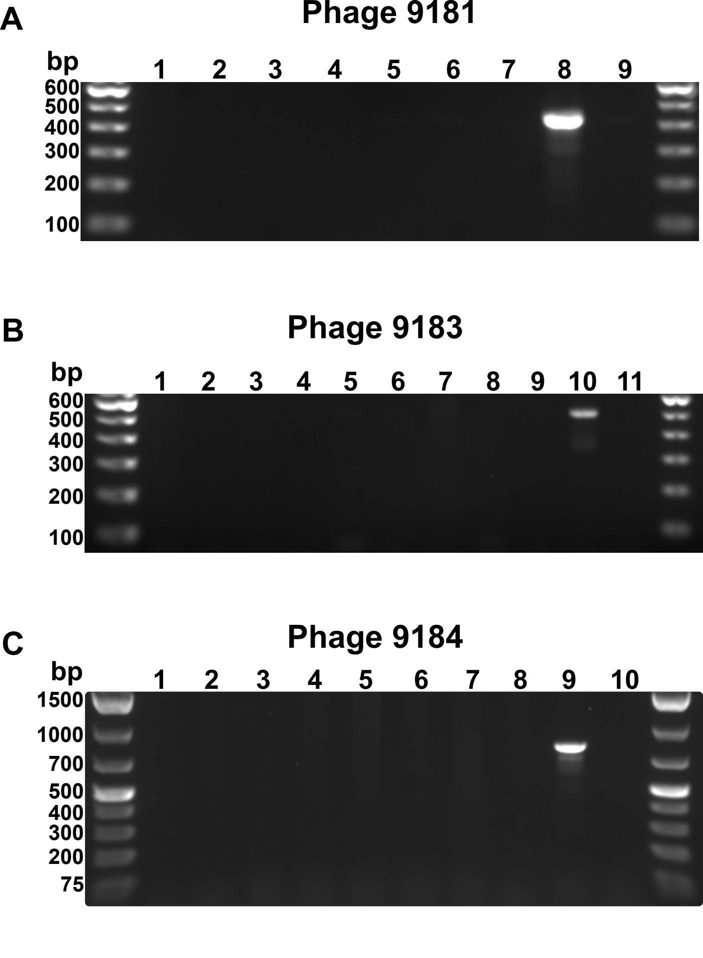

### Supplemental Figure 2

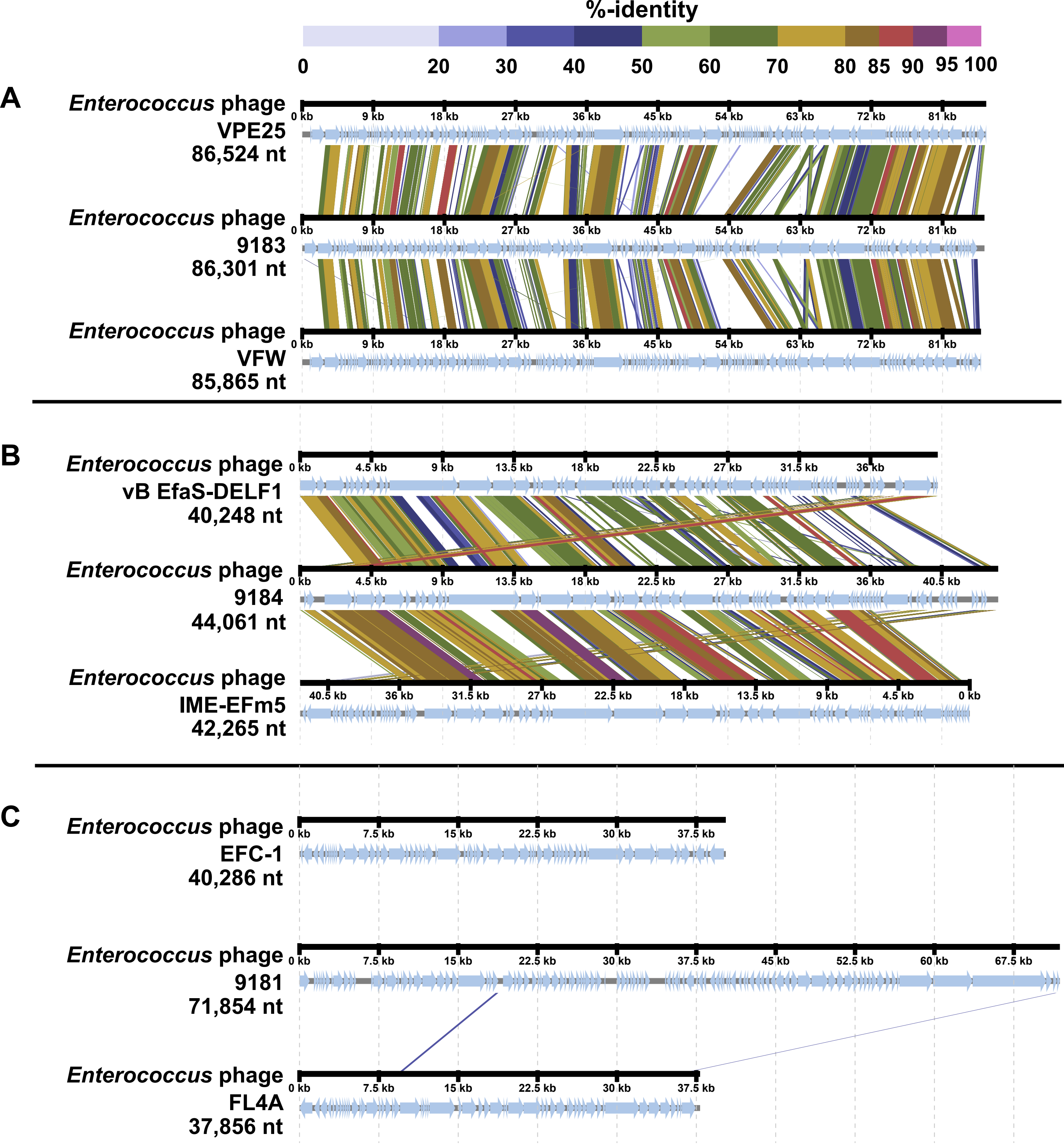

### Supplemental Figure 3

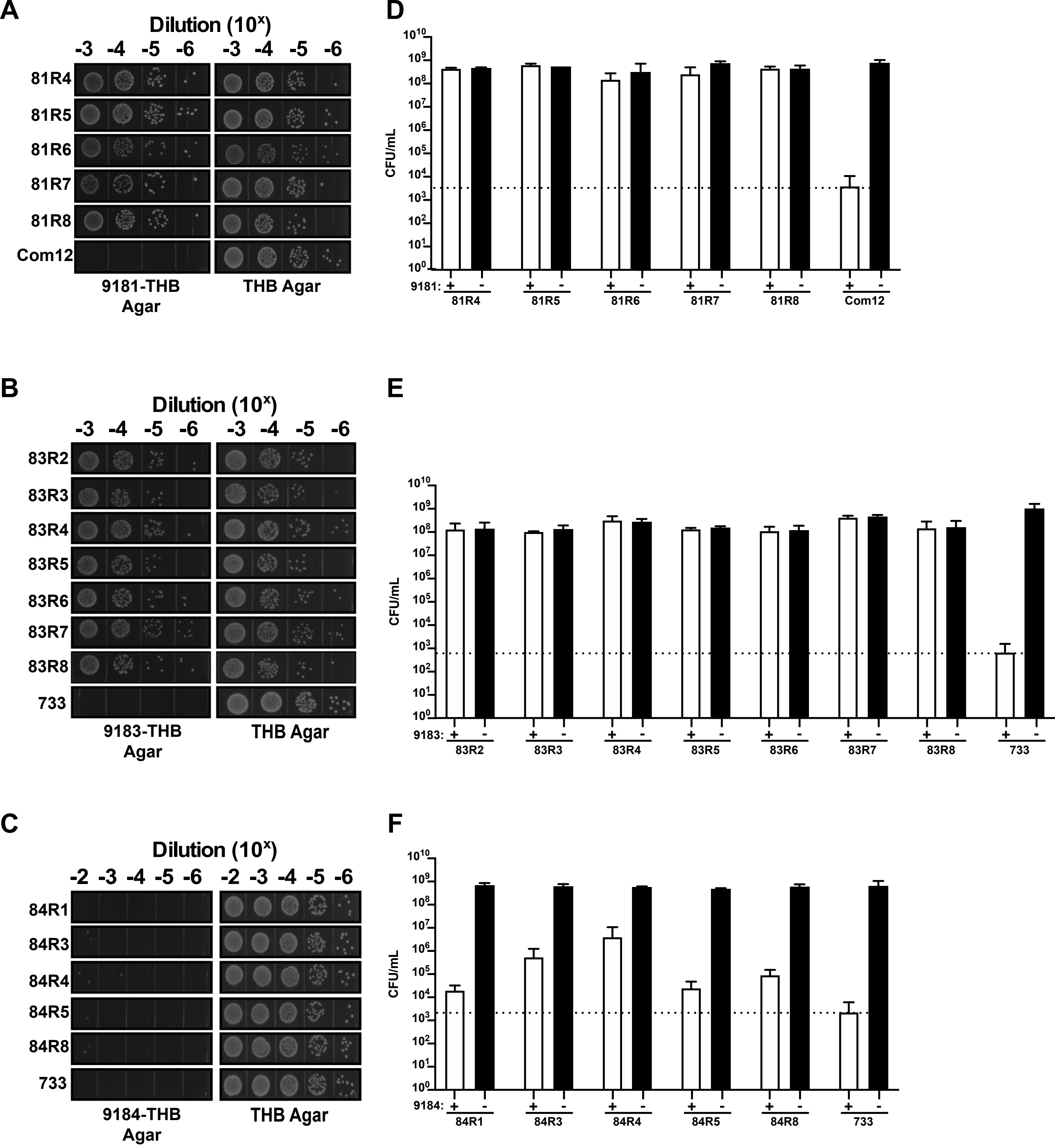

### Supplemental Figure 5

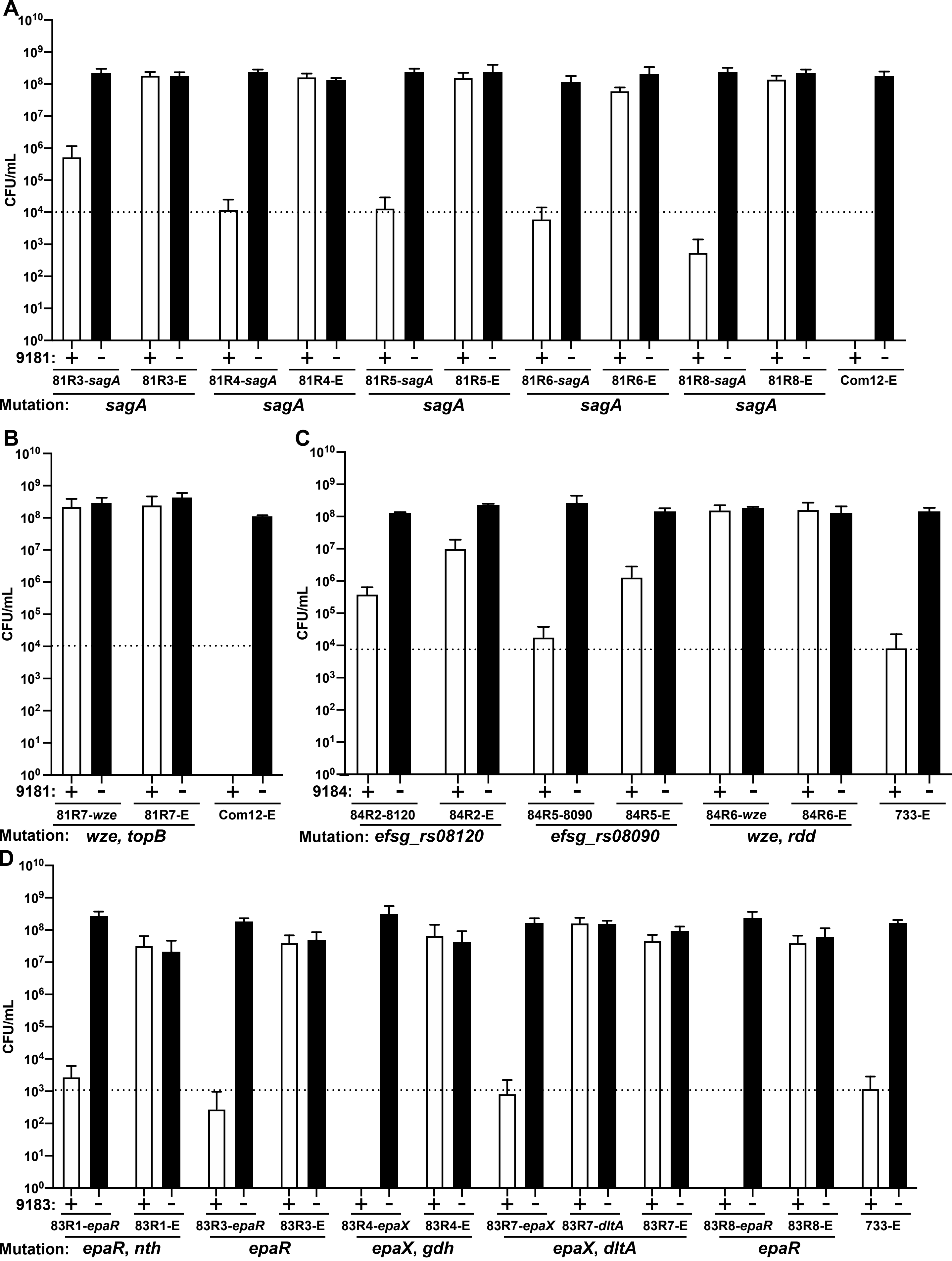

### Supplemental Figure 6

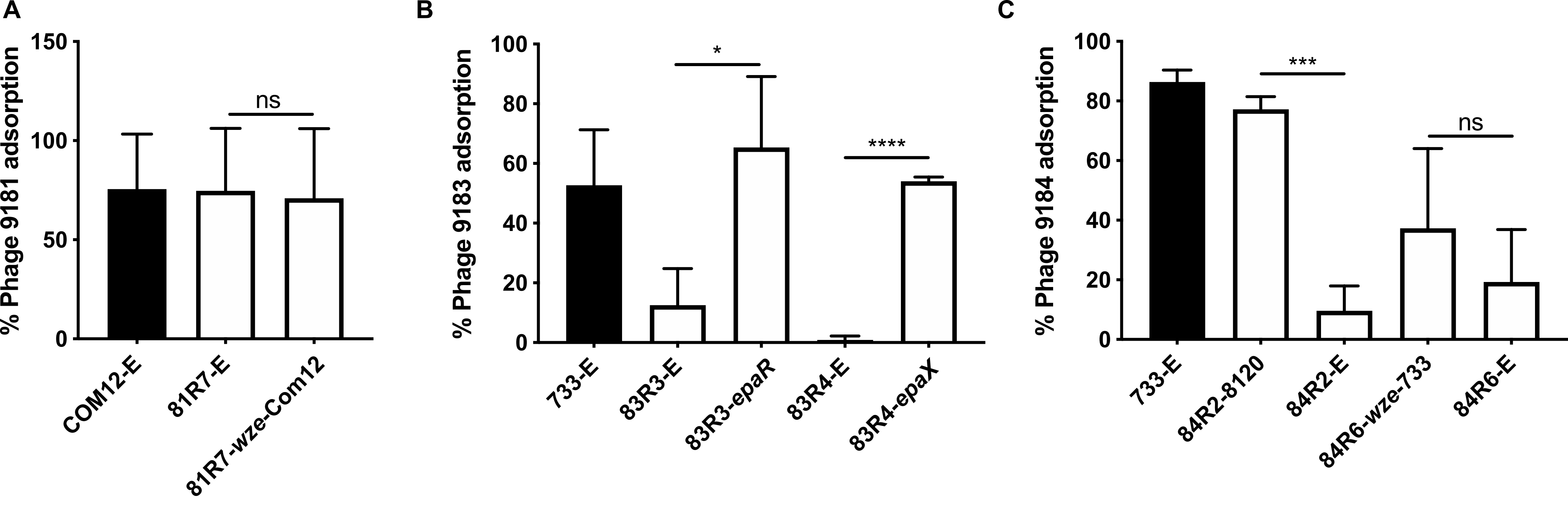
