## Supplemental Figure 4 for "Lytic bacteriophages facilitate antibiotic sensitization of *Enterococcus faecium*"

**A**

| Identities | Positives | Gaps |
|---|---|---|
| 513/538(95%) | 515/538(95%) | 20/538(3%) |

```
Com12  1    VKKSLISAVMVCSMTLTAVASPIAAAADDFDSQIQQQDQKIADLKNQQADAQSQIDALES  60
            +KKSLISAVMVCSMTLTAVASPIAAAADDFDSQIQQQDQKIADLKNQQADAQSQIDALES
Com15  1    MKKSLISAVMVCSMTLTAVASPIAAAADDFDSQIQQQDQKIADLKNQQADAQSQIDALES  60

Com12  61   QVSEINTQAQDLLAKQDTLRQESAQLVKDIADLQERIEKREDTIQKQAREAQVSNTSSNY  120
            QVSEINTQAQDLLAKQDTLRQESAQLVKDIADLQERIEKREDTIQKQAREAQVSNTSSNY
Com15  61   QVSEINTQAQDLLAKQDTLRQESAQLVKDIADLQERIEKREDTIQKQAREAQVSNTSSNY  120

Com12  121  IDAVLNADSLADAIGRVQAMTTMVKANNDLMEQQKQDKKAVEDKKAENDAKLKELAENQA  180
            IDAVLNADSLADAIGRVQAMTTMVKANNDLMEQQKQDKKAVEDKKAENDAKLKELAENQA
Com15  121  IDAVLNADSLADAIGRVQAMTTMVKANNDLMEQQKQDKKAVEDKKAENDAKLKELAENQA  180

Com12  181  ALESQKGDLLSKQADLNVLKTSLAAEQATAEDKKADLNRQKAEAEAEQARIREQQRLAEQ  240
            ALESQKGDLLSKQADLNVLKTSLAAEQATAEDKKADLNRQKAEAEAEQARIREQQRLAEQ
Com15  181  ALESQKGDLLSKQADLNVLKTSLAAEQATAEDKKADLNRQKAEAEAEQARIREQQRLAEQ  240

Com12  241  ARQQAAQEKAEKEAREQAEAEAQATQASSTAQSSASEESSAAQSSTTEESSSAAQSSTTE  300
            ARQQAAQEKAEKEAREQAEAEAQATQASS AQSSA+EESSAAQSSTTEESSSAAQSSTTE
Com15  241  ARQQAAQEKAEKEAREQAEAEAQATQASSAAQSSATEESSAAQSSTTEESSSAAQSSTTE  300

Com12  301  ESTTAPESSTTEESTTAPESSTTEESTTVPESSTTEESTTVPESSTTEESTTVPESSTTE  360
            ESTTAPESSTTEESTT PESSTTEESTT PESSTTEESTTVPESSTTEESTTVPES
Com15  301  ESTTAPESSTTEESTTXPESSTTEESTTAPESSTTEESTTVPESSTTEESTTVPES----  356

Com12  361  ESTTVPETSTEESTTPAPTTPSTDQSVDPGNSTGSNATNNT--------TNTTPTPTPSG  412
                    STEESTTPAPTTPSTDQSVDPGNSTGSNATNNT        TNTTPTPTPSG
Com15  357  --------STEESTTPAPTTPSTDQSVDPGNSTGSNATNNTTNTTPXXXTNTTPTPTPSG  408

Com12  413  SVNGAAIVAEAYKYIGTPYVWGGKDPSGFDCSGFTRYVYLQVTGRDIGGWTVPQESAGTK  472
            SVNGAAIVAEAYKYIGTPYVWGGKDPSGFDCSGFTRYVYLQVTGRDIGGWTVPQESAGTK
Com15  409  SVNGAAIVAEAYKYIGTPYVWGGKDPSGFDCSGFTRYVYLQVTGRDIGGWTVPQESAGTK  468

Com12  473  ISVSQAKAGDLLFWGSPGGTYHVAIALGGGQYIHAPQPGESVKVGSVQWFAPDFAVSM  530
            ISVSQAKAGDLLFWGSPGGTYHVAIALGGGQYIHAPQPGESVKVGSVQWFAPDFAVSM
Com15  469  ISVSQAKAGDLLFWGSPGGTYHVAIALGGGQYIHAPQPGESVKVGSVQWFAPDFAVSM  526
```

81R3 and 81R4 (W433G and W433C, respectively)
81R5 (G460D)
81R6 (note: L insertion between Y451 and L452
81R8 (G435V)
Active Site Residues (C443, H494, H506)
Peptidoglycan Clamp Residues (W433 and W462)

**B**

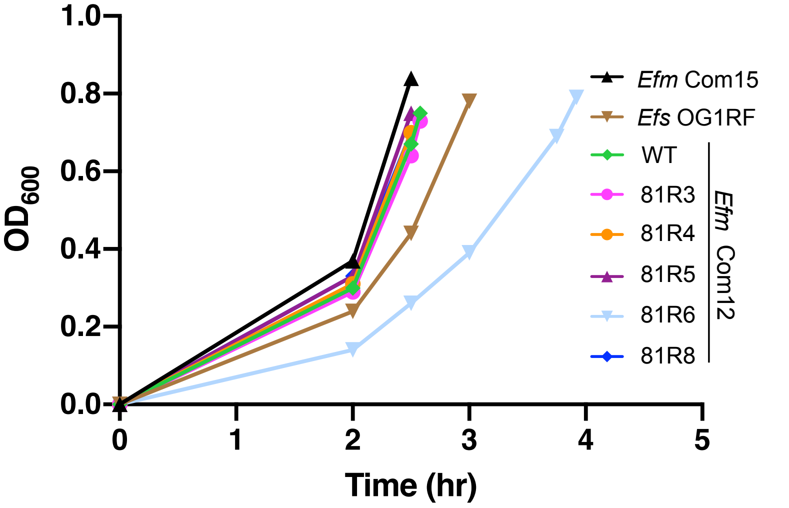


**C** **D**

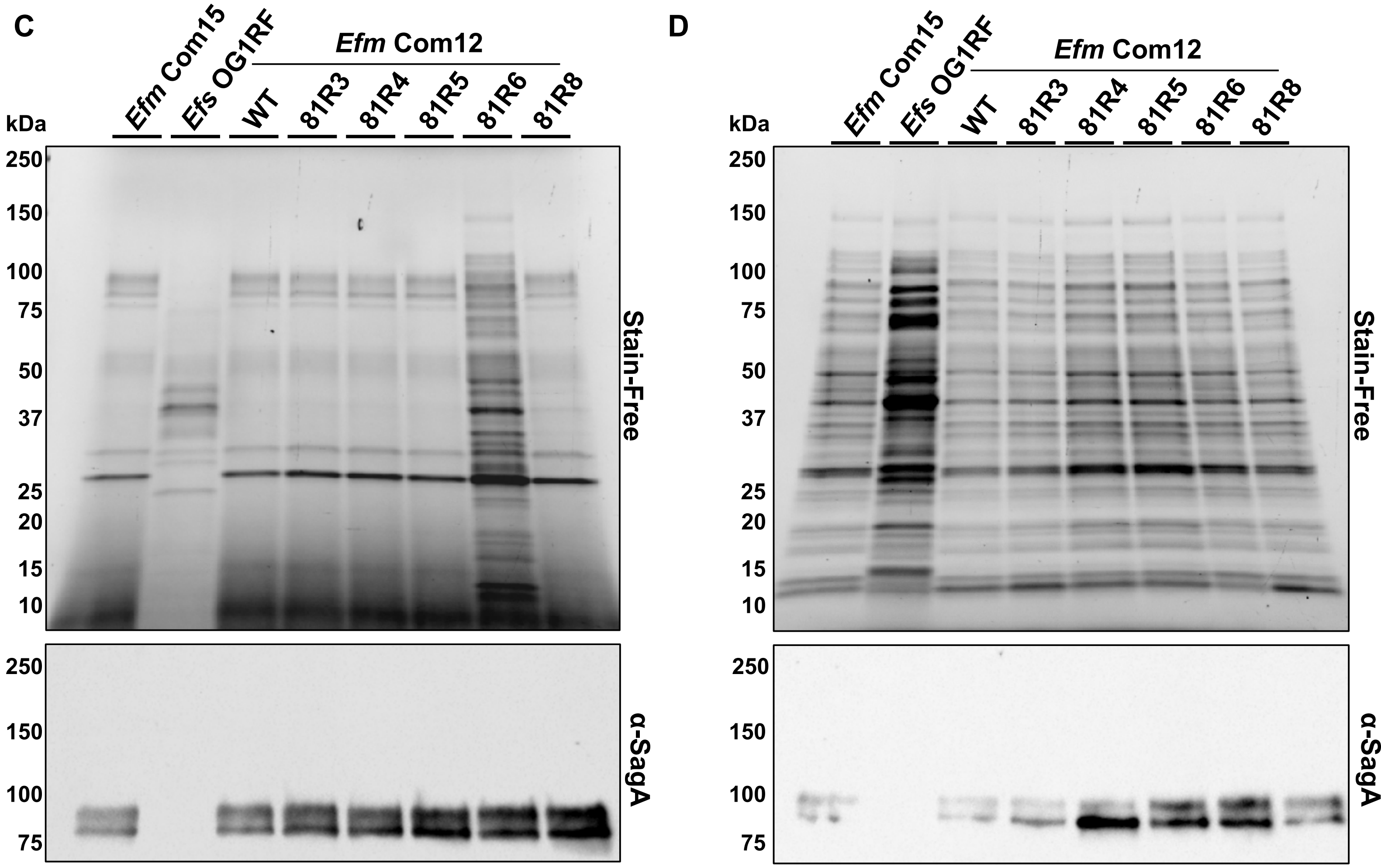


**Figure S4**
