## Supplementary Table 1 for "Lytic bacteriophages facilitate antibiotic sensitization of *Enterococcus faecium*"

**Table S1A. *Enterococcus faecium* phage 9181 genome organization and features**

**Phage 9181 genome annotation**

| Feature ID | Gene Start | Gene Stop | Length | Strand | blastp Best-Hit cut-off (0.001) Number | Best-Hit Accession | % Gaps | % Identities | % Positives | E-value | Predicted Function |
|---|---|---|---|---|---|---|---|---|---|---|---|
| ORF1 | 15 | 959 | 945 | + | N-acetylmuramoyl-L-alanine amidase Enterococcus hirae | OJG47739 | 6 | 53 | 66 | 7.88E-98 | N-acetylmuramoyl-L-alanine amidase |
| ORF2 | 1362 | 1601 | 240 | + | | | | | | | hypothetical protein |
| ORF3 | 1605 | 1853 | 249 | + | | | | | | | hypothetical protein |
| ORF4 | 1866 | 2048 | 183 | + | | | | | | | hypothetical protein |
| ORF5 | 2045 | 2185 | 141 | + | | | | | | | hypothetical protein |
| ORF6 | 2199 | 2483 | 285 | + | | | | | | | hypothetical protein |
| ORF7 | 2496 | 2906 | 411 | + | hypothetical protein EFP29_60 Enterococcus phage EF-P29 | APU00246 | 0 | 45 | 68 | 2.70E-31 | hypothetical protein |
| ORF8 | 3020 | 3178 | 159 | + | hypothetical protein Enterococcus casseliflavus | WP_010749468 | 0 | 43 | 61 | 5.02E-04 | hypothetical protein |
| ORF9 | 3193 | 4158 | 966 | + | hypothetical protein X878_0035 Enterococcus phage VD13 | YP_009036396 | 10 | 47 | 64 | 6.70E-70 | hypothetical protein |
| ORF10 | 4191 | 4730 | 540 | + | | | | | | | hypothetical protein |
| ORF11 | 4727 | 5146 | 420 | + | | | | | | | hypothetical protein |
| ORF12 | 5148 | 5348 | 201 | + | Enterococcus phage IMEEF1 | YP_009603964 | 3 | 58 | 70 | 8.41E-15 | hypothetical protein |
| ORF13 | 6777 | 6971 | 195 | + | | | | | | | hypothetical protein |
| ORF14 | 6976 | 7929 | 954 | + | hypothetical protein Enterococcus phage vB_EfaS_Ef2.2 | QBZ69248 | 1 | 50 | 70 | 7.56E-109 | DNA primase |
| ORF15 | 7990 | 8268 | 279 | + | | | | | | | hypothetical protein |
| ORF16 | 8268 | 9251 | 984 | + | hypothetical protein Enterococcus wangshanyuanii | WP_088271390 | 10 | 36 | 53 | 1.69E-51 | Rnl2 family RNA ligase |
| ORF17 | 9253 | 9459 | 207 | + | hypothetical protein Enterococcus faecium | WP_002324739 | 0 | 65 | 76 | 6.90E-08 | hypothetical protein |
| ORF18 | 9469 | 9618 | 150 | + | | | | | | | hypothetical protein |
| ORF19 | 9608 | 9787 | 180 | + | | | | | | | hypothetical protein |
| ORF20 | 9787 | 10323 | 537 | + | hypothetical protein Pseudomonas | WP_092194385 | 9 | 39 | 54 | 4.74E-20 | HNH endonuclease |

| | | | | | | | | | | | |
|---|---|---|---|---|---|---|---|---|---|---|---|
| ORF21 | 10382 | 10759 | 378 | + | hypothetical protein Enterococcus gallinarum | WP_142976764 .1 | 0 | 79 | 79 | 1.00E-20 | ribonucleoside-diphosphate reductase |
| ORF22 | 10795 | 11613 | 819 | + | DNA replication protein Enterococcus phage vB_EfaS_EF1c55 | QEM41680 | 2 | 57 | 75 | 3.19E-95 | DNA replication initiator protein |
| ORF23 | 11610 | 12983 | 1374 | + | replicative DNA helicase Streptococcus phage SPQS1 | YP_008320518 | 1 | 59 | 78 | 0.00E+00 | DNA helicase |
| ORF24 | 12994 | 13791 | 798 | + | | | | | | | hypothetical protein |
| ORF25 | 13788 | 14348 | 561 | + | | | | | | | hypothetical protein |
| ORF26 | 14430 | 14984 | 555 | + | LPS glycosyltransferase Enterococcus phage Entf1 | QDB70555 | 2 | 33 | 49 | 1.26E-05 | LPS glycosyltransferase |
| ORF27 | 14984 | 17605 | 2622 | + | DNA polymerase I Enterococcus phage vB_EfaS_Ef2.2 | QBZ69269 | 4 | 51 | 66 | 0.00E+00 | DNA polymerase I |
| ORF28 | 17671 | 17919 | 249 | + | hypothetical protein | | | | | | hypothetical protein |
| ORF29 | 17919 | 18248 | 330 | + | | | | | | | hypothetical protein |
| ORF30 | 18271 | 18801 | 531 | + | hypothetical protein Enterococcus faecalis | WP_016624495 | 4 | 26 | 55 | 7.23E-12 | hypothetical protein |
| ORF31 | 18896 | 19243 | 348 | + | hypothetical protein Enterococcus phage vB_EfaS_Ef2.2 | QBZ69253 | 0 | 51 | 67 | 2.64E-27 | HNH endonuclease |
| ORF32 | 19227 | 20279 | 1053 | + | hypothetical protein Enterococcus phage vB_EfaS_EF1c55 | QEM41676 | 1 | 54 | 71 | 4.96E-126 | exonuclease |
| ORF33 | 20279 | 20608 | 330 | + | | | | | | | hypothetical protein |
| ORF34 | 20602 | 21168 | 567 | + | crossover junction endodeoxyribonuclease RuvC Enterococcus phage SAP6 | YP_009604008 | 1 | 44 | 65 | 3.79E-44 | crossover junction endodeoxyribonuclease RuvC |
| ORF35 | 21152 | 21742 | 591 | + | hypothetical protein Enterococcus phage vB_EfaS_Ef2.2 | QBZ69257 | 5 | 43 | 64 | 8.23E-39 | adenylate kinase |
| ORF36 | 21717 | 22448 | 732 | + | hypothetical protein A958_gp43 Enterococcus phage BC611 | YP_006488770 | 3 | 31 | 50 | 4.25E-16 | RNA polymerase sigma factor |
| ORF37 | 22519 | 22734 | 216 | + | | | | | | | hypothetical protein |
| ORF38 | 22734 | 22961 | 228 | + | | | | | | | hypothetical protein |
| ORF39 | 22980 | 23780 | 801 | + | hypothetical protein EFP01_126 Enterococcus phage EFP01 | APZ82053 | 0 | 75 | 89 | 1.17E-147 | deoxyadenosine kinase deoxyguanosine kinase |
| ORF40 | 23837 | 24568 | 732 | + | PnuC-like NrdR-regulated deoxyribonucleotide | QDB71576 | 0 | 73 | 87 | 3.66E-126 | PnuC-like NrdR-regulated deoxyribonucleotide transporter |

| | | | | | | | | | | |
|---|---|---|---|---|---|---|---|---|---|---|
| | | | | | transporter Enterococcus phage vB_OCPT_Ben | | | | | |
| ORF41 | 24670 | 25059 | 390 | + | | | | | | hypothetical protein |
| ORF42 | 25056 | 25277 | 222 | + | | | | | | Double-strand recombination repair protein |
| ORF43 | 25270 | 25446 | 177 | + | | | | | | hypothetical protein |
| ORF44 | 25458 | 25661 | 204 | + | | | | | | hypothetical protein |
| ORF45 | 25661 | 26080 | 420 | + | | | | | | hypothetical protein |
| ORF46 | 26064 | 26549 | 486 | + | | | | | | hypothetical protein |
| ORF47 | 26543 | 26797 | 255 | + | | | | | | hypothetical protein |
| ORF48 | 26797 | 27000 | 204 | + | | | | | | hypothetical protein |
| ORF49 | 26997 | 27512 | 516 | + | | | | | | hypothetical protein |
| ORF50 | 27509 | 27958 | 450 | + | | | | | | hypothetical protein |
| ORF51 | 27948 | 28082 | 135 | + | | | | | | hypothetical protein |
| ORF52 | 28219 | 28551 | 333 | + | putative HNH endonuclease Lactococcus phage 340 | YP_008318136 | 8 | 39 | 51 | 1.08E-12 | HNH endonuclease |
| ORF53 | 28806 | 28910 | 105 | - | | | | | | | hypothetical protein |
| ORF54 | 29273 | 29425 | 153 | - | | | | | | | hypothetical protein |
| ORF55 | 29777 | 29890 | 114 | + | | | | | | | hypothetical protein |
| ORF56 | 29993 | 30400 | 408 | + | hypothetical protein PEf771_147 Enterococcus phage PEf771 | QEP29564 | 10 | 41 | 57 | 4.63E-23 | terminal repeat-encoded protein P |
| ORF57 | 30416 | 30883 | 468 | + | | | | | | | hypothetical protein |
| ORF58 | 30887 | 31168 | 282 | + | hypothetical protein EFP01_182 Enterococcus phage EFP01 | APZ82109 | 0 | 46 | 73 | 2.96E-22 | hypothetical protein |
| ORF59 | 31161 | 31466 | 306 | + | | | | | | | hypothetical protein |
| ORF60 | 31488 | 31826 | 339 | + | hypothetical protein Enterococcus phage vB_EfaS_PHB08 | QBX32893 | 5 | 54 | 65 | 1.40E-21 | hypothetical protein |
| ORF61 | 31868 | 32149 | 282 | + | hypothetical protein EFP29_02 Enterococcus phage EF-P29 | APU00188 | 0 | 81 | 89 | 2.59E-46 | hypothetical protein |
| ORF62 | 32237 | 32392 | 156 | + | | | | | | | hypothetical protein |
| ORF63 | 32394 | 32561 | 168 | + | | | | | | | hypothetical protein |

| | | | | | | | | | | | |
|---|---|---|---|---|---|---|---|---|---|---|---|
| ORF64 | 32673 | 33122 | 450 | + | | | | | | | hypothetical protein |
| ORF65 | 33332 | 33427 | 96 | - | | | | | | | hypothetical protein |
| ORF66 | 33635 | 33739 | 105 | + | | | | | | | hypothetical protein |
| ORF67 | 33761 | 33880 | 120 | - | | | | | | | hypothetical protein |
| ORF68 | 33961 | 34110 | 150 | - | | | | | | | hypothetical protein |
| ORF69 | 34175 | 34354 | 180 | + | | | | | | | hypothetical protein |
| ORF70 | 34395 | 34508 | 114 | - | | | | | | | hypothetical protein |
| ORF71 | 34521 | 34652 | 132 | - | | | | | | | hypothetical protein |
| ORF72 | 34698 | 35060 | 363 | - | DUF1642 domain-containing protein Enterococcus gilvus | WP_010781701 | 8 | 34 | 59 | 9.30E-10 | DUF1642 domain-containing protein |
| ORF73 | 35153 | 35398 | 246 | - | | | | | | | hypothetical protein |
| ORF74 | 35395 | 35790 | 396 | - | | | | | | | hypothetical protein |
| ORF75 | 35803 | 35952 | 150 | - | | | | | | | hypothetical protein |
| ORF76 | 36157 | 36384 | 228 | - | | | | | | | hypothetical protein |
| ORF77 | 36507 | 37277 | 771 | - | | | | | | | hypothetical protein |
| ORF78 | 37264 | 37518 | 255 | - | | | | | | | hypothetical protein |
| ORF79 | 37478 | 37690 | 213 | - | | | | | | | hypothetical protein |
| ORF80 | 37671 | 38156 | 486 | - | | | | | | | hypothetical protein |
| ORF81 | 38149 | 38421 | 273 | - | | | | | | | hypothetical protein |
| ORF82 | 38396 | 38488 | 93 | - | | | | | | | hypothetical protein |
| ORF83 | 38778 | 39251 | 474 | - | | | | | | | hypothetical protein |
| ORF84 | 39253 | 39684 | 432 | - | DUF1642 domain-containing protein Enterococcus faecium | WP_142972308 | 11 | 37 | 53 | 4.03E-11 | DUF1642 domain-containing protein |
| ORF85 | 39684 | 39974 | 291 | - | | | | | | | hypothetical protein |
| ORF86 | 39986 | 40141 | 156 | - | hypothetical protein A5816_000554 Enterococcus sp. 3G1_DIV0629 | OTO28288 | 0 | 50 | 75 | 2.88E-04 | hypothetical protein |
| ORF87 | 40141 | 40302 | 162 | - | hypothetical protein HMPREF9524_01966 Enterococcus faecium TX0133a01 | EFR67890 | 0 | 43 | 61 | 3.88E-04 | hypothetical protein |

| | | | | | | | | | | | |
|---|---|---|---|---|---|---|---|---|---|---|---|
| ORF88 | 40299 | 40487 | 189 | - | Uncharacterised protein Enterococcus hirae | VTQ86473 | 0 | 54 | 76 | 9.00E-06 | hypothetical protein |
| ORF89 | 40499 | 40705 | 207 | - | hypothetical protein Enterococcus durans | WP_144775119 | 0 | 99 | 99 | 1.44E-41 | hypothetical protein |
| ORF90 | 40702 | 41226 | 525 | - | DUF1642 domain-containing protein Enterococcus faecium | WP_104889537 | 9 | 52 | 65 | 1.26E-52 | DUF1642 domain-containing protein |
| ORF91 | 41219 | 41428 | 210 | - | hypothetical protein Enterococcus faecium | WP_010722442 | 0 | 99 | 100 | 1.17E-40 | hypothetical protein |
| ORF92 | 41484 | 41765 | 282 | - | | | | | | | hypothetical protein |
| ORF93 | 41762 | 41932 | 171 | - | | | | | | | hypothetical protein |
| ORF94 | 41922 | 42161 | 240 | - | | | | | | | hypothetical protein |
| ORF95 | 42182 | 42397 | 216 | - | | | | | | | HNH endonuclease |
| ORF96 | 42487 | 42951 | 465 | - | | | | | | | hypothetical protein |
| ORF97 | 43031 | 43522 | 492 | - | | | | | | | hypothetical protein |
| ORF98 | 43519 | 43902 | 384 | - | | | | | | | hypothetical protein |
| ORF99 | 44003 | 44392 | 390 | - | | | | | | | hypothetical protein |
| ORF100 | 44523 | 44957 | 435 | - | | | | | | | hypothetical protein |
| ORF101 | 44960 | 45754 | 795 | - | dUTP diphosphatase Rummeliibacillus sp. TYF005 | WP_124217124 | 3 | 45 | 63 | 5.46E-32 | deoxyuridine 5-triphosphate nucleotidohydrolase |
| ORF102 | 45769 | 46302 | 534 | - | hypothetical protein T548_0137 Lactococcus phage phiL47 | YP_009007015 | 7 | 28 | 47 | 3.93E-05 | hypothetical protein |
| ORF103 | 46458 | 47102 | 645 | + | terminase small subunit Enterococcus phage EF-P29 | APU00269 | 3 | 54 | 73 | 9.24E-62 | terminase small subunit |
| ORF104 | 47099 | 48376 | 1278 | + | terminase large subunit Enterococcus phage vB_EfaS_Ef7.1 | QBZ69408 | 0 | 65 | 80 | 0.00E+00 | terminase large subunit |
| ORF105 | 48389 | 49963 | 1575 | + | phage portal protein Streptococcus phage SPQS1 | YP_008320482 | 3 | 61 | 76 | 0.00E+00 | portal protein |
| ORF106 | 49976 | 50722 | 747 | + | hypothetical protein Enterococcus phage vB_EfaS_Ef2.2 | QBZ69220 | 4 | 33 | 51 | 1.74E-28 | capsid head morphogenesis protein |
| ORF107 | 50824 | 51468 | 645 | + | DUF4355 domain-containing protein partial *Oceanospirillum linum* | WP_139363748 | 0 | 71 | 87 | 3.01E-64 | DUF4355 domain-containing protein |
| ORF108 | 51506 | 52354 | 849 | + | putative major head protein Enterococcus phage VD13 | YP_009036381 | 1 | 50 | 72 | 2.61E-92 | major capsid head protein |

| | | | | | | | | | | | |
|---|---|---|---|---|---|---|---|---|---|---|---|
| ORF109 | 52357 | 52875 | 519 | + | major tail protein Staphylococcus phage vB_SauS_IMEP5 | ANM47024 | 2 | 55 | 71 | 2.58E-38 | major tail protein |
| ORF110 | 52946 | 53329 | 384 | + | head-tail connector family protein Enterococcus phage EF-P10 | AQT27721 | 2 | 52 | 75 | 6.58E-39 | head-tail connector family protein |
| ORF111 | 53341 | 53706 | 366 | + | hypothetical protein Enterococcus phage vB_EfaS_IME198 | YP_009218888 | 4 | 50 | 63 | 4.81E-29 | hypothetical protein |
| ORF112 | 53694 | 54071 | 378 | + | hypothetical protein A958_gp18 Enterococcus phage BC611 | YP_006488745 | 2 | 35 | 54 | 7.15E-16 | hypothetical protein |
| ORF113 | 54082 | 54567 | 486 | + | hypothetical protein Enterococcus phage vB_EfaS_Ef2.2 | QBZ69227 | 0 | 64 | 77 | 1.43E-58 | tail protein |
| ORF114 | 54590 | 55291 | 702 | + | hypothetical protein Enterococcus phage vB_EfaS_Ef2.2 | QBZ69228 | 0 | 64 | 79 | 4.12E-103 | Major tail-protein |
| ORF115 | 55417 | 55860 | 444 | + | Ig domain-containing protein Staphylococcus equorum | WP_069832504 | 7 | 51 | 68 | 1.39E-33 | major tail protein |
| ORF116 | 55981 | 56415 | 435 | + | hypothetical protein Enterococcus phage vB_EfaS_IME198 | YP_009218892 | 1 | 58 | 76 | 5.45E-51 | hypothetical protein |
| ORF117 | 56471 | 56671 | 201 | + | | | | | | | hypothetical protein |
| ORF118 | 56688 | 59813 | 3126 | + | minor capsid protein Enterococcus phage IMEEF1 | YP_009603974 | 13 | 42 | 58 | 0.00E+00 | tail tape measure protein |
| ORF119 | 59828 | 63598 | 3771 | + | tail fiber protein Enterococcus phage Entf1 | QDB70491 | 6 | 36 | 57 | 7.92E-89 | tail fiber protein |
| ORF120 | 63610 | 70560 | 6951 | + | BppU family phage baseplate upper protein Enterococcus faecium | WP_016628906 | 1 | 87 | 92 | 0.00E+00 | tail fiber protein |
| ORF121 | 70560 | 71102 | 543 | + | | | | | | | hypothetical protein |
| ORF122 | 71222 | 71554 | 333 | + | holin Enterococcus phage vB_EfaS-DELF1 | BBQ04297 | 0 | 54 | 78 | 5.88E-26 | holin |
| ORF123 | 71568 | 71843 | 276 | + | phage holin Enterococcus faecium | WP_086319065 | 0 | 67 | 85 | 2.79E-39 | holin |

**Table S1B. *Enterococcus faecium* phage 9183 genome organization and features**

**Phage 9183 genome annotation**

| Feature ID | Gene Start | Gene Stop | Length | Strand | blastp Best-Hit cut-off (0.001) Number | Best-Hit Accession | % Gaps | % Identities | % Positives | E-value | Predicted Function |
|---|---|---|---|---|---|---|---|---|---|---|---|
| ORF1 | 313 | 1884 | 1572 | + | AAA family ATPase Pediococcus pentosaceus | WP_055126681 | 10 | 30 | 49 | 3.60E-51 | recombinase recD CDS |

| | | | | | | | | | | | |
|---|---|---|---|---|---|---|---|---|---|---|---|
| ORF2 | 1982 | 3919 | 1938 | + | hypothetical protein Enterococcus phage VFW | SCZ83951 | 0 | 71 | 84 | 0.00E+00 | recD-like DNA helicase CDS |
| ORF3 | 3912 | 4460 | 549 | + | None | | | | | | hypothetical protein |
| ORF4 | 4596 | 4925 | 330 | + | hypothetical protein Enterococcus phage VPE25 | SCO93385 | 0 | 57 | 82 | 3.72E-37 | hypothetical protein CDS |
| ORF5 | 4922 | 5386 | 465 | + | hypothetical protein Enterococcus phage VPE25 | SCO93386 | 0 | 65 | 83 | 9.58E-64 | hypothetical protein CDS |
| ORF6 | 5386 | 5790 | 405 | + | None | | | | | | hypothetical protein CDS |
| ORF7 | 5787 | 7073 | 1287 | + | DNA ligase phage-associated Enterococcus phage VPE25 | SCO93389 | 0 | 70 | 83 | 0.00E+00 | DNA ligase, phage-associated CDS |
| ORF8 | 7070 | 7258 | 189 | + | None | | | | | | hypothetical protein CDS |
| ORF9 | 7354 | 7545 | 192 | + | hypothetical protein Enterococcus phage VPE25 | SCO93390 | 0 | 57 | 74 | 6.76E-15 | hypothetical protein CDS |
| ORF10 | 7532 | 7747 | 216 | + | | | | | | | hypothetical protein CDS |
| ORF11 | 7728 | 7919 | 192 | + | | | | | | | hypothetical protein CDS |
| ORF12 | 8088 | 8294 | 207 | + | | | | | | | hypothetical protein CDS |
| ORF13 | 8446 | 8703 | 258 | + | | | | | | | hypothetical protein CDS |
| ORF14 | 8843 | 9364 | 522 | + | Lysine decarboxylase family Enterococcus phage VFW | SCZ83963 | 1 | 61 | 80 | 1.31E-67 | Lysine decarboxylase family CDS |
| ORF15 | 9379 | 10026 | 648 | + | Enterococcus phage VPE25 | SCO93397 | 6 | 59 | 69 | 4.10E-79 | Purine trans deoxyribosylase Nucleoside deoxyribosyltransferase-I CDS |
| ORF16 | 10143 | 10991 | 849 | + | Deoxyguanosine kinase Enterococcus phage VFW | SCZ83966 | 0 | 52 | 71 | 1.06E-93 | Deoxyguanosine kinase CDS |
| ORF17 | 11015 | 11779 | 765 | + | NrdR-regulated deoxyribonucleotide transporter PnuC-like Enterococcus phage VPE25 | SCO93399 | 0 | 87 | 94 | 5.69E-156 | NrdR-regulated deoxyribonucleotide transporter PnuC-like CDS |
| ORF18 | 11865 | 12071 | 207 | + | glutaredoxin-like protein NrdH Enterococcus canis | WP_082703267 | 3 | 46 | 62 | 5.38E-10 | NrdH-like glutaredoxin CDS |
| ORF19 | 12146 | 13189 | 1044 | + | hypothetical protein Enterococcus phage VPE25 | SCO93402 | 1 | 62 | 79 | 7.90E-154 | DNA response regulator CDS |
| ORF20 | 13278 | 14126 | 849 | + | hypothetical protein Enterococcus phage VPE25 | SCO93403 | 1 | 61 | 78 | 2.58E-118 | HNH endonuclease CDS |

| | | | | | | | | | | | |
|---|---|---|---|---|---|---|---|---|---|---|---|
| ORF21 | 14119 | 14712 | 594 | + | | | | | | | hypothetical protein CDS |
| ORF22 | 14726 | 14926 | 201 | + | | | | | | | hypothetical protein CDS |
| ORF23 | 15020 | 15982 | 963 | + | hypothetical protein Enterococcus phage VFW | SCZ83974 | 2 | 73 | 85 | 5.99E-167 | single strand binding protein CDS |
| ORF24 | 16056 | 16523 | 468 | + | hypothetical protein Enterococcus phage VFW | SCZ83975 | 3 | 24 | 54 | 1.66E-05 | hypothetical protein CDS |
| ORF25 | 16525 | 16941 | 417 | + | | | | | | | hypothetical protein CDS |
| ORF26 | 17016 | 18083 | 1068 | + | hypothetical protein Enterococcus phage VPE25 | SCO93409 | 0 | 85 | 92 | 0.00E+00 | ATP GTP binding protein CDS |
| ORF27 | 18141 | 18830 | 690 | + | | | | | | | hypothetical protein CDS |
| ORF28 | 18820 | 19020 | 201 | + | | | | | | | hypothetical protein CDS |
| ORF29 | 19017 | 19757 | 741 | + | hypothetical protein Enterococcus phage VPE25 | SCO93413 | 1 | 51 | 69 | 1.65E-79 | hypothetical protein CDS |
| ORF30 | 19837 | 20013 | 177 | + | hypothetical protein SSG_02354 Enterococcus faecium EnGen0190 | EOH43274 | 0 | 82 | 91 | 7.54E-28 | hypothetical protein CDS |
| ORF31 | 20023 | 20259 | 237 | + | hypothetical protein Peptostreptococcus sp. MV1 | WP_052045454 | 10 | 55 | 71 | 7.14E-21 | hypothetical protein CDS |
| ORF32 | 20351 | 20854 | 504 | + | phage protein Enterococcus phage VPE25 | SCO93419 | 0 | 70 | 83 | 7.69E-82 | hypothetical protein CDS |
| ORF33 | 20851 | 22431 | 1581 | + | Replicative DNA helicase DnaB Enterococcus phage VPE25 | SCO93420 | 0 | 82 | 93 | 0.00E+00 | Replicative DNA helicase DnaB CDS |
| ORF34 | 22444 | 23493 | 1050 | + | Phage toprim domain containing protein YorJ B.subtilis homolog Enterococcus phage VPE25 | SCO93421 | 0 | 70 | 86 | 0.00E+00 | DNA primase CDS |
| ORF35 | 23505 | 25328 | 1824 | + | Single-stranded-DNA-specific exonuclease RecJ Enterococcus phage VPE25 | SCO93422 | 2 | 60 | 76 | 0.00E+00 | Single-stranded-DNA-specific exonuclease RecJ CDS |
| ORF36 | 25413 | 25988 | 576 | + | hypothetical protein Enterococcus phage VPE25 | SCO93423 | 6 | 36 | 59 | 1.01E-28 | hypothetical protein CDS |
| ORF37 | 26176 | 26355 | 180 | + | | | | | | | hypothetical protein CDS |
| ORF38 | 26352 | 26690 | 339 | + | | | | | | | hypothetical protein CDS |
| ORF39 | 26695 | 27207 | 513 | + | | | | | | | hypothetical protein CDS |
| ORF40 | 27245 | 27394 | 150 | + | | | | | | | hypothetical protein CDS |

| | | | | | | | | | | | |
|---|---|---|---|---|---|---|---|---|---|---|---|
| ORF41 | 27472 | 27702 | 231 | + | | | | | | | hypothetical protein CDS |
| ORF42 | 27727 | 27870 | 144 | + | hypothetical protein Enterococcus phage VPE25 | SCO93428 | 0 | 67 | 87 | 4.92E-14 | hypothetical protein CDS |
| ORF43 | 28004 | 28522 | 519 | + | hypothetical protein Enterococcus phage VPE25 | SCO93430 | 7 | 47 | 64 | 3.36E-35 | DTMP kinase, thymidylate kinase CDS |
| ORF44 | 28523 | 29356 | 834 | + | Thymidylate synthase Enterococcus phage VPE25 | SCO93431 | 1 | 55 | 73 | 5.74E-103 | Thymidylate synthase CDS |
| ORF45 | 29340 | 29756 | 417 | + | Deoxyuridine 5-triphosphate nucleotidohydrolase Enterococcus phage VPE25 | SCO93432 | 0 | 78 | 90 | 1.59E-75 | Deoxyuridine 5-triphosphate nucleotidohydrolase CDS |
| ORF46 | 29756 | 30100 | 345 | + | HAD hydrolase family protein Longirhabdus pacifica | WP_128894611 | 3 | 50 | 68 | 5.41E-31 | HAD hydrolase family protein CDS |
| ORF47 | 30097 | 31731 | 1635 | + | hypothetical protein A2138_23700 Deltaproteobacteria bacterium RBG_16_71_12 | OGQ17127 | 7 | 33 | 52 | 4.86E-83 | nucleoside-diphosphate-sugar pyrophosphorylase-like protein CDS |
| ORF48 | 31736 | 32956 | 1221 | + | hypothetical protein CBD54_002560 Alphaproteobacteria bacterium TMED194 | RPH07865 | 6 | 30 | 50 | 5.41E-45 | hypothetical protein CDS |
| ORF49 | 33043 | 33567 | 525 | + | Ribonuclease HI Enterococcus phage VPE25 | SCO93434 | 0 | 87 | 93 | 1.14E-107 | Ribonuclease HI CDS |
| ORF50 | 33564 | 34535 | 972 | + | Nucleotide sugar synthetase-like protein Enterococcus phage VPE25; | SCO93435 | 1 | 48 | 68 | 5.98E-98 | Nucleotide sugar synthetase-like protein CDS |
| ORF51 | 34525 | 34779 | 255 | + | | | | | | | hypothetical protein CDS |
| ORF52 | 34772 | 35380 | 609 | + | hypothetical protein Enterococcus phage VPE25 | SCO93436 | 1 | 52 | 68 | 1.03E-19 | Thymidine kinase CDS |
| ORF53 | 35460 | 39314 | 3855 | + | DNA polymerase III alpha subunit Enterococcus phage VFW | SCZ84010 | 0 | 78 | 89 | 0.00E+00 | DNA polymerase III alpha subunit CDS |
| ORF54 | 39381 | 39644 | 264 | + | hypothetical protein Enterococcus phage VPE25 | SCO93444 | 1 | 35 | 57 | 9.42E-08 | hypothetical protein CDS |
| ORF55 | 39641 | 39919 | 279 | + | | | | | | | hypothetical protein CDS |
| ORF56 | 40197 | 40610 | 414 | - | | | | | | | hypothetical protein CDS |

| | | | | | | | | | | | |
|---|---|---|---|---|---|---|---|---|---|---|---|
| ORF57 | 40850 | 41422 | 573 | + | hypothetical protein Enterococcus phage VFW | SCZ84015 | 9 | 32 | 50 | 2.72E-07 | hypothetical protein CDS |
| ORF58 | 41412 | 41699 | 288 | + | | | | | | | hypothetical protein CDS |
| ORF59 | 41744 | 41962 | 219 | + | hypothetical protein Bacillus cereus | WP_073526565 | 3 | 70 | 80 | 3.62E-27 | DUF2829 domain-containing protein CDS |
| ORF60 | 41962 | 42705 | 744 | + | Deoxyuridine 5-triphosphate nucleotidohydrolase Enterococcus phage VPE25 | SCO93451 | 3 | 58 | 72 | 1.68E-83 | Deoxyuridine 5-triphosphate nucleotidohydrolase CDS |
| ORF61 | 42706 | 43005 | 300 | + | | | | | | | hypothetical protein CDS |
| ORF62 | 42995 | 43594 | 600 | + | Guanylate kinase Enterococcus phage VPE25 | SCO93453 | 5 | 41 | 65 | 5.56E-40 | Guanylate kinase CDS |
| ORF63 | 43595 | 44179 | 585 | + | non-essential protein Enterococcus phage VPE25 | SCO93454 | 0 | 75 | 90 | 5.21E-103 | RusA family crossover junction endodeoxyribonuclease CDS |
| ORF64 | 44148 | 44531 | 384 | + | | | | | | | hypothetical protein CDS |
| ORF65 | 44946 | 45575 | 630 | + | hypothetical protein Enterococcus phage VPE25 | SCO93459 | 1 | 53 | 74 | 3.72E-73 | sigma-70 family RNA polymerase sigma factor CDS |
| ORF66 | 45677 | 47614 | 1938 | + | DNA gyrase subunit B Enterococcus phage VPE25 | SCO93463 | | | | 0.00E+00 | Topoisomerase IV subunit B CDS |
| ORF67 | 47709 | 47966 | 258 | + | | | | | | | hypothetical protein CDS |
| ORF68 | 47959 | 49962 | 2004 | + | DNA gyrase subunit A Enterococcus phage VFW | SCZ84033 | 2 | 60 | 77 | 0.00E+00 | DNA topoisomerase IV subunit A CDS |
| ORF69 | 50114 | 50290 | 177 | + | | | | | | | hypothetical protein CDS |
| ORF70 | 50393 | 50623 | 231 | + | | | | | | | hypothetical protein CDS |
| ORF71 | 50820 | 51230 | 411 | + | | | | | | | hypothetical protein CDS |
| ORF72 | 51220 | 51531 | 312 | + | | | | | | | hypothetical protein CDS |
| ORF73 | 51528 | 52100 | 573 | + | | | | | | | hypothetical protein CDS |
| ORF74 | 52217 | 52405 | 189 | + | | | | | | | hypothetical protein CDS |
| ORF75 | 52431 | 52904 | 474 | + | | | | | | | hypothetical protein CDS |
| ORF76 | 52921 | 53103 | 183 | + | | | | | | | hypothetical protein CDS |
| ORF77 | 53139 | 54116 | 978 | - | prophage LambdaBa02 site-specific recombinase phage integrase family Enterococcus phage VFW | SCZ84050 | 0 | 86 | 92 | 0.00E+00 | site-specific recombinase phage integrase family CDS |

| | | | | | | | | | | | |
|---|---|---|---|---|---|---|---|---|---|---|---|
| ORF78 | 54176 | 55339 | 1164 | - | N-acetylmuramoyl-L-alanine amidase Enterococcus phage VPE25 | SCO93486 | 11 | 48 | 59 | 1.91E-102 | N-acetylmuramoyl-L-alanine amidase CDS |
| ORF79 | 55451 | 55810 | 360 | - | hypothetical protein Enterococcus phage VPE25 | SCO93487 | 0 | 67 | 84 | 8.04E-52 | holin CDS |
| ORF80 | 55810 | 56181 | 372 | - | hypothetical protein Enterococcus phage VPE25 | SCO93488 | 0 | 72 | 84 | 8.92E-40 | hypothetical protein CDS |
| ORF81 | 56159 | 56557 | 399 | - | hypothetical protein Enterococcus phage VPE25 | SCO93489 | 0 | 61 | 83 | 8.36E-57 | hypothetical protein CDS |
| ORF82 | 56564 | 56698 | 135 | - | | | | | | | hypothetical protein CDS |
| ORF83 | 56701 | 57186 | 486 | - | hypothetical protein Enterococcus faecalis | WP_010774487 | 12 | 42 | 56 | 1.77E-18 | hypothetical protein CDS |
| ORF84 | 57207 | 60089 | 2883 | - | BppU family phage baseplate upper protein Enterococcus faecium | WP_104807894 | 4 | 53 | 68 | 0.00E+00 | BppU family phage baseplate upper protein CDS |
| ORF85 | 60103 | 64134 | 4032 | - | hypothetical protein Enterococcus faecium | WP_142972363 | 6 | 48 | 59 | 4.22E-150 | minor tail protein CDS |
| ORF86 | 64164 | 66476 | 2313 | - | hypothetical protein Enterococcus faecalis | WP_057086899 | 8 | 48 | 63 | 0.00E+00 | Phage endopeptidase, tail-spike protein CDS |
| ORF87 | 66473 | 67264 | 792 | - | hypothetical protein Enterococcus phage VPE25 | SCO93493 | 0 | 62 | 78 | 9.10E-121 | Phage tail protein CDS |
| ORF88 | 67277 | 71119 | 3843 | - | Phage tail length tape-measure protein Enterococcus phage VPE25 | SCO93494 | 2 | 54 | 72 | 0.00E+00 | Phage tail length tape-measure protein CDS |
| ORF89 | 71373 | 71705 | 333 | - | hypothetical protein Enterococcus phage VPE25 | SCO93496 | 0 | 76 | 90 | 1.98E-53 | hypothetical protein CDS |
| ORF90 | 71872 | 72480 | 609 | - | Phage major tail protein phi13 Enterococcus phage VFW | SCZ84062 | 0 | 89 | 92 | 2.55E-107 | Phage major tail protein, phage 13 family CDS |
| ORF91 | 72504 | 72881 | 378 | - | hypothetical protein Enterococcus phage VPE25 | SCO93498 | 0 | 84 | 90 | 2.68E-69 | hypothetical protein CDS |
| ORF92 | 72884 | 73333 | 450 | - | hypothetical protein Enterococcus phage VPE25 | SCO93499 | 0 | 71 | 80 | 4.30E-70 | Phage head-tail joining protein, HK97 gp10 family CDS |
| ORF93 | 73326 | 73679 | 354 | - | hypothetical protein Enterococcus phage VPE25 | SCO93500 | 0 | 78 | 92 | 1.49E-60 | Phage head-tail adaptor protein CDS |
| ORF94 | 73683 | 74060 | 378 | - | hypothetical protein Enterococcus phage VPE25 | SCO93501 | 0 | 72 | 85 | 9.18E-61 | Phage head-tail connector protein CDS |

| | | | | | | | | | | | |
|---|---|---|---|---|---|---|---|---|---|---|---|
| ORF95 | 74200 | 75069 | 870 | - | prophage pi2 protein 34 Enterococcus phage VPE25 | SCO93502 | 0 | 68 | 81 | 6.18E-141 | Prophage pi2 protein 34 CDS |
| ORF96 | 75270 | 76475 | 1206 | - | hypothetical protein Enterococcus phage VFW | SCZ84068 | 1 | 79 | 87 | 0.00E+00 | Phage major capsid protein CDS |
| ORF97 | 76465 | 77673 | 1209 | - | hypothetical protein Enterococcus phage VFW | SCZ84069 | 3 | 61 | 71 | 3.51E-150 | Phage prohead protease, HK97 family CDS |
| ORF98 | 77688 | 78965 | 1278 | - | hypothetical protein Enterococcus phage VPE25 | SCO93505 | 0 | 77 | 89 | 0.00E+00 | Phage portal protein CDS |
| ORF99 | 78978 | 80684 | 1707 | - | hypothetical protein Enterococcus phage VFW | SCZ84071 | 0 | 85 | 92 | 0.00E+00 | Phage terminase large subunit CDS |
| ORF100 | 81115 | 81606 | 492 | - | Phage-related protein Enterococcus phage VPE25 | SCO93509 | 0 | 86 | 96 | 4.62E-101 | Phage terminase small subunit, P27 family CDS |
| ORF101 | 81609 | 82220 | 612 | - | hypothetical protein Bacillus | WP_063263132 | 14 | 38 | 51 | 5.16E-12 | GIY-YIG homing endonuclease CDS |
| ORF102 | 82217 | 82636 | 420 | - | hypothetical protein Enterococcus phage VPE25 | SCO93510 | 3 | 59 | 78 | 1.57E-54 | HNH endonuclease CDS |
| ORF103 | 82611 | 82769 | 159 | - | | | | | | | hypothetical protein CDS |
| ORF104 | 82955 | 83212 | 258 | + | | | | | | | hypothetical protein CDS |
| ORF105 | 83214 | 83657 | 444 | + | | | | | | | hypothetical protein CDS |
| ORF106 | 83654 | 84031 | 378 | + | | | | | | | hypothetical protein CDS |
| ORF107 | 84028 | 84219 | 192 | + | | | | | | | hypothetical protein CDS |
| ORF108 | 84216 | 84497 | 282 | + | | | | | | | hypothetical protein CDS |
| ORF109 | 84526 | 85398 | 873 | + | hypothetical protein Enterococcus phage VPE25 | SCO93512 | 3 | 42 | 62 | 2.53E-63 | hypothetical protein CDS |

**Table S1C. *Enterococcus faecium* phage 9184 genome organization and features**

**Phage 9184 genome annotation**

| Feature ID | Gene Start | Gene Stop | Length | Strand | blastp Best-Hit cut-off (0.001) Number | Best-Hit Accession | % Gaps | % Identities | % Positives | E-value | Predicted Function |
|---|---|---|---|---|---|---|---|---|---|---|---|
| ORF1 | 260 | 439 | 180 | + | hypothetical protein Enterococcus phage vB_EfaS-DELF1 | BBQ04339 | 0 | 78 | 85 | 2.55E-25 | hypothetical protein |
| ORF2 | 444 | 902 | 459 | + | terminase small subunit Enterococcus phage IME-EFm5 | YP_009200920 | 1 | 78 | 86 | 3.11E-76 | terminase small subunit |

| | | | | | | | | | | | |
|---|---|---|---|---|---|---|---|---|---|---|---|
| ORF3 | 1545 | 3323 | 1779 | + | terminase large subunit Enterococcus phage IME-EFm1 | YP_009042651 | 0 | 94 | 98 | 0 | terminase large subunit |
| ORF4 | 3391 | 3567 | 177 | + | sensor histidine kinase Enterococcus phage IME-EFm1 | YP_009042652 | 0 | 97 | 98 | 2.96E-30 | sensor histidine kinase |
| ORF5 | 3571 | 4776 | 1206 | + | portal protein Enterococcus phage IME-EFm5 | YP_009200917 | 0 | 81 | 90 | 0.00E+00 | portal protein |
| ORF6 | 4784 | 5278 | 495 | + | prohead protease Enterococcus phage IME-EFm5 | YP_009200916 | 0 | 95 | 99 | 1.88E-105 | prohead protease |
| ORF7 | 5348 | 6595 | 1248 | + | capsid protein Enterococcus phage Nonaheksakonda | AZS06457 | 6 | 61 | 76 | 3.10E-162 | capsid protein |
| ORF8 | 6675 | 7013 | 339 | + | head-tail joining protein Enterococcus phage IME-EFm5 | YP_009200913 | 0 | 79 | 91 | 2.64E-61 | head-tail connector protein |
| ORF9 | 6943 | 7278 | 336 | + | head-tail adaptor protein Enterococcus phage IME-EFm1 | YP_009042657 | 0 | 95 | 96 | 7.59E-71 | head-tail adaptor protein |
| ORF10 | 7280 | 7651 | 372 | + | head-tail joining protein Enterococcus phage IME-EFm5 | YP_009200912 | 0 | 89 | 93 | 4.06E-74 | head-tail joining protein |
| ORF11 | 7651 | 8016 | 366 | + | head-tail joining protein Enterococcus phage IME-EFm5 | YP_009200911 | 0 | 87 | 94 | 1.97E-71 | head-tail joining protein |
| ORF12 | 8089 | 8649 | 561 | + | major tail protein Enterococcus phage IME-EFm1 | YP_009042660 | 1 | 87 | 95 | 1.35E-113 | major tail protein |
| ORF13 | 8708 | 9088 | 381 | + | putative tail tape measure chaperone protein Enterococcus phage IME-EFm5 | YP_009200909 | 0 | 81 | 93 | 2.90E-56 | tail tape measure protein |
| ORF14 | 9121 | 9318 | 198 | + | tail tape measure chaperone protein Enterococcus phage IME-EFm1 | YP_009042662 | 0 | 89 | 97 | 2.11E-35 | tail tape measure chaperone protein |
| ORF15 | 9383 | 13873 | 4491 | + | transglycosylase SLT domain-containing protein Enterococcus durans | WP_119219106 | 1 | 75 | 85 | 0.00E+00 | tail length tape-measure protein |
| ORF16 | 13945 | 14967 | 1023 | + | minor tail protein Enterococcus phage IME-EFm5 | YP_009200906 | 0 | 92 | 97 | 0.00E+00 | minor tail protein |
| ORF17 | 14954 | 17174 | 240 | + | minor tail protein Enterococcus phage IME-EFm5 | YP_009200906 | 0 | 71 | 81 | 0.00E+00 | BppU family phage baseplate upper protein |
| ORF18 | 17247 | 18650 | 1404 | + | minor tail protein Enterococcus phage IME-EFm5 | YP_009200905 | 1 | 75 | 86 | 0.00E+00 | minor tail protein |

| | | | | | | | | | | | |
|---|---|---|---|---|---|---|---|---|---|---|---|
| ORF19 | 18665 | 19651 | 987 | + | tail assembly protein Enterococcus phage IME-EFm5 | YP_009200904 | 2 | 58 | 72 | 1.74E-110 | tail assembly protein |
| ORF20 | 19830 | 20108 | 279 | + | holin Enterococcus phage IME-EFm1 | YP_009042670 | 0 | 97 | 99 | 1.24E-57 | holin |
| ORF21 | 20122 | 20403 | 282 | + | holin Enterococcus phage IME-EFm5 | YP_009200902 | 0 | 98 | 99 | 2.26E-58 | holin |
| ORF22 | 20420 | 21445 | 1026 | + | N-acetylmuramoyl-L-alanine amidase Enterococcus phage IME-EFm5 | YP_009200901 | 0 | 95 | 96 | 0.00E+00 | N-acetylmuramoyl-L-alanine amidase |
| ORF23 | 21524 | 22219 | 696 | - | hypothetical protein EFm5_30 Enterococcus phage IME-EFm5 | YP_009200900 | 0 | 90 | 95 | 3.40E-150 | Deoxyguanosine kinase |
| ORF24 | 22517 | 23347 | 831 | - | hypothetical protein phiSHEF2_24 Enterococcus phage phiSHEF2 | YP_009613304 | 0 | 63 | 80 | 8.82E-24 | DNA polymerase |
| ORF25 | 23314 | 23958 | 645 | - | hypothetical protein Streptococcus pyogenes | WP_136291116 | 6 | 33 | 57 | 5.20E-27 | ABC transporter ATP-binding protein |
| ORF26 | 24003 | 26033 | 2031 | - | DNA polymerase Enterococcus phage vB_EfaS-DELF1 | BBQ04302 | 1 | 62 | 77 | 0.00E+00 | DNA polymerase |
| ORF27 | 26142 | 26348 | 207 | - | hypothetical protein Enterococcus phage vB_EfaS-DELF1 | BBQ04303 | 0 | 65 | 78 | 1.57E-24 | hypothetical protein |
| ORF28 | 26455 | 27240 | 786 | - | hypothetical protein IME_032 Enterococcus phage IME-EFm1 | YP_009042680 | 5 | 50 | 71 | 3.54E-71 | hypothetical protein |
| ORF29 | 27299 | 27517 | 219 | - | | | | | | | hypothetical protein |
| ORF30 | 27514 | 28323 | 810 | - | hypothetical protein EFm5_22 Enterococcus phage IME-EFm5 | YP_009200892 | 0 | 78 | 87 | 1.61E-146 | Protein of unknown function DUF1351 |
| ORF31 | 28313 | 28483 | 171 | - | hypothetical protein IME_035 Enterococcus phage IME-EFm1 | YP_009042683 | 0 | 89 | 93 | 9.29E-26 | hypothetical protein |
| ORF32 | 28480 | 28701 | 222 | - | | | | | | | hypothetical protein |
| ORF33 | 28701 | 28928 | 228 | - | hypothetical protein EFm5_20 Enterococcus phage IME-EFm5 | YP_009200890 | 0 | 92 | 97 | 1.47E-42 | hypothetical protein |
| ORF34 | 29107 | 29436 | 330 | - | hypothetical protein IME_030 Enterococcus phage IME-EFm1 | YP_009042678 | 3 | 51 | 72 | 9.10E-32 | hypothetical protein |
| ORF35 | 29474 | 30235 | 762 | - | metallo-beta-lactamase domain protein Enterococcus phage vB_EfaS-DELF1 | BBQ04313 | 0 | 74 | 86 | 1.26E-131 | Metallo-beta-lactamase domain protein |

| | | | | | | | | | | | |
|---|---|---|---|---|---|---|---|---|---|---|---|
| ORF36 | 30210 | 30680 | 471 | - | HNH homing endonuclease-like protein Enterococcus phage IME-EFm5 | YP_009200887 | 0 | 79 | 92 | 1.55E-90 | HNH homing endonuclease-like protein |
| ORF37 | 30670 | 31161 | 492 | - | hypothetical protein IME_041 Enterococcus phage IME-EFm1 | YP_009042689 | 2 | 61 | 73 | 1.33E-55 | hypothetical protein |
| ORF38 | 31301 | 31501 | 201 | - | hypothetical protein EFm5_15 Enterococcus phage IME-EFm5 | YP_009200885 | 0 | 86 | 97 | 4.58E-36 | hypothetical protein |
| ORF39 | 31501 | 31680 | 180 | - | hypothetical protein EFm5_14 Enterococcus phage IME-EFm5 | YP_009200884 | 0 | 66 | 76 | 6.47E-18 | hypothetical protein |
| ORF40 | 31677 | 32417 | 741 | - | DNA replication protein Enterococcus phage IME-EFm5 | YP_009200883 | 0 | 82 | 91 | 2.59E-150 | bifunctional DNA primase-polymerase |
| ORF41 | 32414 | 33154 | 741 | - | adenine-specific methyltransferase Enterococcus phage vB_EfaS_Ef5.4 | QBZ69846 | 2 | 63 | 79 | 7.63E-108 | DNA methylase |
| ORF42 | 33210 | 33368 | 159 | - | hypothetical protein Enterococcus phage vB_EfaS-DELF1 | BBQ04319 | 7 | 71 | 82 | 1.93E-17 | hypothetical protein |
| ORF43 | 33365 | 34642 | 1278 | - | helicase Enterococcus phage IME-EFm5 | YP_009200880 | 0 | 90 | 95 | 0.00E+00 | DNA helicase |
| ORF44 | 34639 | 34914 | 276 | - | hypothetical protein EFm5_08 Enterococcus phage IME-EFm5 | YP_009200878 | 2 | 77 | 90 | 1.21E-44 | putative RNA-binding protein |
| ORF45 | 34914 | 35030 | 117 | - | hypothetical protein EFm5_07 Enterococcus phage IME-EFm5 | YP_009200877 | 0 | 61 | 79 | 4.99E-07 | hypothetical protein |
| ORF46 | 35027 | 35194 | 168 | - | hypothetical protein EFm5_06 Enterococcus phage IME-EFm5 | YP_009200876 | 0 | 73 | 85 | 3.36E-20 | hypothetical protein |
| ORF47 | 35187 | 35537 | 351 | - | endonuclease Enterococcus phage IME-EFm5 | YP_009200875 | 0 | 93 | 98 | 5.11E-74 | Endonuclease |
| ORF48 | 35534 | 35749 | 216 | - | | | | | | | hypothetical protein |
| ORF49 | 35746 | 35973 | 228 | - | hypothetical protein EFm5_03 Enterococcus phage IME-EFm5 | YP_009200873 | 1 | 66 | 83 | 9.04E-21 | hypothetical protein |
| ORF50 | 35963 | 36055 | 93 | - | | | | | | | hypothetical protein |
| ORF51 | 36055 | 36252 | 198 | - | | | | | | | hypothetical protein |
| ORF52 | 36249 | 36464 | 216 | - | | | | | | | hypothetical protein |
| ORF53 | 36461 | 36658 | 198 | - | | | | | | | hypothetical protein |

| | | | | | | | | | | | |
|---|---|---|---|---|---|---|---|---|---|---|---|
| ORF54 | 36735 | 38357 | 1623 | - | DNA primase Enterococcus phage vB_EfaS-DELF1 | BBQ04327 | 9 | 31 | 51 | 3.86E-50 | DNA primase |
| ORF55 | 38404 | 38607 | 204 | - | hypothetical protein Enterococcus phage vB_EfaS-DELF1 | BBQ04330 | 6 | 60 | 72 | 1.27E-12 | hypothetical protein |
| ORF56 | 38678 | 38860 | 183 | - | hypothetical protein EFm5_68 Enterococcus phage IME-EFm5 | YP_009200938 | 0 | 82 | 95 | 1.09E-28 | putative swarming motility protein |
| ORF57 | 38853 | 39032 | 180 | - | hypothetical protein EFm5_67 Enterococcus phage IME-EFm5 | YP_009200937 | 0 | 64 | 83 | 3.69E-19 | hypothetical protein |
| ORF58 | 39032 | 39535 | 504 | - | hypothetical protein EFm5_66 Enterococcus phage IME-EFm5 | YP_009200936 | 0 | 75 | 86 | 4.21E-28 | tail length tape-measure protein |
| ORF59 | 39601 | 39831 | 231 | - | hypothetical protein CUN38_04900 Enterococcus faecium | PQC93482 | 0 | 80 | 89 | 2.27E-34 | hypothetical protein |
| ORF60 | 39831 | 40040 | 210 | - | | | | | | | hypothetical protein |
| ORF61 | 40056 | 40238 | 183 | - | hypothetical protein Enterococcus faecalis | WP_033659461 | 0 | 58 | 77 | 5.57E-14 | hypothetical protein |
| ORF62 | 40251 | 40613 | 363 | - | hypothetical protein IME_057 Enterococcus phage IME-EFm1 | YP_009042705 | 0 | 71 | 87 | 4.63E-42 | hypothetical protein |
| ORF63 | 40625 | 40951 | 327 | - | hypothetical protein IME_059 Enterococcus phage IME-EFm1 | YP_009042707 | 0 | 82 | 91 | 1.40E-58 | DUF1140 protein |
| ORF64 | 40945 | 41178 | 234 | - | hypothetical protein IME_060 Enterococcus phage IME-EFm1 | YP_009042708 | 1 | 60 | 81 | 5.08E-24 | hypothetical protein |
| ORF65 | 41234 | 41467 | 234 | - | | | | | | | hypothetical protein |
| ORF66 | 41524 | 41826 | 303 | - | | | | | | | hypothetical protein |
| ORF67 | 42246 | 42410 | 165 | + | | | | | | | hypothetical protein |
| ORF68 | 42440 | 42664 | 225 | + | | | | | | | hypothetical protein |
| ORF69 | 42661 | 42843 | 183 | + | | | | | | | hypothetical protein |
| ORF70 | 42809 | 42988 | 180 | + | | | | | | | hypothetical protein |
| ORF71 | 42999 | 43190 | 192 | + | hypothetical protein EFm5_55 Enterococcus phage IME-EFm5 | YP_009200925 | 0 | 71 | 89 | 2.29E-26 | hypothetical protein |
| ORF72 | 43190 | 43366 | 177 | + | hypothetical protein IME_069 Enterococcus phage IME-EFm1 | YP_009042717 | 10 | 78 | 86 | 3.14E-23 | hypothetical protein |
| ORF73 | 43565 | 43942 | 378 | + | HNH endonuclease Enterococcus phage IME-EFm5 | YP_009200923 | 0 | 89 | 96 | 4.02E-76 | HNH endonuclease |
