## Supplementary Table 2 for "Lytic bacteriophages facilitate antibiotic sensitization of *Enterococcus faecium*"

**Table S2A. Spontaneous, non-synonymous mutations in secreted antigen A (SagA) gene promote phage 9181 resistance**

| Phage 9181 Resistant Mutant | Com12 Contig Number | Contig Position | Variation Type | Variation Frequency (%) | Fold Coverage | Gene locus: SNP/Indel | AA change | Putative function |
|---|---|---|---|---|---|---|---|---|
| 81R3 | NZ_GG670310 | 172959 | SNP | 100 | 160 | EFVG_RS16270:313G ⇒ T | Trp433 ⇒ Leu | SagA |
| 81R4 | NZ_GG670310 | 172960 | SNP | 100 | 279 | EFVG_RS16270: 314G ⇒T | Trp433 ⇒ Cys | SagA |
| 81R5 | NZ_GG670310 | 173040 | SNP | 100 | 119 | EFVG_RS16270:394G ⇒A | Gly460 ⇒ Asp | SagA |
| 81R6 | NZ_GG670310 | 173014 | Insertion | 71.79 | 156 | EFVG_RS16270:368-370 CTT duplication | Leu insertion between Tyr451 & Leu452 | SagA |
| 81R7 | NZ_GG670306 | 96441 | SNP | 100 | 114 | EFVG_RS05165:325G ⇒ T | Gly109 ⇒ Trp | DNA topoisomerase III |
| 81R7 | NZ_GG670306 | 418448 | SNP | 100 | 115 | EFVG_RS06695: 499C ⇒ T | Pro167 ⇒ Ser | Capsule tyrosine-protein kinase (*wze*) |
| 81R8 | NZ_GG670310 | 172965 | SNP | 99.43 | 176 | EFVG_RS16270: 319G ⇒ T | Gly435 ⇒ Val | SagA |

SNP – Single Nucleotide Polymorphism; Indel – Insertion/Deletion; SagA – secreted antigen A; AA – amino acid

**Table S2B. Spontaneous, non-synonymous mutations in the *epa* locus promotes phage 9183 resistance**

| Phage 9183 Resistant Mutant | 1,141,733 Contig Number | Contig Position | Variation Type | Variation Frequency (%) | Fold Coverage | Gene locus: SNP/Indel | AA change | Putative function |
|---|---|---|---|---|---|---|---|---|
| 83R1 | NZ_GG688464 | 157195 | SNP #1 | 100 | 79 | EFSG_RS16205: 1012C ⇒ T | Arg338 ⇒ Cys | Polyprenyl Glycosylphosphotransferase (*epaR*) |
| 83R1 | NZ_GG688461 | 1077065 | SNP #2 | 100 | 6 | EFSG_RS11135: 28G ⇒ T | Ala10 ⇒ Ser | Endonuclease III (*nth*) |
| 83R2 | No mutation detected | | | | | | | |
| 83R3 | NZ_GG688464 | 156490 | Deletion | 100 | 189 | EFSG_RS16205: Deletion of 309T | Phe103 Frameshift | Polyprenyl Glycosylphosphotransferase (*epaR*) |
| 83R4 | NZ_GG688462 | 47014 | SNP | 100 | 167 | EFSG_RS12860: 741C ⇒T | Trp104 ⇒ Cys | Gluconate 5-Dehydrogenase (*gdh*) |
| 83R4 | NZ_GG688464 | 161642 | Deletion | 88.29 | 111 | EFSG_RS16230: Deletion of 197A | Asn66 Frameshift | TarS-like Glycosyltransferase (*epaX*) |
| 83R5 | NZ_GG688461 | 891434 | SNP #1 | 100 | 78 | EFSG_RS10305: 5A ⇒ G | Glu2 ⇒ Gly | General Stress Response Protein A (*gnsA*) |
| 83R5 | NZ_GG688464 | 156807 | Deletion | 95.83 | 48 | EFSG_RS16205: Deletion of 630A | Glu211 Frameshift | Polyprenyl Glycosylphosphotransferase (*epaR*) |
| 83R5 | NZ_GG688461 | 332892 | SNP #2 | 43.75 | 80 | EFSG_RS07565: 985C ⇒ T | Leu329 ⇒ Phe | SorC family transcriptional regulator (*sorC*) |
| 83R6 | NZ_GG688464 | 157113 | SNP | 100 | 75 | EFSG_RS16205: 930G ⇒ T | Met310 ⇒ Ile | Polyprenyl Glycosylphosphotransferase (*epaR*) |
| 83R7 | NZ_GG688464 | 67506 | SNP | 100 | 74 | EFSG_RS15760: 22G ⇒ T | Glu8 ⇒ Stop | D-alanine--poly(phosphoribitol) ligase subunit (*dltA*) |

| | | | | | | | | |
|---|---|---|---|---|---|---|---|---|
| 83R7 | NZ_GG688464 | 161642 | Deletion | 92.05 | 88 | EFSG_RS16230: Deletion of 197A | Asn66 Frameshift | Tar-S-like Glycosyltransferase (*epaX*) |
| 83R8 | NZ_GG688464 | 157127 | SNP | 100 | 59 | EFSG_RS16205: 944A ⇒ G | Glu315 ⇒ Gly | Polyprenyl Glycosylphosphotransferase (*epaR*) |

SNP – Single Nucleotide Polymorphism; Indel – Insertion/Deletion; AA – Amino Acid

**Table S2C. Spontaneous, non-synonymous mutations in the capsule locus and *rdd* (*efsg_rs09545*) genes promotes phage 9184 resistance**

| Phage 9184 Resistant Mutant | 1,141,733 Contig Number | Contig Position | Variation Type | Variation Frequency (%) | Fold Coverage | Gene locus: SNP/Indel | AA change | Putative function |
|---|---|---|---|---|---|---|---|---|
| 84R1 | NZ_GG688464 | 439094 | Insertion | 97.66 | 128 | EFSG_RS08105: Insertion of A at 588 | Val197 Frameshift | Capsule EpsG family polymerase (*wzy*) |
| 84R2 | NZ_GG688461 | 441541 | SNP | 100 | 157 | EFSG_RS08120: 61C ⇒ T | Gln21 ⇒ Stop | Capsule nucleotide sugar dehydrogenase |
| 84R3 | NZ_GG688461 | 441541 | SNP | 100 | 138 | EFSG_RS08120: 61C ⇒ T | Gln21 ⇒ Stop | Capsule nucleotide sugar dehydrogenase |
| 84R4 | NZ_GG688461 | 441541 | SNP | 100 | 128 | EFSG_RS08120: 61C ⇒ T | Gln21 ⇒ Stop | Capsule nucleotide sugar dehydrogenase |
| 84R5 | NZ_GG688461 | 436239 | Deletion | 100 | 110 | EFSG_RS08090: Deletion of C at 641 | Ala214 Frameshift | Capsule Aminotransferase |

| | | | | | | | | |
|---|---|---|---|---|---|---|---|---|
| 84R6 | NZ_GG688461 | 744354 | SNP | 100 | 57 | EFSG_RS09545: 557C ⇒ T | Ala186 ⇒ Val | RDD family protein |
| 84R6 | NZ_GG688461 | 431883 | SNP | 98.92 | 93 | EFSG_RS08070: 250G ⇒ T | Asp84 ⇒ Tyr | Capsule tyrosine-protein kinase (*wze*) |
| 84R8 | No mutation detected | | | | | | | |

SNP – Single Nucleotide Polymorphism; Indel – Insertion/Deletion; AA – Amino Acid; RDD – Arginine Aspartate Aspartate;
