## Supplementary Table 3 for "Lytic bacteriophages facilitate antibiotic sensitization of *Enterococcus faecium*"

Table S3A. **Phage 9181 resistance enhances antimicrobial susceptibility by E-test**

| | | Ampicillin | | Ceftriaxone | | Daptomycin | |
|---|---|---|---|---|---|---|---|
| **Strain** | **Strain Mutation** | Mean MIC (µg/mL) | Standard deviation (µg/mL) | MIC (µg/mL) | Standard deviation (µg/mL) | MIC (µg/mL) | Standard deviation (µg/mL) |
| **Com12** | n/a | 1.33 | 0.29 | >32 | 0 | 1.83 | 0.29 |
| **81R3** | *sagA* | 0.56 | 0.41 | **2.67**$^{****}$ | **1.53** | 1.83 | 0.29 |
| **81R4** | *sagA* | **0.5**$^{*}$ | **0.25** | **2.67**$^{****}$ | **0.58** | 1.83 | 0.29 |
| **81R5** | *sagA* | 0.83 | 0.29 | >32 | 0 | 1.83 | 0.29 |
| **81R6** | *sagA* | **0.29**$^{**}$ | **0.18** | **0.83**$^{****}$ | **0.14** | 2.17 | 0.76 |
| **81R7** | *wze*, *topB* | 0.92 | 0.52 | >32 | 0 | 1.83 | 0.29 |
| **81R8** | *sagA* | **0.44**$^{*}$ | **0.28** | **1.33**$^{****}$ | **0.29** | 1.83 | 0.29 |

*$P$ < 0.05, $^{**}P$ < 0.01, $^{****}P$ <0.0001 by unpaired t-test; MIC, minimum inhibitory concentration; n/a, not applicable

Table S3B. **Phage 9183 resistance enhances antimicrobial susceptibility by E-test**

| | | Ampicillin | | Ceftriaxone | | Daptomycin | |
|---|---|---|---|---|---|---|---|
| **Strain** | Strain Mutation | MIC (µg/mL) | Standard deviation (µg/mL) | MIC (µg/mL) | Standard deviation (µg/mL) | MIC (µg/mL) | Standard deviation (µg/mL) |
| **733** | n/a | 1.08 | 0.38 | > 32 | 0 | 1.67 | 0.29 |
| **83R1** | *epaR* *nth* | **0.20*** | **0.17** | **0.29******** | **0.08** | **0.5**** | **0** |
| **83R3** | *epaR* | **0.36*** | **0.16** | **0.33******** | **0.14** | **0.42**** | **0.14** |
| **83R4** | *epaX* *gdh* | **0.34*** | **0.19** | **0.33******** | **0.14** | **0.33**** | **0.14** |
| **83R5** | *epaR* *gnsA* *sorC* | **0.27*** | **0.20** | **0.88******** | **0.98** | **0.50**** | **0.25** |
| **83R6** | *epaR* | **0.30*** | **0.10** | **0.25******** | **0** | **0.67*** | **0.29** |
| **83R7** | *epaX* | **0.31*** | **0.16** | **0.67******** | **0.29** | **0.38**** | **0.13** |
| **83R8** | *epaR* | **0.27*** | **0.20** | **0.54******** | **0.40** | **0.67*** | **0.29** |

*$P$ < 0.05, **$P$ < 0.01, ****$P$ <0.0001 by unpaired t-test; MIC, minimum inhibitory concentration; n/a, not applicable

Table S3C. **Phage 9184 resistance does not alter antimicrobial susceptibility by E-test**

| | | Ampicillin | | Ceftriaxone | | Daptomycin | |
|---|---|---|---|---|---|---|---|
| Strain | Strain Mutation | MIC (µg/mL) | Standard deviation (µg/mL) | MIC (µg/mL) | Standard deviation (µg/mL) | MIC (µg/mL ) | Standard deviation (µg/mL) |
| 733 | n/a | 1.25 | 0.43 | >32 | 0 | 1.83 | 0.29 |
| 84R1 | *efsg_rs08105* | 1.25 | 0.43 | >32 | 0 | 1.83 | 0.29 |
| 84R2 | *efsg_rs08120* | 1.33 | 0.76 | >32 | 0 | 1.83 | 0.29 |
| 84R3 | *efsg_rs08120* | 1.23 | 0.93 | >32 | 0 | 2 | 0 |
| 84R4 | *efsg_rs08120* | 1.17 | 0.58 | >32 | 0 | 1.67 | 0.29 |
| 84R5 | *efsg_rs08090* | 1.17 | 0.58 | >32 | 0 | 1.67 | 0.29 |
| 84R6 | *rdd*, *wze* | 1.33 | 0.76 | >32 | 0 | 2 | 0 |

MIC, minimum inhibitory concentration; n/a, not applicable
