## Supplementary Table 4 for "Lytic bacteriophages facilitate antibiotic sensitization of *Enterococcus faecium*"

| Table S4. Bacterial strains, phages, plasmids and primers | | |
|---|---|---|
| **Strains, phages, plasmids and primers** | **Characteristics and/or description** | **Reference/ Source** |
| ***Enterococcus faecium*** | | |
| **1,141,733** | clinical isolate (blood); USA, 2006 | (1) |
| **Com12** | human fecal isolate; USA, 2006 | (1) |
| **Com15** | Human fecal isolate; USA, 2006 | (1) |
| **1,231,408** | clinical isolate (blood); USA, 2005; $Amp^R$, $Cip^R$ | (1) |
| **1,231,501** | clinical isolate (blood); USA, 2005 | (1) |
| **1,231,410** | clinical isolate (blood); USA, 2005; $Van^R$, $Amp^R$ $Cip^R$ | (1) |
| **1,231,502** | clinical isolate (blood); USA, 2005; $Van^R$, $Amp^R$ $Cip^R$ | (1) |
| **1,230,933** | clinical isolate (blood); USA, 2005; $Van^R$, $Amp^R$ $Cip^R$ | (1) |
| **U37** | clinical isolate (tissue source unknown); USA; 1998; $Van^R$, $Amp^R$, $Erm^R$, $Gen^R$, $Str^R$, $Tet^R$ | (2) |
| **H11** | clinical isolate (tissue source unknown); USA; 1998; $Van^R$, $Amp^R$, $Erm^R$, $Gen^R$, $Str^R$, $Tc^R$ | (2) |
| **81R3-*sagA*** | 81R3 (*sagA* SNP) strain carrying pAM401-*sagA* complementation vector | This study |
| **81R3-E** | 81R3 (*sagA* SNP) strain carrying pAM401 empty vector | This study |
| **81R4-*sagA*** | 81R4 (*sagA* SNP) strain carrying pAM401-*sagA* complementation vector | This study |
| **81R4-E** | 81R4 (sagA SNP) strain carrying pAM401 empty vector | This study |
| **81R5-*sagA*** | 81R5 (*sagA* SNP) strain carrying pAM401-*sagA* complementation vector | This study |
| **81R5-E** | 81R5 (sagA SNP) strain carrying pAM401 empty vector | This study |
| **81R6-*sagA*** | 81R6 (*sagA* SNP) strain carrying pAM401-*sagA* complementation vector | This study |
| **81R6-E** | 81R6 (*sagA* SNP) strain carrying pAM401 empty vector | This study |
| **81R7-*wze*-Com12** | 81R7 (*wze*, *topB* SNPs) strain carrying pLZ12a-*wze*-Com12 complementation vector | This study |
| **81R7-E** | 81R7 (*wze*, *topB* SNPs) strain carrying pLZ12a empty vector | This study |
| **81R8-*sagA*** | 81R8 (*sagA* SNP) strain carrying pAM401-*sagA* complementation vector | This study |

| | | |
|---|---|---|
| **81R8-E** | 81R8 (*sagA* SNP) strain carrying pAM401 empty vector | This study |
| **83R1-*epaR*** | 83R1 (*epaR*, *nth* SNPs) strain carrying pLZ12a-*epaR* complementation vector | This study |
| **83R1-E** | 83R1 (*epaR*, *nth* SNPs) strain carrying pLZ12a empty vector | This study |
| **83R3-*epaR*** | 83R1 (*epaR* SNP) strain carrying pLZ12a-*epaR* complementation vector | This study |
| **83R3-E** | 83R3 (*epaR* SNP) strain carrying pLZ12a empty vector | This study |
| **83R4-*epaX*** | 83R4 (*epaX, gdh* SNP) strain carrying-*epaX* complementation vector | This study |
| **83R4-E** | 83R4 (*epaX, gdh* SNP) strain carrying pLZ12a empty vector | This study |
| **83R5-*epaR*** | 83R5 (*epaR, gnsA*, *sorC* SNPs) strain carrying pLZ12a-*epaR* complementation vector | This study |
| **83R5-E** | 83R5 (*epaR*, *gnsA*, *sorC* SNPs) strain carrying pLZ12a empty vector | This study |
| **83R6-*epaR*** | 83R6 (*epaR* SNP) strain carrying pLZ12a-*epaR* complementation vector | This study |
| **83R6-E** | 83R6 (*epaR* SNP) strain carrying pLZ12a empty vector | This study |
| **83R7-*epaX*** | 83R7 (*epaX*, *dltA* SNPs) strain carrying pLZ12a-*epaX* complementation vector | This study |
| **83R7-*dlta*** | 83R7 (*epaX*, *dltA* SNPs) strain carrying pLZ12a-*dltA* complementation vector | This study |
| **83R7-E** | 83R7 (*epaX*, *dltA* SNPs) strain carrying pLZ12a empty vector | This study |
| **83R8-*epaR*** | 83R8 (*epaR* SNP) strain carrying pLZ12a-*epaR* complementation vector | This study |
| **83R8-E** | 83R8 (*epaR* SNP) strain carrying pLZ12a empty vector | This study |
| **84R2-8120** | 84R2 (*efsg_rs08120* SNP) strain carrying pLZ12a-8120 vector | This study |
| **84R2-E** | 84R2 (*efsg_rs08120* SNP) strain carrying pLZ12a empty vector | This study |
| **84R3-8120** | 84R3 (*efsg_rs08120* SP) strain carrying pLZ12a-8120 vector | This study |
| **84R3-E** | 84R3 (*efsg_rs08120* SNP) strain carrying pLZ12a empty vector | This study |
| **84R4-8120** | 84R4 (*efsg_rs08120* SP) strain carrying pLZ12a-8120 vector | This study |
| **84R4-E** | 84R4 (*efsg_rs08120* SNP) strain carrying pLZ12a empty vector | This study |
| **84R5-8090** | 84R5 (*efsg_rs08105* SNP) strain carrying pLZ12a-8090 complementation vector | This study |
| **84R5-E** | 84R5 *(efsg_rs08105* SNP) strain carrying pLZ12a empty vector | This study |
| **84R6-*wze*-733** | 84R6 (*wze*, *rdd* SNP) strain carrying pLZ12a-*wze* complementation vector | This study |
| **84R6-E** | 84R6 (*wze*, *rdd* SNP) strain carrying pLZ12a empty vector | This study |

| | | |
|---|---|---|
| **UCH1** | clinical isolate (blood); $Dap^{R}$, $Amp^{R}$, $Van^{R}$; University of Colorado Hospital | This study |
| **UCH2** | clinical isolate (blood); University of Colorado Hospital | This study |
| **UCH3** | clinical isolate (blood); $Dap^{SDD}$, $Amp^{R}$, $Van^{R}$; University of Colorado Hospital | This study |
| **UCH4** | clinical isolate (blood); $Dap^{R}$, $Amp^{R}$, $Van^{R}$; University of Colorado Hospital | This study |
| **UCH5** | clinical isolate (blood); $Dap^{R}$, $Amp^{R}$, $Van^{R}$; University of Colorado Hospital | This study |
| **UCH6** | clinical isolate (blood); $Dap^{R}$, $Lin^{I}$, $Amp^{R}$, $Van^{R}$; University of Colorado Hospital | This study |
| **UCH7** | clinical isolate (blood); $Dap^{SDD}$, $Amp^{R}$, $Van^{R}$; University of Colorado Hospital | This study |
| **UCH8** | clinical isolate (blood); $Amp^{R}$, $Van^{R}$, $Str^{R}$; University of Colorado Hospital | This study |
| **UCH9** | clinical isolate (blood); $Amp^{R}$, $Van^{R}$; University of Colorado Hospital | This study |
| **UCH10** | clinical isolate (spleen); $Dap^{R}$, $Amp^{R}$; University of Colorado Hospital | This study |
| **UCH11** | clinical isolate (ascites); University of Colorado Hospital | This study |

| | | |
|---|---|---|
| **Phage resistant strains obtained *in vitro*** | | |
| **81R3** | phage 9181 resistant mutant of Com12; *sagA* SNP; Trp 433 Leu | This study |
| **81R4** | phage 9181 resistant mutant of Com12; *sagA* SNP; Trp 433 Cys | This study |
| **81R5** | phage 9181 resistant mutant of Com12; *sagA* SNP; Gly 460 Asp | This study |
| **81R6** | phage 9181 resistant mutant of Com12; *sagA* insertion; Leu insertion between Tyr 451 and Leu 452 | This study |
| **81R7** | phage 9181 resistant mutant of Com12; *wze* SNP; Pro 167 Ser; *topB* SNP; Gly 109 Trp | This study |
| **81R8** | phage 9181 resistant mutant of Com12; *sagA* SNP; Gly 435 Val | This study |
| **83R1** | phage 9183 resistant mutant of 1,141,733; *epaR* SNP; Arg 338 Cys; *nth* SNP; Ala 10 Ser | This study |
| **83R2** | Phage 9183 resistant mutat of 1,141,733; unknown mutation causing phage 9183 resistance | This study |
| **83R3** | phage 9183 resistant mutant of 1,141,733; *epaR* deletion; Phe 103 frameshift | This study |
| **83R4** | phage 9183 resistant mutant of 1,141,733; *epaX* deletion; Asn 66 frameshift; *gdh* SNP; Trp 104 Cys | This study |

| | | |
|---|---|---|
| **83R5** | phage 9183 resistant mutant of 1,141,733; *epaR* deletion; Glu 211 frameshift; *gnsA* SNP; Glu 2 Gly; *sorC* SNP; Leu 329 Phe | This study |
| **83R6** | phage 9183 resistant mutant of 1,141,733; *epaR* SNP; Met 310 Ile | This study |
| **83R7** | phage 9183 resistant mutant of 1,141,733; *epaX* deletion; Asn 66 frameshift; *dltA* SNP; Glu 8 Stop | This study |
| **83R8** | phage 9183 resistant mutant of 1,141,733; *epaR* SNP; Glu 315 Gly | This study |
| **84R2** | phage 9184 resistant mutant of 1,141,733; *efsg_rs08120* SNP; Gln 21 Stop | This study |
| **84R3** | phage 9184 resistant mutant of 1,141,733; *efsg_rs08120* SNP; Gln 21 Stop | This study |
| **84R4** | phage 9184 resistant mutant of 1,141,733; *efsg_rs08120* SNP; Gln 21 Stop | This study |
| **84R5** | phage 9184 resistant mutant of 1,141,733; *efsg_rs08090* deletion; Ala 214 frameshift | This study |
| **84R6** | phage 9184 resistant mutant of 1,141,733; *wze* SNP; Asp 84 Tyr; *rdd* SNP; Ala 186 Val | This study |
| **84R8** | Phage 9184 resistant mutant of 1,141,733; unknown mutation causing phage 9183 resistance | This study |
| ***Enterococcus faecalis*** | | |
| **OG1RF** | Human oral isolate; $Rf^R$, $Fa^R$ | (3) |
| **UCH12** | clinical isolate (blood); $Str^R$, $Gen^R$; University of Colorado Hospital | This study |
| **UCH13** | clinical isolate (blood); $Str^R$, $Gen^R$; University of Colorado Hospital | This study |
| **UCH14** | clinical isolate (blood); $Str^R$; University of Colorado Hospital | This study |
| **UCH15** | clinical isolate (blood); University of Colorado Hospital | This study |
| **UCH16** | clinical isolate (spine tissue); $Dox^R$; University of Colorado Hospital | This study |
| **UCH17** | clinical isolate (joint tissue); $Dox^R$; University of Colorado Hospital | This study |
| **UCH18** | clinical isolate (heart valve tissue); University of Colorado Hospital | This study |
| **UCH19** | clinical isolate (heart valve tissue); University of Colorado Hospital | This study |
| **UCH20** | clinical isolate (heart valve tissue); University of Colorado Hospital | This study |
| ***Escherichia coli*** | | |

| | | |
|---|---|---|
| **TG1** | *[F' traD36 proAB lacIqZ ΔM15] supE thi-1 Δ(lac-proAB) Δ(mcrBhsdSM)5(rK - mK -)* | Lucigen |
| **Phages** | | |
| **phage 9181** | raw sewage isolate, prolate-head, Siphoviridae | This study |
| **phage 9183** | raw sewage isolate, icosahedral-head, Siphoviridae | This study |
| **phage 9184** | raw sewage isolate, icosahedral-head, Siphoviridae | This study |
| **Plasmids** | | |
| **pAM401** | *E. coli*-*E. faecalis* shuttle vector; pIP501 origin; $Cm^R$, $Tc^R$ | (4) |
| **pAM401-SagA** | pAM401 plasmid expressing *E. faecium* Com15 *sagA* promoter fused to *sagA* ORF with His-6 tag | (5) |
| **pLZ12A** | *bacA* promoter cloned into shuttle vector pLZ12; pSH71 origin; $Cm^R$ | (6, 7) |
| **pLZ12A-*wze*-Com12** | pLZ12A plasmid expressing *E. faecium* Com12 *wze* from the P-*bacA* promoter | This study |
| **pLZ12A-*epaR*** | pLZ12A plasmid expressing *E. faecium* 1,141,733 *epaR* from the *bacA* promoter | This study |
| **pLZ12A-*epaX*** | pLZ12A plasmid expressing *E. faecium* 1,141,733 *epaX* from the *bacA* promoter | This study |
| **pLZ12A-*dltA*** | pLZ12A plasmid expressing *E. faecium* 1,141,733 *dltA* from the *bacA* promoter | This study |
| **pLZ12A-8120** | pLZ12A plasmid expressing *E. faecium* 1,141,733 *efsg_rs08120* from the *bacA* promoter | This study |
| **pLZ12A-8090** | pLZ12A plasmid expressing *E. faecium* 1,141,733 *efsg_rs08090* from the *bacA* promoter | This study |
| **pLZ12A-*wze*-733** | pLZ12A plasmid expressing *E. faecium* 1,141,733 *wze* from the *bacA* promoter | This study |
| **Primers** | | |
| ***wze*-Com12-comp-F** | NNNNNNGAATTCATGGCACGAACACAGAAACA | This Study |

| | | |
|---|---|---|
| ***wze*-Com12-comp-R** | NNNNNNGGATCCTCGGTGGATGTCTTCGATCA | This Study |
| ***epaR*-comp-F** | NNNNNNCTGCAGATGAATAAAAATGGGGAGTGGAATG | This Study |
| ***epaR*-comp-R** | NNNNNNGGATCCCTCCTTGGATAGCTGACTGAATC | This Study |
| ***epaX*-comp-F** | NNNNNNGAATTCATGTGTGAGATTAGTATTATTGTTCCTG | This Study |
| ***epaX*-comp-R** | NNNNNNGGATCCTGAAATGGTCCTCCCTACCT | This Study |
| ***dltA*-comp-F** | NNNNNNGAATTCATGGAAATCAAAACGATTATTGAAGC | This Study |
| ***dlta*-comp-R** | NNNNNGGATCCAACGATTGGTATAAGCGCAATG | This Study |
| **8120-comp-F** | NNNNNNCTGCAGATGAAAGTATCAGTTTTTGGTCTC | This Study |
| **8120-comp-R** | NNNNNNGGATCCTTCATGGTTTAATCCCGTCTAA | This Study |
| **8090-comp-F** | NNNNNNGAATTCTTGGAAAATAAACGAATATTATTAGCATCT | This Study |
| **8090-comp-R** | NNNNNNGGATCCTGAGAGCGATACTTGACAATAGG | This Study |
| ***wze*-733-comp-F** | NNNNNNGAATTCATGGCACGAACACAGAAACA | This Study |
| ***wze*-733-comp-R** | NNNNNNGGATCCCGATCATTTCTCTTGTCTCCTCTC | This Study |
| **16270-Com12-gap-F** | NNNNNNCAGAACAAGCACGTCAACAAG | This Study |
| **16270-com12-gap-R** | NNNNNNAATCCTGAGCAGTCAAATCCA | This Study |
| **Phage-9181-Lysin-F** | GCAACGCATAACCAACCTAAC | This Study |
| **Phage-9181-Lysin-R** | GTCTCCACCTTGATAGCCATAC | This Study |
| **Phage-9183-Integrase-F** | GCAGACATTCGTGCTTTCTTT | This Study |
| **Phage-9183-Integrase-R** | CTCCTCGTTGATCAAACCATTTC | This Study |
| **Phage-9184-Lysin-F** | GGGTAACTCAACAGCCATACA | This Study |
| **Phage-9184-Lysin-R** | AGTTCTTGTCCGCCTTGATAG | This Study |

**Dap$^R$ - daptomycin resistance; Dap$^{SDD}$ - daptomycin susceptibility dose dependent; Lin$^I$ - linezolid intermediate; Amp$^R$ - ampicillin resistant; Van$^R$ - vancomycin resistant; Str$^R$ - Streptomycin resistant; Erm$^R$ – erythromycin resistant; Gen$^R$ – gentamicin resistant; Cip$^R$ – ciprofloxacin resistant; Dox$^R$ – doxycycline resistant; Rf$^R$ – rifampin resistant; Fa$^R$ – fusidic acid resistant; Restriction sites are underlined**

**References:**

1. Palmer KL, Godfrey P, Griggs A, Kos VN, Zucker J, Desjardins C, Cerqueira G, Gevers D, Walker S, Wortman J, Feldgarden M, Haas B, Birren B, Gilmore MS. 2012. Comparative genomics of enterococci: variation in *Enterococcus faecalis*, clade structure in *E. faecium*, and defining characteristics of *E. gallinarum* and *E. casseliflavus*. mBio 3:e00318-11. doi:10.1128/mBio.00318-11.

2. Rice LB, Carias LL, Donskey CL, Rudin SD. 1998. Transferable, plasmid-mediated VanB-type glycopeptide resistance in *Enterococcus faecium*. Antimicrob Agents Chemother 42:963-4.

3. Bourgogne A, Garsin DA, Qin X, Singh KV, Sillanpaa J, Yerrapragada S, Ding Y, Dugan-Rocha S, Buhay C, Shen H, Chen G, Williams G, Muzny D, Maadani A, Fox KA, Gioia J, Chen L, Shang Y, Arias CA, Nallapareddy SR, Zhao M, Prakash VP, Chowdhury S, Jiang H, Gibbs RA, Murray BE, Highlander SK, Weinstock GM. 2008. Large scale variation in *Enterococcus faecalis* illustrated by the genome analysis of strain OG1RF. Genome Biol 9:R110. doi:10.1186/gb-2008-9-7-r110.

4. Wirth R, An FY, Clewell DB. 1986. Highly efficient protoplast transformation system for *Streptococcus faecalis* and a new *Escherichia coli*-*S. faecalis* shuttle vector. J Bacteriol 165:831-6. doi:10.1128/jb.165.3.831-836.1986.

5. Rangan KJ, Pedicord VA, Wang YC, Kim B, Lu Y, Shaham S, Mucida D, Hang HC. 2016. A secreted bacterial peptidoglycan hydrolase enhances tolerance to enteric pathogens. Science 353:1434-1437. doi:10.1126/science.aaf3552.

6. Chatterjee A, Johnson CN, Luong P, Hullahalli K, McBride SW, Schubert AM, Palmer KL, Carlson PE, Jr., Duerkop BA. 2019. Bacteriophage resistance alters antibiotic-mediated intestinal expansion of enterococci. Infect Immun 87:e00085-19. doi:10.1128/iai.00085-19.

7. Perez-Casal J, Caparon MG, Scott JR. 1991. Mry, a trans-acting positive regulator of the M protein gene of *Streptococcus pyogenes* with similarity to the receptor proteins of two-component regulatory systems. J Bacteriol 173:2617-24. doi:10.1128/jb.173.8.2617-2624.1991.
